# BCL-2 and BOK regulate apoptosis by interaction of their C-terminal transmembrane domains

**DOI:** 10.1101/2024.05.13.593225

**Authors:** Tobias B. Beigl, Alexander Paul, Thomas Fellmeth, Dang Nguyen, Lynn Barber, Sandra Weller, Benjamin Schäfer, Bernhard F. Gillissen, Walter E. Aulitzky, Hans-Georg Kopp, Markus Rehm, David W. Andrews, Kristyna Pluhackova, Frank Essmann

## Abstract

The Bcl-2 family controls apoptosis by direct interactions of pro- and anti-apoptotic proteins. The principle mechanism is binding of the BH3 domain of pro-apoptotic proteins to the hydrophobic groove of anti-apoptotic siblings, which is therapeutically exploited by approved BH3-mimetic anti-cancer drugs. Evidence suggests that also the transmembrane domain (TMD) of Bcl-2 proteins affects Bcl-2 interactions. We developed a highly-specific split luciferase assay, enabling the analysis of TMD interactions of pore-forming apoptosis effectors BAX, BAK, and BOK with anti-apoptotic Bcl-2 proteins in living cells. We confirm homotypic interaction of the BAX-TMD, but also newly identify interaction of the TMD of anti-apoptotic BCL-2 with the TMD of BOK, a so far very peculiar pro-apoptotic Bcl-2 protein. Interaction of BOK-TMD with BCL-2-TMD localizes at the endoplasmic reticulum (ER). Molecular dynamics simulations in an ER membrane model confirm dynamic BOK-TMD and BCL-2-TMD homo- and heterodimers and stable heterotetramers. Inhibition of BOK-induced apoptosis by BCL-2 depends specifically on their TMDs. Thus, TMDs of Bcl-2 proteins are a relevant interaction interface for apoptosis regulation and provide a novel potential drug target.

## Introduction

Proteins of the Bcl-2 family constitute the central hub for intracellular regulation of cell survival and demise by either preventing or promoting apoptotic cell death. Classically, the Bcl-2 protein family is divided into three subgroups according to their structure and function in apoptosis signaling (Figure S1A, comprehensively reviewed in (Kale et al. 2018b)): Pro-survival/anti-apoptotic Bcl-2 family proteins (i) (including BCL-2, BCL-XL, BCL-W, MCL-1 and BFL-1/A1) interact with and inhibit pro-apoptotic siblings by accommodating their BH3 domain, one of the four different Bcl-2 homology (BH) domains in a hydrophobic groove of the anti-apoptotic counterparts which results in mutual sequestration. Pro-apoptotic multidomain effector proteins (ii) BAX, BAK and BOK, once activated, oligomerize and initialize cell death by forming pores in the mitochondrial outer membrane (MOM). The third group, the so-called BH3-only proteins (iii), can be split into two subgroups according to their proposed mode of action: Sensitizer BH3-only proteins like BAD and NOXA act by occupying the hydrophobic groove of anti-apoptotic Bcl-2 proteins thereby blocking their capacity to hold active effector proteins in check. Activator BH3-only proteins like BID and BIM, however, allegedly directly interact with the pore-forming effectors and induce their conversion into an active conformation. Since active effectors can be blocked by anti-apoptotic Bcl-2 proteins, cells are ultimately sentenced to death once the anti-apoptotic capacity is exhausted and oligomerization of effector proteins mediates MOM permeabilization (MOMP). Upon MOMP, pro-apoptotic factors are released from mitochondria, importantly cytochrome c, which instigates activation of apoptosis-specific proteases (caspases) and destruction of the cell.

The BH3 domain and hydrophobic groove are indisputably recognized as important interaction sites of Bcl-2 proteins. Hence, the individual amino acid sequence of the specific BH3 domains in pro-apoptotic BH3-only and effector proteins and the amino acid composition of the hydrophobic groove in anti-apoptotic proteins cause variations in their mutual affinity and thus constitute the basis of the complex interaction network of Bcl-2 proteins (Banjara et al. 2020; Osterlund et al. 2022). Importantly, elucidation of the structural basis for Bcl-2 protein interaction inspired the development of small-molecule drugs that mimic pro-apoptotic BH3 domains to specifically bind to the hydrophobic groove of anti-apoptotic Bcl-2 proteins and block their activity. These BH3 mimetics effectively exploit the Bcl-2 interaction network to push cancer cells over the threshold to apoptosis. The orally available BH3 mimetic ABT-199/Venetoclax that specifically blocks BCL-2 is effective in the treatment of BCL-2-dependent hematopoietic malignancies such as multiple myeloma (MM), acute myeloid leukemia (AML) and chronic lymphoid leukemia (CLL) (Roberts et al. 2016; Deeks 2016; DiNardo et al. 2019).

Successively increasing detailed elucidation of Bcl-2 protein interaction instigated refinement of models of the Bcl-2 regulatory network from the initial “rheostat” (Korsmeyer et al. 1993) to “direct activation” (Letai et al. 2002), “embedded together” (Leber et al. 2007), “unified” (Llambi et al. 2011), and “hierarchical” (Chen et al. 2015) model. In addition to mutual binding via BH3 domain and hydrophobic groove, the dynamic Bcl-2 interaction is modulated by protein abundance, post-translational modification, and subcellular localization (Kale et al. 2018b). The most important site where Bcl-2 proteins exert their function, i.e. regulation of cytochrome c release, are mitochondria. Consequently, composition of the MOM proposedly affects membrane interaction with Bcl-2 proteins and membrane insertion. Interestingly, the mitochondria-specific membrane component cardiolipin has even been proposed to “glue” together BAX transmembrane domains during oligomerization (Lai et al. 2019). A C-terminal transmembrane domain (TMD) is present in most Bcl-2 proteins which was early recognized to target the family members to specific intracellular membranes (Zhu et al. 1996; Horie et al. 2002). Although generally referred to as TMD, the individual Bcl-2 C-termini insert into the lipid bilayer to varying extent. In general, TMDs consist of roughly 20 amino acids that form a single-pass α-helix flanked by charged amino acids on either side (Schinzel et al. 2004). In contrast, tail-anchor sequences tend to have fewer contiguous hydrophobic residues (∼15) followed by less than 12 more hydrophilic residues (Brito et al. 2019). Naturally, amino acid sequence of the TMD crucially affects subcellular localization of Bcl-2 proteins. For example, specific targeting of BAK or BCL-XL to MOM depends on flanking basic amino acid side chains as well as on hydrophobicity (Kaufmann et al. 2003). The TMD of BCL-2 is more hydrophobic in the N-terminal half and not flanked by basic amino acids on both sides, resulting in BCL-2 localization at various subcellular membranes including MOM, endoplasmic reticulum (ER) and the nuclear envelope (Schinzel et al. 2004; Kale et al. 2018b). TMDs were shown to be a critical structural feature in Bcl-2 family proteins in several studies (Jeong et al. 2004; Guedes et al. 2013; Stehle et al. 2018; Chi et al. 2020) since absence, mutation or post-translational modification of TMDs substantially affects targeting and function of Bcl-2 proteins (Nechushtan et al. 1999; Gardai et al. 2004; Simonyan et al. 2016; Kale et al. 2018a; Lucendo et al. 2020).

In addition to subcellular localization increasing evidence indicates that direct TMD-TMD interaction of Bcl-2 proteins critically impacts on apoptosis regulation, e.g. enlargement of BAX-oligomeric pores depends on BAX dimer-dimer interactions via their TMDs (Zhang et al. 2016). Also, interaction of pro- and anti-apoptotic Bcl-2 proteins such as MCL-1 and BOK via TMDs modulates cell death regulation (Andreu-Fernández et al. 2017; Lucendo et al. 2020) stressing fine-tuning of the BH3 domain:hydrophobic groove-based interaction scheme. The BH3-only proteins BIML and PUMA bind to their anti-apoptotic counterparts BCL-2 and BCL-XL by both BH3 domain and TMD which substantially influences binding affinity and is described as a crucial “double-bolt lock” mechanism (Liu et al. 2019; Pemberton et al. 2023).

The first identified member and namesake of the family, the anti-apoptotic protein BCL-2, efficiently inhibits BAX and BAK pore formation in the MOM. However, in non-apoptotic cells a large fraction of BCL-2 resides at the ER, which mainly depends on the BCL-2 TMD (Kaufmann et al. 2003). Recently, TOM20-mediated translocation of BCL-2 from ER to mitochondria-associated membranes (MAMs) and mitochondria upon apoptosis induction was postulated by Lalier et al. (Lalier et al. 2021). Andreu-Fernández et al. showed, that BCL-2 TMD peptides not only form stable homodimers but also interact with BAX-TMD in biological membranes (Andreu-Fernández et al. 2017). Survival promotion by BCL-2 takes place also at the ER since BCL-2 regulates calcium signaling by binding to inositol 1,4,5-trisphosphate receptors (IP3Rs) in ER membranes (Rong et al. 2009; Monaco et al. 2012; Chang et al. 2014).

In contrast to BAX and BAK, the pro-apoptotic protein BOK is localized at the ER (Echeverry et al. 2013). Also BOK possesses a C-terminal membrane-localizing sequence, termed as TMD, although no empirical data confirmed its membrane-spanning character. Early after identification of BOK a yeast two-hybrid assay suggested interaction of BOK with BFL1/ A1 and MCL-1 (Hsu et al. 1997). The interaction site of BOK with anti-apoptotic MCL-1 was recently pinpointed to their TMDs which is to date the only described direct interaction of BOK with anti-apoptotic Bcl-2 proteins (Stehle et al. 2018; Lucendo et al. 2020). However, BOK-induced apoptosis was formerly shown not to be counteracted by any anti-apoptotic Bcl-2 family protein (Llambi et al. 2016). Thus, despite sporadic progress in illuminating Bcl-2 TMD interaction, the bulk of the TMD network and its fine tuning of apoptosis signaling remains in the dark, overshadowed by the dominant BH3 domain:hydrophobic groove interaction.

Interaction of Bcl-2 TMDs was studied by various molecular biological methods, (Zhang et al. 2016; Andreu-Fernández et al. 2017; Lucendo et al. 2020) and also in-silico modelling (Wassenaar et al. 2015), was recently used to study the homomeric and heteromeric interactions in transmembrane dimers of MCL-1, BAK and BAX in an artificial phosphocholine membrane (Lucendo et al. 2020).

Here, employing a novel split-luciferase assay, we reveal direct interaction of BCL-2-TMD with TMD of pro-apoptotic effector protein BOK. Fluorophore-conjugated TMD peptides and full-length proteins co-localize at the ER. This TMD-directed co-localization is independent of the BH3 domain. High-throughput multiscale MD simulation of BCL-2-TMD/BOK-TMD oligomerization in an ER membrane mimic supports interaction of BOK-TMD and BCL-2-TMD. Simulations propose dynamic heterodimers and -trimers, but unexpectedly also stable tetramers of two BOK-TMDs and two BCL-2-TMDs, suggesting that BOK is retained in the ER membrane by BCL-2. Functionally, BCL-2 inhibits BOK-induced apoptosis and consequently TMD amino acid sequence in BOK critically affects inhibition by BCL-2. The newly identified interaction of the TMDs of BOK and BCL-2 is a new component of the regulatory interaction network of Bcl-2 family proteins representing BH3 domain-independent regulation of apoptosis.

## Results

### BCL-2-TMD and BOK-TMD interaction revealed by a novel bimolecular split luciferase assay

Several studies proposed interaction of selected Bcl-2 proteins via their TMDs, e.g. BAX and BCL-XL (Todt et al. 2013; Andreu-Fernández et al. 2017; Lucendo et al. 2020). Since these reports show a mere fraction of TMD interaction among Bcl-2 proteins, we initially set out to systematically analyze interaction of the TMDs in anti-apoptotic and pro-apoptotic effector Bcl-2 proteins. We designed a bimolecular split luciferase assay based on the NanoBiT system which has the advantage of a) very low background signal because the subunits of NanoBiT, large BiT (LgBiT) and small BiT (SmBiT), have low affinity and b) can be performed in living cells (England et al. 2016). The generated plasmids encode for LgBiT or SmBiT - fused by a hydrophilic linker sequence to the TMD of interest (Figure 1A, B). Upon expression, interaction of a respective pair of TMDs brings LgBiT and SmBiT in close proximity and allows for formation of the functional luciferase which then can process the substrate coelenterazine H to produce a luminescence signal. Additionally, plasmids encode for preceding fluorophores mCitrine (LgBiT-TMD) or mTurquoise2 (SmBiT-TMD), respectively, separated by a T2A self-cleaving sequence from the NanoBiT-TMD to allow normalization of the luminescence signal to the fluorescence of mCitrine and/or mTurquoise2 (Figure 1B).

**Figure 1:**
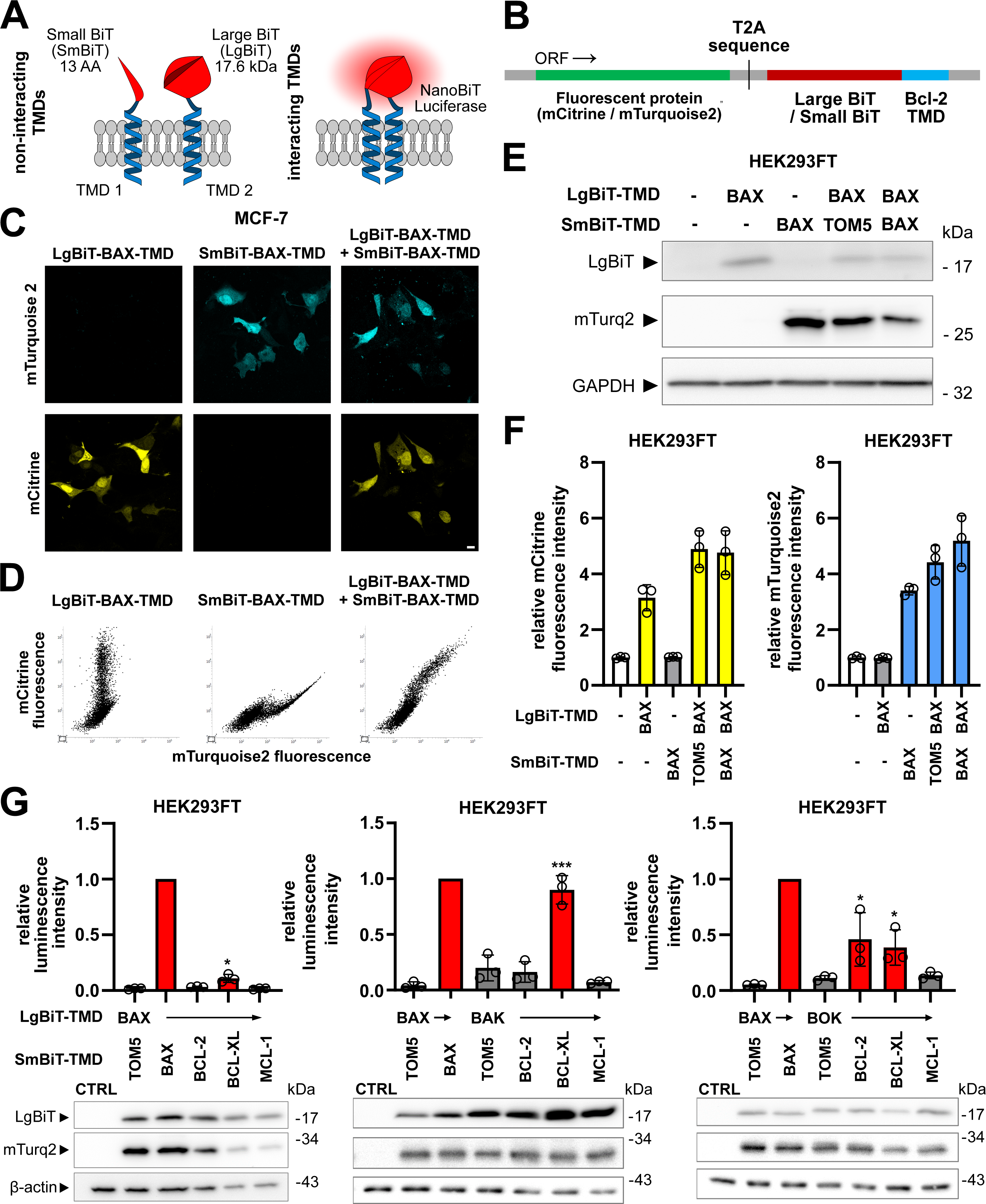
A bimolecular split luciferase assay reveals that TMDs of BCL-2 interact with TMDs of BOK but not with TMDs of BAX or BAK. **A** - Schematic of developed split-luciferase assay and its working principle. **B** - Schematic of plasmid insert structure used for NanoBiT-TMD fusion protein expression featuring simultaneous expression of mCitrine (LgBiT-TMDs) or mTurquoise2 (SmBiT-TMDs). **C** - Specific expression of mCitrine and mTurquoise2 from NanoBiT-TMD plasmids. MCF-7 cells were transiently transfected with plasmids for the expression of LgBiT-BAX-TMD, SmBiT-BAX-TMD or a combination of both. Cells were fixed after 18 h followed by cLSM. Images validate specific detection of mTurquoise2 and mCitrine. Representative images out of three independent experiments. Scale bar = 10 µm. **D** - Proportional co-expression of mCitrine and mTurquoise2. Scatter plots show mCitrine/mTurquoise2 fluorescence intensity detected in single HEK293FT cells transfected as in C and analyzed after 18 h using flow cytometry. Representative graphs out of three independent experiments. **E** - Western Blot verifies NanoBiT-TMD expression. Whole cell lysates of HEK293FT cells transfected with plasmids for the expression of indicated NanoBiT-TMDs (CTRL is untransfected) and harvested 18 h post-transfection were analyzed for expression of LgBiT and mTurquoise2 by Western blot. Representative blot from two independent experiments. GAPDH was used as loading control. **F** - Bar graphs show specific fluorescence of mCitrine and mTurquoise2 during split-luciferase assay. mCitrine and mTurquoise2 fluorescence intensity of cells transfected as in E detected by a multimode plate reader. Intensities are shown relative to control as mean ± sd of three independent experiments. **G** - TMDs of all effector proteins interact with BCL-XL-TMD, while only BOK-TMD interacts with BCL-2-TMD.Split-luciferase assay in HEK293FT cells transfected with plasmids for the expression of indicated NanoBiT-TMDs. Cells were harvested after 24 h and samples were both used for split-luciferase assay and Western blot. BAX-TMD/TOM5-TMD serves as a negative control, while BAX-TMD/BAX-TMD serves as positive control. SmBiT-TMDs of anti-apoptotic Bcl-2 proteins (BCL-2/BCL-XL/MCL-1) were combined with LgBiT-BAX-TMD (left), LgBiT-BAK-TMD (middle) or LgBIT-BOK-TMD (right). Graphs show luminescence intensity relative to the positive control and normalized to mTurquoise2 fluorescence intensity. Shown is the mean ± sd from three independent experiments. In blots below, corresponding whole cell lysates were analysed for expression of LgBiT and mTurquoise2 (mTurq2). Representative blots from three independent experiments are shown.

To verify functionality of the split-luciferase assay, we transfected HEK293FT cells with combinations of LgBiT-BAX-TMD and either SmBiT-BAX-TMD or SmBiT-TOM5-TMD and analyzed luminescence, fluorescence and protein expression. Since BAX pores enlarge by BAX-TMD dimerization (Zhang et al. 2016), the BAX-TMD homotypic interaction served as a positive control. The TMD of TOM5, a component of the translocase of the outer membrane (TOM) mitochondrial import complex, served as a negative control non-binding partner for the BAX-TMD. Indeed, luminescence detected from co-expression of SmBiT-TOM5-TMD with LgBiT-BAX-TMD was similar to the background signal (background 185.9 RLU vs. 303.6 RLU). The co-expression of SmBiT-BAX-TMD and LgBiT-BAX-TMD resulted in a strong luminescence signal (16572.0 RLU) in line with homotypic interaction (Figure S1A, B). Simultaneous expression of fluorescent proteins was confirmed by confocal laser-scanning microscopy (cLSM) and flow cytometry (Figure 1C, D) showing a proportional signal of mCitrine and mTurquoise2 fluorescence in cells co-transfected with equal amounts of LgBiT-BAX-TMD (mCitrine) and SmBiT-BAX-TMD or SmBiT-TOM5-TMD (mTurquoise2). In an additional control experiment, titration of SmBiT:LgBIT ratio resulted in a peak luminescence signal at a ratio of 1:1 suggesting efficient homotypic dimerization of BAX-TMDs (Figure S1C). Moreover, Western blot confirmed expression of LgBiT-fused TMDs and expression of mTurquoise2 as a surrogate marker for expression of SmBiT-TMD (Figure 1E). Next, we simultaneously analyzed fluorescence and luminescence in a multimode plate reader. Specific mCitrine or mTurquoise2 fluorescence was detected in HEK293 cells transfected with respective plasmids (Figure 1F). Since the signal for mCitrine and mTurquoise2 was comparable in 1:1 co-transfected cells (Figure 1C, D, F; Figure S1), in subsequent experiments luminescence is normalized to fluorescence of mTurquoise2.

After successful validation of the split-luciferase system, we next analyzed interaction of TMDs from Bcl-2 effector proteins with TMDs of anti-apoptotic proteins BCL-2, BCL-XL and MCL-1 in HEK293FT cells. In these co-transfection experiments, we set the normalized luminescence of BAX-TMD homotypic interaction to 1. Combining LgBiT-TMD of effectors BAX, BAK or BOK with SmBiT-TMD of anti-apoptotic BCL-2, BCL-XL and MCL-1 we found relative normalized luciferase activity of BCL-2-TMD with BAX-TMD (0.03) or BAK-TMD (0.16) (Figure 1G) which were comparable to the negative control (BAX-/TOM5-TMD) that amounted to 0.05 relative normalized luminescence. Significant luminescence was detected for the combination of BCL-2-TMD with BOK-TMD reaching 46% of the positive control (BAX-/BAX-TMD) luminescence (Figure 1G). The interaction of BOK-TMD with BCL 2-TMD was confirmed in MCF-7 cells with a relative normalized luminescence of 30% compared to the positive control (Figure S1E). In addition, all effector TMDs produced a luminescence signal when co-expressed with BCL-XL-TMD indicating promiscuous interaction (Figure 1G).

Taken together, we developed and validated a plasmid-based, bimolecular split luciferase assay to analyze interaction of TMDs from Bcl-2 family proteins and revealed direct interaction of the BCL-2-TMD with BOK-TMD.

### BCL-2-TMD and BOK-TMD co-localize at the ER

As co-localization is a prerequisite for interaction, we next investigated the subcellular localization of TMD fused to fluorescent proteins. We assumed that the exposed TMD sequence in Bcl-2 proteins is sufficient to target the protein to a specific subcellular membrane, e.g. ER-like localization for BCL-2-TMD fused with enhanced green fluorescent protein (EGFP) (Egan et al. 1999; Kaufmann et al. 2003). Co-localization of fluorophore-coupled TMD peptides with fluorescent organelle markers then allows estimation of the TMD’s (and protein’s) subcellular localization.

MCF-7 cells that express subcellular markers for ER or mitochondria (see materials&methods) were transfected with plasmids for the expression of mTurquoise2 with C-terminal fused TMD. The subcellular localization of the TMD peptides was then imaged by cLSM. BAX-TMD and BAK-TMD were predominantly localized to the mitochondria, while BOK-TMD and BCL-2-TMD both co-localized with the ER marker (Figure 2 A, B). BCL-XL-TMD peptides co-localized with the mitochondria, although some overlap with the ER was observed (Figure S2). Surprisingly, MCL-1-TMD was primarily localized to the ER, but also in the cytosol. We quantified the co-localization by calculating the Pearson’s correlation coefficient for individual cells between TMD peptides and ER- and mitochondrial markers respectively (Figure 2C). BOK-TMD and BCL-2-TMD showed the highest correlation coefficient with the ER among the TMDs tested with a mean of r_BOK-TMD/ER_ = 0.50 and r_BCL-2-TMD/ER_ = 0.42 comparable to the ER-specific cb5-TMD peptide (mean r_cb5-TMD/ER_ = 0.40).

**Figure 2:**
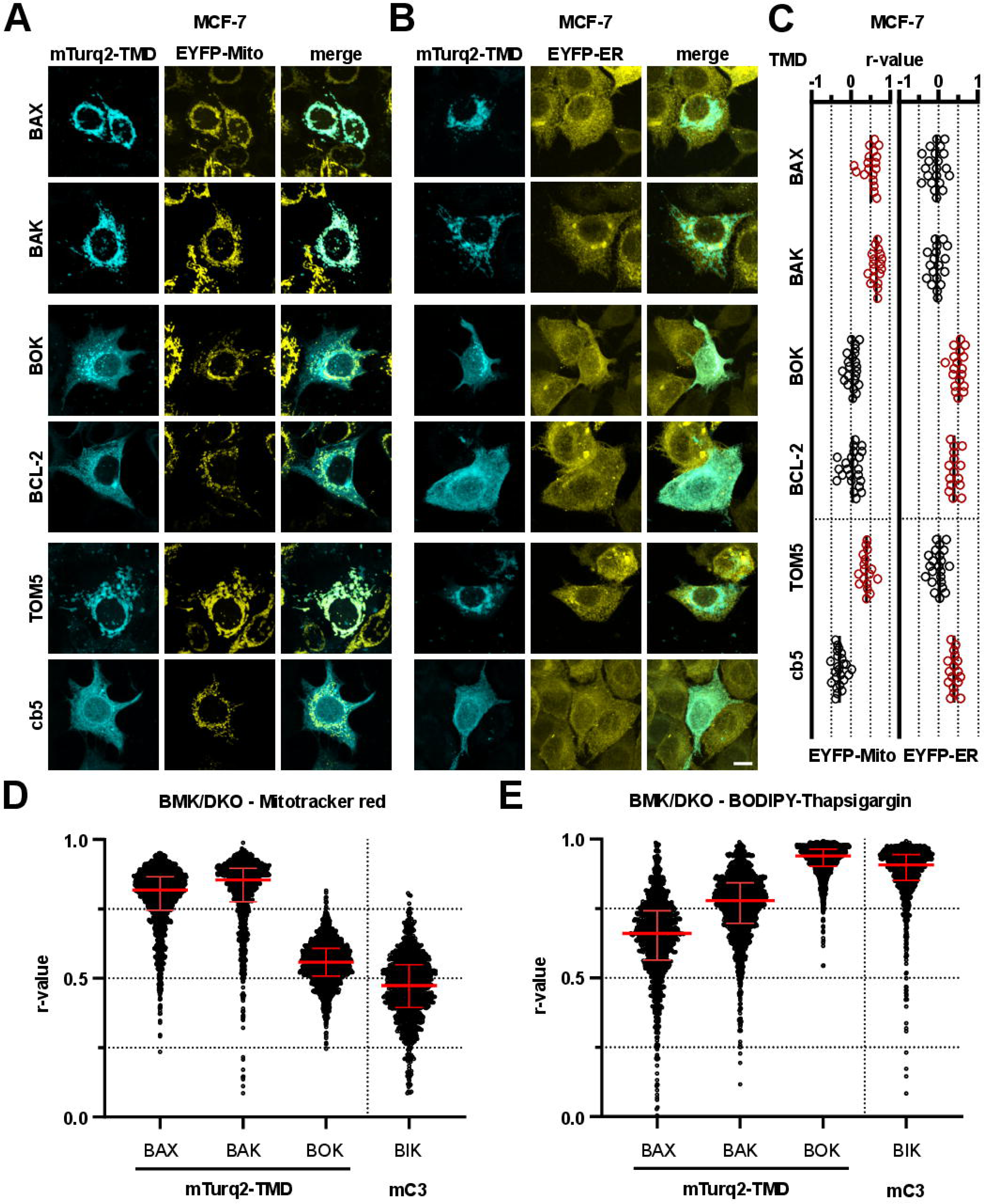
BCL-2-TMD and BOK TMD are predominantly ER-localized. **A, B** - Subcellular localization of TMD peptides. MCF-7 cells stably expressing (A) EYFP-Mito or (B) EYFP-ER were transfected with plasmids for the expression of mTurquoise2 (mTurq2)-labelled Bcl-2 TMD peptides. Cells were fixed after 24 h and TMD localization was analysed by cLSM. Images are maximum projection of z-stacks representative of three independent experiments. Scale bar = 10 µm. **C** - Quantitative analysis of the co-localization from cLSM in A, B (three independent experiments). Scatter plots show Pearson’s r correlation coefficients of in total ≥ 15 single cells. Mean is marked as a vertical line. Data with higher Pearson’s r (for EYFP-Mito or EYFP-ER) is colored red. **D, E** - Subcellular localization analysis of effector TMD peptides in BMK/DKO cells. BMK/DKO cell lines expressing mTurquoise2-BAX-TMD, -BAK-TMD and -BOK-TMD were labelled with DRAQ5 (nuclei) and (D) Mitotracker red or (E) BODIPY-Thapsigargin. Images were acquired using an Opera Phenix spinning disk microscope, cells were analyzed and Pearson’s r calculated. BAX-TMD and BAK-TMD predominantly localize to mitochondria, while BOK-TMD localizes to ER. Shown is median + IQR of n ≥ 1000 cells.

To further validate the subcellular localization of BAX-TMD, BAK-TMD and BOK-TMD, we generated cell lines from BAX-/-/BAK-/- baby mouse kidney cells (BMK/DKO) (Degenhardt et al. 2002) exogenously expressing mTurquoise2-conjugated N-terminally to BAX-TMD, BAK-TMD or BOK-TMD. In BMK/DKO cells, BAX-TMD and BAK-TMD displayed a mitochondria-like distribution, whereas BOK-TMD showed an ER-like distribution as imaged by the Opera Phenix confocal microscope (Figure S2). For high content analysis of TMD co-localization with mitochondria and ER markers, BMK/DKO cell lines were stained with Draq5 (nuclei), Mitotracker red (mitochondria) and BODIPY-Thapsigargin (ER) before imaging. The Pearson’s correlation coefficient (r) in the cytoplasmic region (excluding nuclei) of at least 1000 individual cells was calculated (Figure 2D, E). As a positive control for ER localization, a BMK/DKO cell line stably expressing mCerulean3-BIK, an ER-localized BH3-only protein (Osterlund et al. 2023), was included. Expectedly, BAX-TMD and BAK-TMD showed predominant localization at the mitochondria (mean r_BAX-TMD/Mito_ = 0.81 and r_BAK-TMD/Mito_ = 0.79), whereas mitochondrial localization of BIK and BOK-TMD was less pronounced (mean r_BOK-TMD/Mito_ = 0.55, r_BIK/Mito_ = 0.46). On the other hand, BAX-TMD and BAK-TMD displayed a less significant localization at ER (mean r_BAX-TMD/ER_ = 0.64 and r_BAK-TMD/ER_ = 0.76) while in contrast, BIK and BOK-TMD clearly strongly localized to the ER (mean r_BOK-TMD/ER_ = 0.92 and r_BIK/ER_ = 0.88). Thus, in BMK/DKO cells, BAX-TMD and BAK-TMD preferentially localize at the mitochondria, whereas BOK-TMD associates with the ER.

Since we found interaction of BOK-TMD and BCL-2-TMD in the split-luciferase assay, we next analyzed co-localization of mCitrine-BOK-TMD and mTurquoise2-BCL-2-TMD in MCF-7 cells. Using Mitotracker staining or co-expression of mCarmine fused to the ER targeting sequence of Calreticulin as an ER marker (Fabritius et al. 2018), we found visible and quantifiable co-localization of BOK-TMD and BCL-2-TMD at the ER with r_BOK-TMD/BCL-2-TMD_ = 0.52 (Figure 3 A, B).

**Figure 3:**
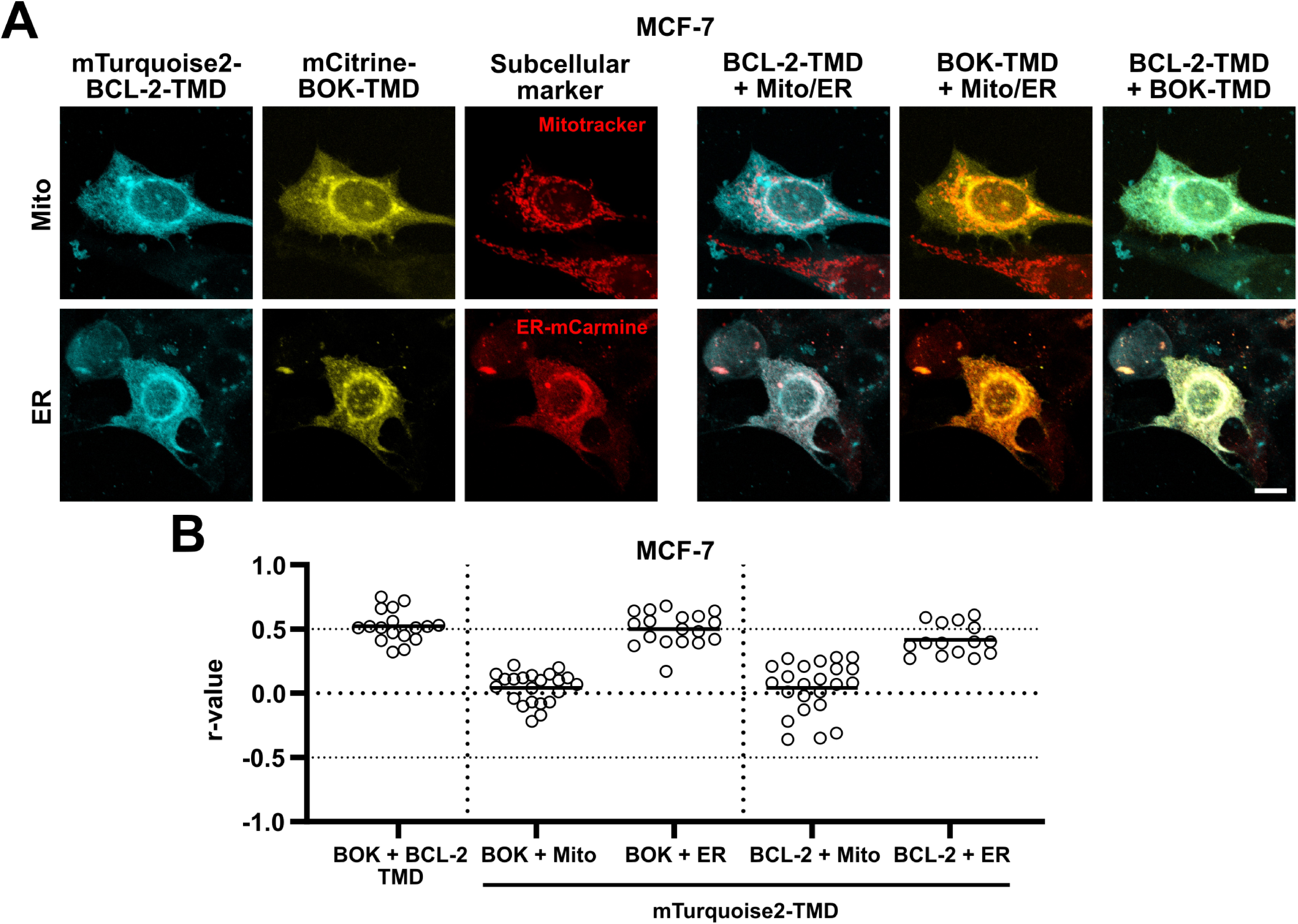
BOK-TMD and BCL-2-TMD co-localize at the ER. **A** - Co-localization of BOK-TMD and BCL-2-TMD at the ER. MCF-7 cells were transfected with plasmids for the expression of mTurquoise2-BCL-2-TMD and mCitrine-BOK-TMD. Cells were stained with Mitotracker red (upper panels) or co-transfected with plasmids for the expression of mCarmine-ER (lower panel). Cells were fixed after 24 h and TMD localization was analysed by cLSM. Images are maximum projection of z-stacks representative of three independent experiments. Scale bar = 10 µm. **B** - Quantitative analysis of co-localization between mTurquoise2-BCL-2-TMD and mCitrine-BOK-TMD from cLSM in D. Scatter plots show Pearson’s r correlation coefficients of in total ≥ 15 single cells from three independent experiments determined using Fiji software. Co-localization data between mTurquoise2-BCL-2-TMD/-BOK-TMD and EYFP-Mito and EYFP-ER from Figure 2 is shown for comparison. Mean is marked as a horizontal line. BCL-2-TMD and mCitrine-BOK-TMD co-localize at the ER.

These analyses show that BAX-TMD and BAK-TMD both localize to the mitochondria providing a valid explanation for their poor interaction with BCL-2-TMD. Furthermore, BOK-TMD and BCL-2-TMD co-localize at the ER giving a rational basis for the newly identified interaction.

### TMDs of BOK and BCL-2 are critical for their co-localization at the ER

Classically, interaction of full-length BCL-2 with BAX and BAK is understood to result from binding of the BH3 domain of BAX and BAK to the hydrophobic groove of BCL-2 (Ding et al. 2010). However, TMD-mediated interaction of Bcl-2 family proteins has been reported repeatedly (Todt et al. 2013; Andreu-Fernández et al. 2017; Lucendo et al. 2020). In order to tackle the question to which extent the interaction between BH3 domain and hydrophobic groove impacts co-localization of BCL-2 and effector proteins BAX, BAK and BOK, we utilized vectors for the expression of EGFP-fused full-length proteins BAX, BAK and BOK and chimeric BCL-2 proteins with swapped transmembrane domains (schematic in Figure 4A, B).

**Figure 4:**
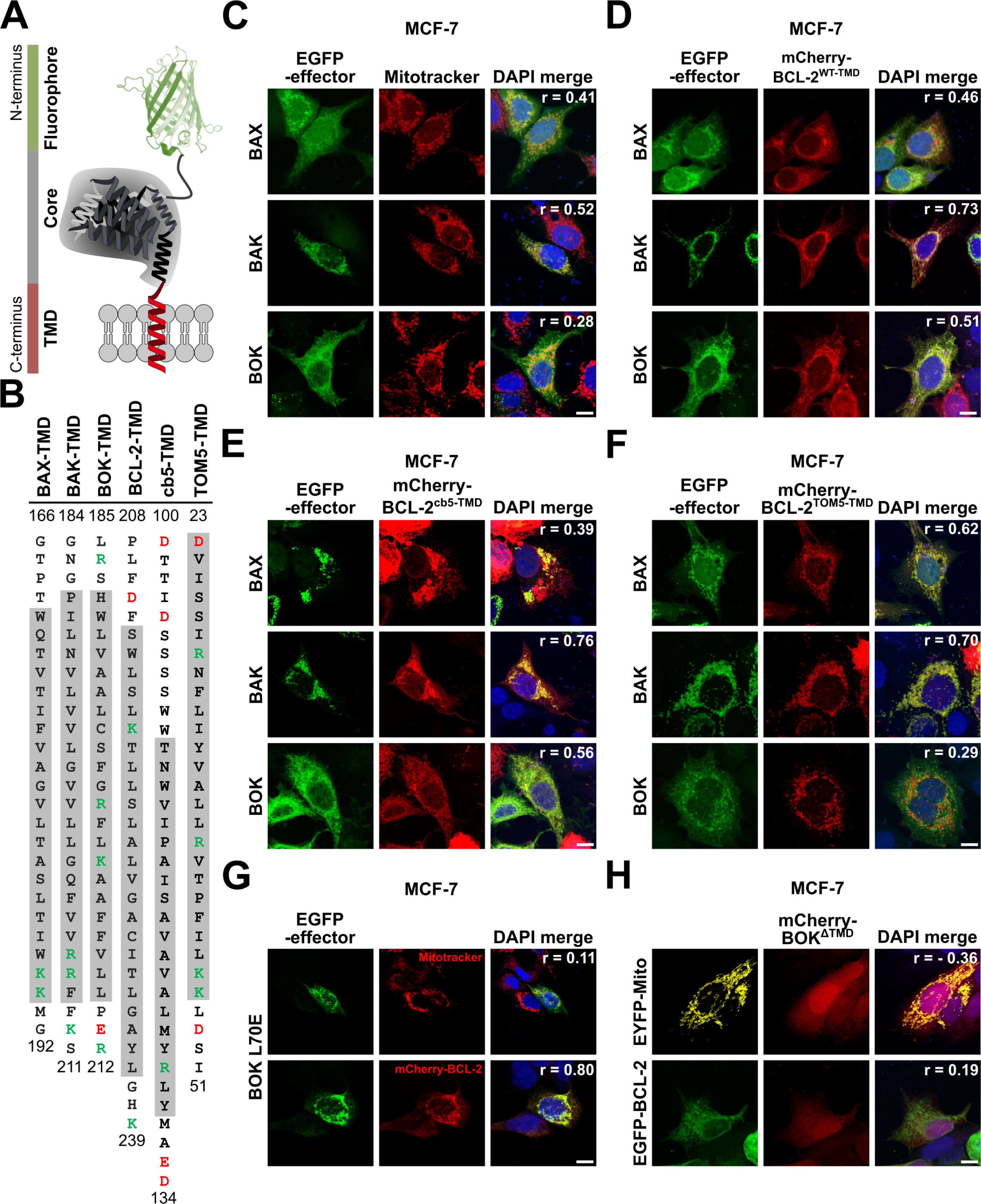
BOK and BCL-2 co-localization at the ER is dictated by their TMDs. **A** - Schematic depiction of fluorophore-tagged full-length Bcl-2 proteins. **B** - Amino acid sequences of BAX-, BAK-, BOK-, BCL-2-, cb5- and TOM5-TMD. Positively charged amino acids are labelled green, negatively charged amino acids are labelled red. Alpha-helical sequence sections are marked in grey. **C – F** - Co-localization of BAX, BAK, and BOK with (chimeric) BCL-2 protein. Co-localization of BOK and BCL-2 is TMD-dependent. MCF-7 cells were transfected with plasmids for the expression of EGFP-BAX, EGFP-BAK or EGFP-BOK. Cells were either stained with Mitotracker red (C) or co-transfected with plasmids for the expression of mCherry-BCL-2 wild-type (D, WT) or chimeric mCherry-BCL-2 proteins containing cb5-TMD (E) or TOM5-TMD (F). After 18 h, cells were fixed and mounted using DAPI-containing mounting medium followed by cLSM. (G) BOK L70E mutation does not alter BOK and BCL-2 co-localization. MCF-7 cells were transfected with plasmids for the expression of EGFP-BOK L70E. Cells were stained with Mitotracker red (upper panels) or co-transfected with plasmids for the expression of mCherry-BCL-2 (lower panels) before fixation after 18 h, mounting on slides using DAPI-containing mounting medium and cLSM. (H) TMD sequence is essential for membrane insertion of BOK and co-localization with BCL-2. MCF-7 cells were transfected with plasmids for the expression of mCherry-BOKΔTMD in combination with EYFP-Mito (upper panel) or EGFP-BCL-2 (lower panel). After 18 h, cells were fixed and mounted on slides using DAPI-containing mounting medium followed by cLSM. (C – H) Images are maximum projections of z-stacks representative of two independent experiments. R-values of respective middle sections are indicated in merged images. Scale bar = 10 µm.

MCF-7 cells with labelled mitochondria or ER that expressed EGFP-tagged full-length BAX, BAK or BOK were imaged by cLSM. Image analysis of cells which did not show clustered EGFP-signals revealed cytosolic and partially mitochondrial localization of BAX, exclusively mitochondrial localization of BAK, and ER localization of BOK (Figure 4C, Figure S4A). Interestingly, mCherry-BCL-2 showed co-localization with each EGFP-fused effector protein (Figure 4D). However, co-localization of EGFP-BAX and EGFP-BAK with mCherry-BCL-2 was detected at the mitochondria (r_BAX/BCL-2_ = 0.46, r_BAK/BCL-2_ = 0.73), whereas EGFP-BOK and mCherry-BCL-2 co-localized at the ER (r_BOK/BCL-2_ = 0.51). To challenge the role of the BH3 domain:hydrophobic groove interaction as mediator of co-localization, we analyzed co-localization of chimeric mCherry-BCL-2 harboring the transmembrane domain sequence of cytochrome b5 (cb5) or TOM5 with EGFP-fused effectors BAX, BAK, and BOK. Exchange of the TMD of BCL-2 to the ER-targeting cb5-TMD reduced co-localization of BCL-2 with BAX and BAK respectively, yet only in cells expressing low levels of BAX or BAK respectively (Figure S3B), while co-localization with BOK remained unchanged (r_BOK/BCL-2-cb5-TMD_ = 0.56, Figure 3E). Interestingly, in cells with BAX/BAK-clustering at the mitochondria, BCL-2cb5-TMD co-localized with these clusters (r_BAX/BCL-2-cb5-TMD_ = 0.39, r_BAK/BCL-2-cb5-TMD_ = 0.76, Figure 4E). Thus, we assume that interaction of the BCL-2 hydrophobic groove with the BH3 domain of active BAX and BAK is dominant over BCL-2-TMD-mediated localization and active BAX and BAK attract BCL-2^cb5-TMD^ to mitochondria. As expected, mCherry-BCL-2 with a conjugated TOM5-TMD was strictly localized to the mitochondria, effectively disrupting the co-localization with BOK (r_BOK/BCL-2-TOM5-TMD_ = 0.29, Figure 4F). Hence, while the co-localization of BCL-2 with BAX or BAK clusters at the mitochondria is BH3 domain-driven, BCL-2 localization at the ER and co-localization with BOK depends on the TMD rather than on the BH3 domain.

Corroborating this conclusion, mutation of the conserved leucine of the BH3 domain in EGFP-BOK (L70E mutation) did not affect co-localization with mCherry-BCL-2 (r_BOK L70E/BCL-2_ = 0.80, Figure 4G). In contrast, deletion of the BOK-TMD from mCherry-BOK (mCherry-BOKΔTMD) resulted in a diffuse cytosolic localization of BOK indicating inability of BOKΔTMD to integrate into membranes (Figure 4H). Co-expression of mCherry-BOKΔTMD with EGFP-BCL-2 did not change the diffuse localization of mCherry-BOKΔTMD. Thus, also TMD removal from BOK effectively abolishes co-localization with BCL-2 (r_BOK-ΔTMD/BCL-2_ = 0.19).

We conclude that co-localization of BCL-2 with BAX and BAK at mitochondria depends on exposure of the active effectoŕs BH3 domain. In contrast, analogous to BOK-TMD and BCL-2-TMD, also full-length BOK and BCL-2 co-localize at the ER, which does not depend on the BH3 domain of BOK but is driven by the TMD of BOK and BCL-2.

### Molecular dynamics reveals BOK- and BCL-2-TMD interactions in the ER membrane

Since the identified co-localization of BCL-2 and BOK at the ER corroborated the TMD interaction found in the split luciferase assay, we set out to investigate the interaction of BOK-TMD with BCL-2-TMD at molecular resolution by high-throughput molecular dynamics (MD) simulations. Because lipid composition of biomembranes substantially modulates interaction of transmembrane proteins (Pluhackova et al. 2016) we prepared a mimic (Pluhackova und Horner 2021) of endoplasmic reticulum membrane and studied homo- and hetero-oligomerization of BCL-2-TMD and BOK-TMD.

### BOK/BOK homodimerization and BOK/BCL-2 heterodimerisation

MD simulations of spontaneous BOK-BOK homodimerization at coarse-grained (CG) resolution resulted in three significant clusters (BOK/BOK-I, -II, and –III) of right-handed dimer structures (Figure 5A, Figures S4-S7, Table S6). In the most populated BOK/BOK-I comprising 55% of all CG structures, the two TMDs are shifted by ∼1.5 nm along the helix axis and residues K202, A203, F206, V207 and P210 of one BOK interact with residues H188, V191, A192, C195, and R199 of the other BOK (Figure 5A). The proximity of A192 from one and A203 from the other BOK TMD enables tight dimer packing. This asymmetric interaction interface appeared stable over the course of atomistic simulations, indicating its specificity. The strong tilt of the peptides in the bilayer and their shift relative to each other enable the charged residues K202, R199, of individual BOK peptides to snorkel to the lipid headgroups of different membrane leaflets (Rabe et al. 2016; Korn und Pluhackova 2022) (Figure S5).

**Figure 5:**
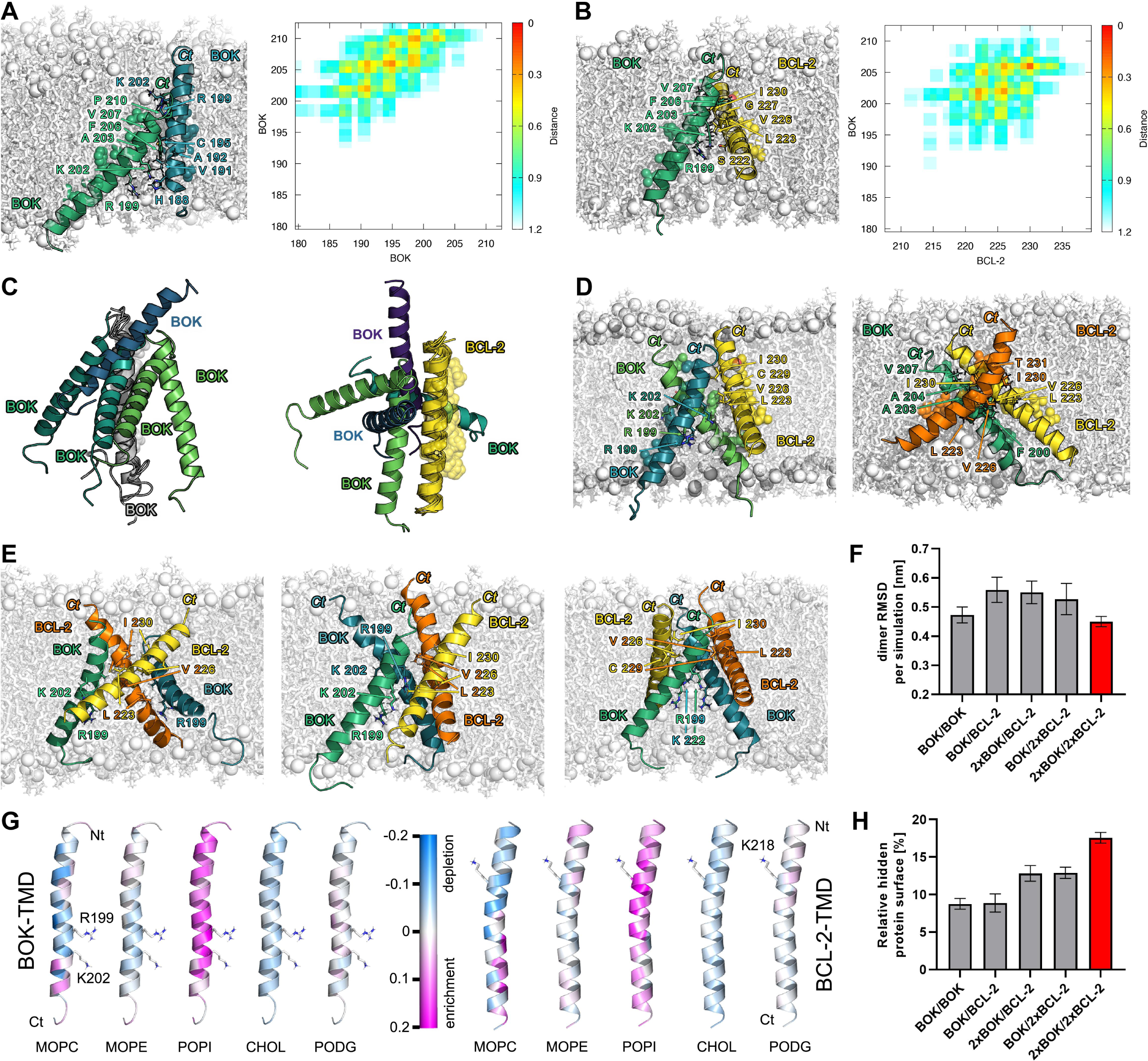
BOK-TMD and BCL-2-TMD interact in ER membranes. **A** - BOK/BOK-I homodimer (55% of formed BOK/BOK homodimers) exhibits an asymmetric interaction interface and causes membrane indentation of both membrane leaflets by snorkeling of R199 and K202 to the lipid headgroups. Left, snapshot after 1µs AA simulation, right contact map for the structure shown on the left. **B** - BOK/BCL2-I heterodimer (44% of formedBOK/BCL-2 heterodimers) is crossed shaped and stabilized by relatively small interaction interface formed by the C-terminal halves of the TMDs. Positively charged residues of both peptides, i.e. R199BOK, K202BOK and K218BCL-2 snorkel to the lipid headgroups of the cytoplasmic membrane leaflet only and cause a significant local thinning of the bilayer. Left, snapshot after 1µs AA simulation, right contact map for the structure shown on the left. **C** - The interaction partners bind to a reference TMD (BOK in grey in case of BOK/BOK homodimers on the left and BCL-2 in yellow for BOK/BCL-2 heterodimers on the right) in diverse positions, suggesting a possibility to form higher-order oligomers. **D** - Representative snapshots after 1µs AA simulations of the most often formed BOK/BOK/BCL-2 heterotrimers on the left and BOK/BCL-2/BCL-2 heterotrimers on the right. Water and ions were omitted for clarity, lipids are shown as white sticks with phosphorus atoms highlighted as spheres. Diverse important amino acid sidechains are shown as sticks and labeled. BOK TMDs are colored green or blue, BCL-2 TMDs are colored yellow or orange. **E** - Stable 2xBOK-TMD/2xBCL-2-TMD heterotetramers with "twisted" compact shape after AA simulation. On the left and on the right, alterating heterotetramers are shown with symmetric BCL-2-TMD and BOK-TMD homodimer in the center, respectively. In the middle a heterotetramer comprised of two BOK-TMDs followed by two BCL-2-TMDs is visualized. All tetramers are right handed. **F** - Average RMSD (estimated on a per dimer basis) values of AA simulations relative to the backmapped CG structure. The error bars denote SEM over individual simulations. N=13 for BOK-TMD homodimer, N=15 for BOK-TMD/BCL-2-TMD heterodimer, N=9 and 10 for 2xBOK-TMD/BCL-2-TMD and BOK-TMD/2xBCL-2-TMD, respectively and N=18 for 2xBOK-TMD/2xBCL-2-TMD. **G** - Lipid enrichment (magenta) or depletion (blue) per BOK-TMD (left) or BCL-2-TMD (right) residue in the heterotetramer simulations. The membrane indenting R199BOK, K202BOK and K218BCL-2 are highlighted as sticks. **H** - Protein surface occupied by protein-protein interactions relative to the total surface of all peptides for all simulation types. The error bars denote SEM and the number of simulations equals to that listed in F. In A, B, D and E water and ions were omitted for clarity, lipids are shown as white sticks with phosphorus atoms highlighted as spheres. The proteins are shown as cartoons with selected amino acid sidechains shown as sticks with label. BOK TMDs are colored green or blue, BCL-2 TMDs are colored yellow or orange.

Remarkably, the side of the BOK TMD helix containing predominantly large hydrophobic residues L190, A193, L194, F197, L201, A204 and L208 (shown in Figure S8) faces exclusively hydrophobic lipid tails in all clusters formed, avoiding protein-protein interaction as also confirmed by the negligible contributions of these residues to the interaction energy of the dimers (shown in Figure S9). This is also true for BOK-TMD/BCL-2-TMD interactions, where BCL-2 also exhibits a lipid facing side (Figure S8).

CG simulations of spontaneous BOK-TMD/BCL2-TMD dimerisation resulted in four clusters of dimers (Figures S10-S14, Table S7). The right-handed BOK/BCL2-I, shown in Figure 5B, was formed in 44%, the right-handed BOK-BCL2-II in 34%, the left-handed BOK-BCL2-III in 8% and the right-handed BOK/BCL2-IV in 7% of all dimers. With the exception of the structures from the rarely-formed BOK/BCL-2-IV cluster, all-atom simulations of BOK-TMD/BCL-2-TMD heterodimers have shown even larger conformational flexibility than atomistic simulations of BOK-TMD/BOK-TMD homodimers, resulting in significant reorientation of the two TMDs and even dissociation in 2 out of 12 simulations. Certain commonalities were identified for the crossed dimers BOK/BCL-2-I, II and III with the exception of the (nearly) dissociated conformations: BOK-TMD most often contacted L223^BCL-2^, V226^BCL-2^ or I230^BCL-2^, the backbone of BCL-2-TMD typically crossed BOK-TMD at A203^BOK^, and R199^BOK^, K202^BOK^, as well as K212^BCL-2^ snorkeled to the cytosolic membrane leaflet causing its deformation and local indentation. It is interesting to note, that none of the left-handed BOK/BCL-2-III dimers stayed left-handed after the atomistic simulation.

The number of interaction sweet spots in BOK/BOK and BOK/BCL-2 dimers (Figure 5C), which is an indicator of higher order oligomers (Han et al. 2016), taken together with high occurrence yet low conformational stability of BOK homodimers and even more strongly BOK/BCL2 heterodimers suggest, that BOK-BCL2 interactions likely prefer oligomeric structures. To test this hypothesis we have inserted two BOK-TMDs and two BCL-2-TMDs into the ER membrane mimic and studied their spontaneous association 50 times at CG level. After 50 µs 50% of the TMDs formed tetramers, 36% were trimers (in 13 simulations one BOK-TMD was stabilized by 2 BCL-2-TMD, and in 5 simulations 2 BOK-TMDs associated with one BCL-2-TMD), in 5% a heterodimer and in 2% BCL2 homodimer were formed. The trimers and tetramers were manually grouped in PyMOL and their stability studied by 1 µs AA simulation, each.

### 2xBOK/BCL-2 and BOK/2xBCL-2 trimers

Eventhough the most 2xBOK/BCL-2 and BOK/2xBCL-2 heterotrimers adopted a compact shape (Figure 5D, Figure S15) the heterotrimers appeared similarly dynamic as BOK/BCL-2 heterodimers, as measured by their RMSD over AA simulation relative to the CG structure (Figure 5F). The analysis of protein surface buried at the protein-protein interaction interfaces (Figure 5G) has shown that in case of trimers 1.5 more protein surface is buried as compared to the dimer interfaces. Combined with our observation of compact trimers, where each peptide contacts two other peptides, this hints to the fact that upon trimerization smaller interaction surfaces are formed. This can be explained by our visual observation of more crossed peptides in the trimers than in dimers where they typically form comparably long interaction surfaces.

### 2xBOK/2xBCL-2-TMD tetramers

Our CG simulations have shown tetramerization to be the preferred oligomerization state of two BOK-TMDs with two BCL-2-TMDs. In 27% of the analyzed tetramers, the TMDs formed compact right handed heterotetramers with slightly twisted shape enabling formation of extensive interaction interfaces. Three examples are shown in Figure 5E. In 19% and 5% crossed dimers of parallel hetero- or homodimers, respectively, were formed, reminding of a hash shape. In 22% and 7% of the tetramers a dimer of parallel and crossed-dimers arose (called halfhash from now on), which could be in intermediate between the former two tetramers (hash and twist). In 19% of the tetramers one peptide was attached in a LH manner to a RH trimer. Atomistic simulations of the twist, hash and intermediate tetramer-shapes have shown that the compact twist shape is the most favored (2/3 of the 18 AA simulations adopted this shape). Interestingly, in 4 other simulations hash or half-hash of homodimers appeared to be stable, contrary to the fact that all heterohashes and half-heterohashes rapidly evolved into compact twisted heterotetramers. The interaction energy plots (Figure S9) per BOK-TMD or BCL-2-TMD residue and visual inspection of the tetramers show a similar involvement of residues in the formation of interaction interfaces as in the dimers and trimers. The increase of the hidden surface area in tetramers by factor 2 relative to the dimers (Figure 5H) hints to further stabilization of the oligomer which is also confirmed by smallest RMSD values compared to dimers and trimers (Figure 5F). In reality the stability of the tetramers compared to dimers and trimers is even higher than the RMSD plot suggest, because the RMSD of larger proteins/protein assemblies is intrinsically higher than that of smaller protein structures (Irving et al. 2001).

Taken together, our multiscaling MD simulations show that BOK-TMD and BCL-2-TMD interact dynamically in the ER membrane by multiple interaction interfaces. The requirements for interaction are best met in tetramers, which is consistent with the experimentally observed optimal 1:1 ratio of the interacting BAX-TMDs in the split luciferase assay. Of note, in simulations containing two BOK-TMDs, BOK residues R199 and E211 contribution to protein-protein interaction energy is far higher than in simulations containing a single BOK-TMD (Figure S9), rendering these residues as targets for future mutagenesis studies with the aim to weaken BOK-TMD homooligomerization.

### Peptide-lipid interaction

The here generated ER-like membrane mimic consisting of charged 1-palmitoyl-2-oleoyl-sn-glycero-3-phosphoinositol (POPI), zwitterionic 1-myristoyl-2-oleoyl-sn-glycero-3-phosphocholine (MOPC) and 1-myristoyl-2-oleoyl-sn-glycero-3-phosphoethanolamine (MOPE) and uncharged 1-palmitoyl-2-oleoyl-sn-glycerol (PODG) and cholesterol allows analysis of the protein-lipid interaction, which often is a strong modulator of protein-protein interactions (Pluhackova et al. 2016; Friess et al. 2018) – also for Bcl-2 proteins (Lutter et al. 2000; Lucken-Ardjomande et al. 2008; Shamas-Din et al. 2015). Regardless of the oligomeric state of the TMDs, MD simulations in the ER-mimic show strong depletion of MOPC, slight depletion of cholesterol, slight enrichment of MOPE and strong enrichment of POPI for both TMDs (Figure 5G, Figure S20). PODG is neither depleted nor enriched in the vicinity of BCL-2-TMD and slightly enriched at BOK-TMD. The strong enrichment of negatively charged POPI likely results from the high amount of positively charged amino acids in the TMDs (BOK-TMD contains 3 Arg, 1 Lys, 1 Glu and 1 Asp; BCL2 TMD comprises 2 Lys and 1 Asp, Table S8, 9).

### BCL-2 antagonizes BOK-induced cell death TMD-dependently

BCL-2 is a bona-fide antagonist of BAX and BAK and effectively counteracts BAX and BAK-induced mitochondrial pore formation and cell death (Ding et al. 2010). In contrast to BAX and BAK, cell death induction by BOK is less thoroughly understood. However, it is generally recognized that either BOK overexpression or blocked degradation induces apoptosis (Einsele-Scholz et al. 2016; Llambi et al. 2016). So far only few studies support a direct regulation of BOK-mediated apoptosis by anti-apoptotic Bcl-2 proteins (Stehle et al. 2018; Lucendo et al. 2020). On the contrary, it is assumed that anti-apoptotic Bcl-2 proteins do not affect BOK-induced cell death (Carpio et al. 2015; Llambi et al. 2016).

While interaction of BOK with Bcl-2 proteins in general and BCL-2 in particular is disputed, the above results strongly suggest that BOK and BCL-2 interact via their TMDs. Consequently, we investigated whether BCL-2 attenuates BOK-induced cell death. To this end, we performed siRNA-mediated knock-down of BCL-2 in HEK293 cells and subsequently transfected these cells to express EGFP-BOK. Expectedly, flow cytometric analysis showed that overexpression of BOK substantially increased the proportion of Annexin-V-APC positive, i.e. apoptotic, cells to almost half of transfected (EGFP positive) cells (37.9% APC+ cells) 18 h post transfection (Figure 6A). Knock down of BCL-2 resulted in an increased proportion of apoptotic cells as compared to control cells (49.7% APC+ cells) indeed indicating an antagonistic role of BCL-2 in BOK overexpression-induced cell death. Because knock down reduced but did not abolish BCL-2 expression as analyzed by Western Blot (Figure S5C) we aimed to verify BCL-2-mediated inhibition of BOK-induced apoptosis by CRISPR/Cas9-mediated knock-out. We co-transfected HEK293 to express CRISPR/Cas9 for BCL2 knock-out together with EGFP-BOK and subsequently analyzed apoptosis induction in EGFP-positive cells by Annexin-V-APC staining. Flow cytometric analysis detected 50.2% apoptotic EGFP-BOK-expressing cells which was enhanced to 60.6% in BCL-2 knock-out cells (Figure S5B). Next, we analyzed BOK-induced apoptosis in the BAX- and BCL-2-deficient cell line DU145 (Figure S5A). Overexpression of EGFP-BOK induced exposure of phosphatidyl serine in 12.3% of DU145 cells after 18 h and 20.7% of DU145 cells 42 h post transfection as assessed using flow cytometry (Figure 6B, Figure S5D). Apoptosis induction was efficiently reduced by co-transfection of mCherry-BCL-2 resulting in 5.7% APC+ cells after 18 h and 8.2% APC+ cells 42 h post transfection (Figure 6B, Figure S5D). In line with apoptosis induction, cleavage of the caspase substrate PARP indicating apoptosis execution was readily detected after EGFP-BOK overexpression which was reduced by co-transfection with mCherry-BCL-2 (Figure 6D, E). Moreover, assessing activity of apoptosis specific caspase-3/7 in EGFP-BOK expressing DU145 cells we detected an increase of 32% in caspase-3/7 activity compared to control cells while caspase-3/7 activity was decreased by 17% by co-expression of BCL-2 compared to control (Figure 6F).

**Figure 6:**
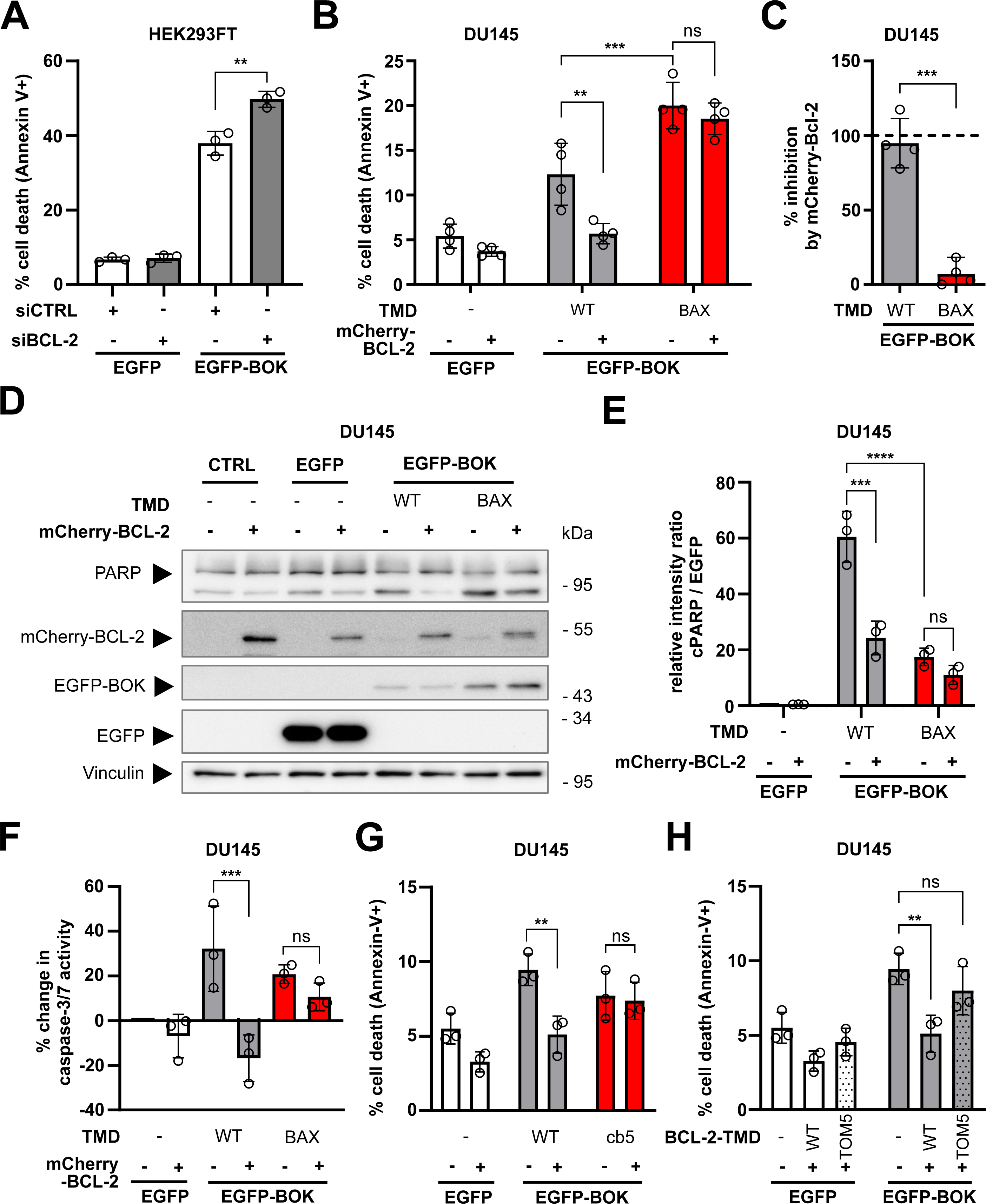
Inhibition of BOK-mediated apoptosis by BCL-2 depends on BOK-TMD. **A** - BOK-induced apoptosis is pronounced after knock-down of BCL-2. HEK293FT were transfected with non-targeted (siCTRL) or BCL-2-targeted (siBCL-2) small-interfering RNA for 24 h and subsequently with plasmids for the expression of EGFP or EGFP-BOK for additional 18 h. Apoptotic cells were detected by Annexin-V-APC staining by flow cytometry. Graph shows mean ± sd from three independent experiments. **B** - Co-expression of mCherry-BCL-2 reduces apoptosis induction by BOK but not BOKBAX-TMD. DU145 cells were transfected with plasmids for the expression of EGFP, EGFP-BOK, or EGFP-BOKBAX-TMD without or with mCherry-BCL-2. After 18 h, cells were stained with Annexin-V-APC and EGFP+/Annexin-V+ cells were detected by flow cytometry. **C** - Relative inhibition of BOK wild-type or BOKBAX-TMD by BCL-2 as calculated from (B). **D** - Abundance of cleaved PARP is enhanced by BOK and BOKBAX-TMD. Expression of BCL-2 reduces abundance of cleaved PARP in BOK-expressing cells but not in BOKBAX-TMD expressing cells. DU145 cells expressing EGFP, EGFP-BOK, or EGFP-BOKBAX-TMD alone or in combination mCherry-BCL-2 were analyzed by Western Blot for the expression of EGFP(-BOK), BCL-2 and PARP. Vinculin served as loading control. Representative Blot from three independent experiments. **E** - Densitometric analysis of three independent experiments as shown in (E). **F** - BOK-induced caspase-3/7 activity is reduced by BCL-2, but not for BOKBAX-TMD. DU145 cells were co-transfected with plasmids for the expression of EGFP, EGFP-BOK or EGFP-BOKBAX-TMD with or without mCherry-BCL-2. After 18 h, caspase-3/7 activity was assessed with the Caspase-Glo 3/7 assay kit (Promega). Percent change in caspase-3/7 activity compared to EGFP control is shown as mean ± sd from three independent experiments. **G** - mCherry-BCL-2 reduces apoptosis induction by BOK but not BOKcb5-TMD. DU145 cells were transfected with plasmids for the expression of EGFP, EGFP-BOK, or EGFP-BOKcb5-TMD without or with mCherry-BCL-2. After 18 h, cells were stained with Annexin-V-APC and EGFP+/Annexin-V+ cells were detected by flow cytometry. (H) Inhibition of BOK-induced cell death by BCL-2 is impaired for BCL-2TOM5-TMD. DU145 cells were transfected with plasmids for the expression of EGFP or EGFP-BOK in combination with mCherry-BCL-2 or mCherry-BCL-2TOM5-TMD. Cells were stained with Annexin-V-APC and EGFP+/Annexin-V+ cells were detected by flow cytometry. (A, B, C, E, F, G) Graphs show mean ± sd from three (A, E, F, G) or four (B, C) independent experiment

With the BAX and BCL-2 deficient cell line DU145, we sought to further confirm whether the interaction of BOK-TMD and BCL-2-TMD is relevant for the observed cell death inhibition. Since the BAX/BCL-2 co-localization seems to be mediated by BH3 domain:hydrophobic groove rather than depend on their TMDs, we analyzed whether cell death induction by chimeric BOK with BAX-TMD is reduced by co-expression of BCL-2. The exchange of BOK-TMD to BAX-TMD shifted localization of BOK^BAX-TMD^ to mitochondria (Figure S3C) and also potentiated cell death induction by BOK overexpression from 12.3% to 20.0% APC+ cells 18 h post transfection (Figure 6B). Manifesting a clear role of TMD interaction, cell death induction by BOK^BAX-TMD^ was indeed not altered by co-expression of BCL-2 (Figure 6C, 94% inhibition of EGFP-BOK-induced cell death reduced to 7.3% inhibition for EGFP-BOK^BAX-TMD^). Failure of BCL-2 to inhibit BOK^BAX-TMD^ induced cell death was also reflected by unchanged level of cleaved PARP and Caspase-3/7 activity after expression of EGFP-BOK^BAX-TMD^ alone compared to co-expression with mCherry-BCL-2 (Figure 6D-F).

Interestingly, Western Blot analysis showed higher expression of mitochondria-localized EGFP-BOK^BAX-TMD^ compared to ER-localized wild-type EGFP-BOK (Figure 6D). Densitometric analysis of cleaved PARP per EGFP suggests that, although showing a higher expression level, BOK^BAX-TMD^ induces apoptosis less efficiently than wild-type BOK (rel. intensity cPARP/EGFP-BOK^BAX-TMD^ = 17.5; rel. intensity cPARP/EGFP-BOK = 60.5, Figure 6E). The increased pro-apoptotic activity of BOK with BAX-TMD hence may result from higher expression BOK^BAX-TMD^ due to increased stability or reduced degradation.

To further challenge the TMD dependency of BCL-2-mediated inhibition of BOK-induced apoptosis, we analyzed cell death induction by BOK carrying the wild-type TMD or the TMD of cb5 in the presence or absence of BCL-2. As for BOK^BAX-TMD^, the low cell death induction by BOK^cb5-TMD^, (7.7% APC+ cells), was not reduced by BCL-2 (7.4% APC+ cells, Figure 6G). Vice-versa, exchanging the TMD of BCL-2 to the TMD of MOM-localized protein TOM5 reduces its calculated inhibitory capacity towards BOK-induced apoptosis when co-transfected in DU145 cells by 67% (cell death increases from 5.1% APC+ cells to 8.0% APC+ cells, Figure 6H).

In conclusion, knock-down or knock-out of BCL-2/BCL-2 increased BOK-induced apoptosis, while co-transfection with BCL-2 reduced cell death induction by overexpression of BOK. The BCL-2-mediated inhibition of BOK-induced apoptosis specifically depends on their TMD. Thus, to the best of our knowledge, for the first time our data uncover a functional involvement of BOK-TMD and BCL-2-TMD interaction in apoptosis signaling.

## Discussion

BH3 mimetics which made it possible to target interactions of Bcl-2 proteins equipped humanity with new powerful options to stand against cancer. BCL-2 as one of the most well-studied anti-apoptotic family members was shown to be a worthwhile target for anti-cancer therapy: cancer cell-protective BCL-2 overexpression and hence sequestration of pro-apoptotic proteins is a driving force of tumorigenesis in hematopoietic malignancies like chronic lymphoid leukemia (CLL) (Robertson et al. 1996; Del Gaizo Moore et al. 2007). The identification of BCL-2 as an oncogene ignited the development of BH3 mimetics and eventually the specific BCL-2 inhibitor ABT-199/Venetoclax. Venetoclax was the first FDA-approved BH3 mimetic for the use as a single agent in CLL and later in combination with azacidine, decitabine or low dose cytarabine in acute myeloid leukemia (AML) (Roberts et al. 2016; DiNardo et al. 2019).

Importantly, the Bcl-2 protein family is furthermore a valuable target for anti-cancer therapy. Recent studies revealed that Venetoclax in addition to potently blocking BCL-2 causes cellular responses that support anti-tumor activity including metabolic reprogramming and activation of the integrated stress response (Roca-Portoles et al. 2020; Weller et al. 2022). Undoubtedly, identifying new target structures within the Bcl-2 family will aid in the endeavour to develop even more specific treatment options.

Here, we established a bimolecular split-luciferase assay and elucidated a novel interaction of BCL-2 and BOK via their TMDs. The presented data furthermore indicates functional relevance of the TMD interaction in apoptosis regulation. Surprisingly, the interactions of BCL-2-TMD and BOK-TMD takes place at ER membranes and MD simulations in an ER membrane mimic revealed formation of stable hetero-tetramers (2xBOK-TMD/2xBCL-2-TMD), structurally dynamic BOK-TMD homodimers, BCL-2-TMD/BOK-TMD hetero-dimers and hetero-trimers. In vitro experiments support a functional significance of the BCL-2-TMD interaction with BOK-TMD as BCL-2-mediated inhibition of BOK-induced cell death was clearly dependent on BOK-TMD sequence.

The newly described interaction of BOK-TMD with BCL-2-TMD is particularly intriguing since the interaction of BOK via it’s BH3 domain is still debated. In fact, specific BH3 domain interaction of BOK with anti-apoptotic Bcl-2 proteins has not yet been reported. However, accompanying the identification of BOK a yeast two-hybrid assay found MCL-1 and A1 as BOK interaction partners (Hsu et al. 1997). Interaction of BOK and MCL-1 was also confirmed by other groups (Llambi et al. 2016; Stehle et al. 2018). More recently, interaction of BOK with MCL-1 via the TMD domain was shown to modulate apoptosis (Lucendo et al. 2020). In the same study, Lucendo et al. show that MCL-1-TMD can tether BOK-TMD to mitochondria and increase the number of mitochondria-associated membranes (MAMs). Consequently, interaction via the TMD appears to be especially important in case of BOK as compared to other Bcl-2 proteins. However, in the present study MCL-1-TMD was localized to the ER rather than mitochondria and split-luciferase assay did not indicate interaction with BOK-TMD. In support of the importance of BOK-TMD for protein-protein interaction, abrogation of BH3-mediated interaction by mutation of the conserved L70 in BOK did not alter the co-localization with BCL-2.

Also the mode of BOK-induced apoptosis is controversial. Some reports claim BOK to be a pro-apoptotic effector like BAX and BAK which kill cells by forming pores in the MOM (Einsele-Scholz et al. 2016; Fernández-Marrero et al. 2017; Shalaby et al. 2023). Other studies deny BOK any independent apoptosis-mediating function and suggest that BOK functions upstream of BAX and BAK (Echeverry et al. 2013; Carpio et al. 2015). A solution to this conundrum might be offered by recent reports showing that BOK localizes to the ER and is involved in calcium transfer to mitochondria via mitochondria-associated membranes (MAMs) thereby promoting MOM depolarization and apoptosis (Carpio et al. 2021). Hand in hand with BOK localization at the ER, a role of BOK in ER stress response has also been reported (Echeverry et al. 2013; Carpio et al. 2015; Walter et al. 2022). In concordance, in the present study BOK and BCL-2 co-localize at the ER suggesting a regulatory role of their TMD interaction at the ER or MAMs. In line with ER localization, BOK and BCL-2 both interact with IP3R calcium channels with opposing effects on apoptosis (Rong et al. 2009; Monaco et al. 2012; Schulman et al. 2016). In fact, for BCL-2 the TMD is necessary and sufficient to bind and inhibit IP3R (Ivanova et al. 2016). However, also IP3R-binding via the BH4 motif of BCL-2 was shown to promote cell survival and decrease apoptosis by mediating Ca2+ leakage from the ER to the cytosol and inhibiting Ca2+ release upon apoptosis induction (Rong et al. 2009; Monaco et al. 2012). On the other hand, stability as well as pro-apoptotic capacity of BOK largely depend on the binding to IP3R as Schulman et al demonstrated that virtually all cellular BOK is bound to IP3R and rapidly degraded via the proteasome when released (Schulman et al. 2016). The authors furthermore revealed that BOK deletion leads to fragmentation of mitochondria and thereby BOK affects bioenergetics suggesting further apoptosis-unrelated functions of BOK (Schulman et al. 2019). The opposing role of BOK and BCL-2 in apoptosis by binding to IP3Rs and their function in Ca2+ signaling indicate ER and MAMs as the functional hub for their TMD interaction, e.g. by competing for binding sites at IP3Rs and hence regulation of IP3R function.

The function of BOK and BCL-2 in Ca2+ signaling at the ER does not exclude an involvement of TMD interaction in canonical pore formation by BOK in the MOM since Bcl-2 family proteins that mainly reside at the ER, e.g.BCL-2 and the BH3-only protein BIK, translocate to mitochondria and interact with mitochondrial localized Bcl-2 family proteins thus regulating MOMP (Lalier et al. 2021; Osterlund et al. 2023). The presented high-throughput MD simulations of BOK-TMD and BCL-2-TMD interaction in an ER membrane mimic indeed indicate that BCL-2-TMD binds to BOK-TMD and preferentially forms higher order oligomers. Interestingly, in these simulations BCL-2-TMD or BCL-2-TMD homodimer associate with BOK-TMD homodimers which indicates that BCL-2 is recruited to BOK homodimers and inhibits BOK oligomerization. Along these lines, although heterodimers and heterotrimers showed high transformability, the tetramers consisting of two BOK-TMD and two BCL-2-TMD peptides showed the highest structural stability among all oligomers studied, further supporting a role of TMD interaction in BOK oligomerization. Additionally, snorkeling of positively charged lysine and arginine of BOK to the lipid headgroups resulted in local membrane indentation demonstrating the capacity of BOK-TMD to modulate membrane structure. Vice versa, membrane composition likely is a major factor that influences TMD interaction, since negatively charged POPI accumulates in the vicinity of both BOK-and BCL-2-TMD. Enrichment of negatively charged POPI at TMDs is in line with reports showing that negatively charged cardiolipin (CL) in the MOM significantly influences interaction of Bcl-2 proteins with mitochondrial membrane (Lutter et al. 2000; Lucken-Ardjomande et al. 2008; Shamas-Din et al. 2015). CL interacts with positively charged amino acids in TMDs of which some can be found in BOK and BCL-2 as well (R199^BOK^, K202^BOK^, and K218^BCL-2^). Also, in our simulations these charged residues attracted negatively charged POPI lipids. Interestingly, K218BCL-2 almost never participated in the interactions of the two TMD peptides although polar/charged residues typically act as intermolecular locks in helix-helix interaction (Senes et al. 2000; Curran und Engelman 2003). However, also other defined motifs which consist foremost of small hydrophobic amino acids like glycine, alanine, serine or threonine are frequently involved in helix-helix interactions and TMD oligomers (Russ und Engelman 2000; Senes et al. 2000; Kim et al. 2005; Gössweiner-Mohr et al. 2022). Some of these motifs are found in the BCL-2-TMD (Figure S3D, VI4 (pos. 227 - 230) and GG4 (pos.234 – 237)). Intriguingly, none of these frequent motifs could be found in BOK-TMD. However, the sequence of BOK-TMD contains a peculiar high number of phenylalanine residues (F197, F200, F205, F206). This is especially interesting with regard to studies which postulate an interaction-promoting effect of phenylalanine in TMDs (Unterreitmeier et al. 2007; Kwon et al. 2016). Studying these helix-helix interaction facilitating motifs and residues of BCL-2-TMD and BOK-TMD by in silico approximations should grant insight into their role in the interaction of BOK-TMD with BCL-2-TMD.

Also post-translational modification of Bcl-2 TMDs strongly impact localization and function of Bcl-2 proteins, exemplified by phosphorylation of the BAX-TMD at S184 by Akt (Guedes et al. 2013; Kale et al. 2018a) that prevents integration into the mitochondrial outer membrane. Intriguingly, the TMD mutation G179E in BAX promotes resistance to anti-cancer treatment by rendering cells less susceptible to Venetoclax (Fresquet et al. 2014). Thus, mutation and post-translational modification of the TMD in Bcl-2 proteins are highly relevant for apoptosis regulation. In addition, as shown for the BH3-only proteins BIML as well as PUMA, high-affinity binding to anti-apoptotic Bcl-2 proteins depends on a “double-bolt lock” mechanism mediated by both TMD and BH3 domain (Liu et al. 2019; Pemberton et al. 2023). Hence, these results might serve as a role model for a new generation of Bcl-2 protein-targeting drugs that target interaction not only via BH3 domain:hydrophobic groove but also the TMD. Interestingly, we found several tumor-specific TMD mutations in the COSMIC database (cancer.sanger.ac.uk/cosmic) underpinning the impact of Bcl-2 TMDs on the apoptosis signaling network by affecting localization, interaction or “double-bolt lock” mechanism.

Taken together, we unveil two new aspects of apoptosis regulation: Firstly, we have discovered that the inhibitory role of BCL-2 on BOK-mediated apoptosis crucially depends on the interaction via their transmembrane domains. Secondly, BOK and BCL-2 are localized at the ER via their TMDs which indicates that BOK-induced apoptosis regulation originates at the ER, not the mitochondria. Our findings support the emerging role of TMD interactions as an important structural component of the Bcl-2 interaction network. We conclude that apoptosis dysregulation as well as survival strategies in healthy and transformed cells are inherently influenced by Bcl-2 TMDs in two ways: i) subcellular localization of Bcl-2 proteins and ii) interaction of specific Bcl-2 protein via their TMDs. Simultaneous targeting of BH3 domain:hydrophobic groove and the TMD interface of Bcl-2 proteins, therefore, harbours not yet exploited potential for anti-cancer therapy e.g. by increasing drug specificity.

## Materials and methods

### Protein sequences

Protein names, function and Uniprot Entries (www.uniprot.org) of protein sequences used for generation of expression vectors are listed in Table S1A. TMD sequences used are listed in Table S1B.

### Antibodies and reagents

Antibodies used were: anti-LgBiT (Promega #N7100), anti-GFP (for EGFP/mTurq2, Santa Cruz #sc-9996), anti-GAPDH (Cell Signaling Technology (CST) #2118), anti-BOK (abcam ab186745), anti-BCL-2 (CST #15071), anti-PARP (CST #9542), anti-Vinculin (protein tech #66305-1-Ig), anti-MCL-1 (CST #5453), anti-BCL-XL (CST #2762), anti-BAX (CST #2772), anti-BAK (CST #3814), anti-β-actin (Sigma #A5316), secondary horseradish peroxidase-conjugated antibodies anti-mouse (CST #7076) and anti-rabbit (CST #7074); Vectors for expression of EGFP, EYFP-Mito (EYFP fused to mitochondrial targeting sequence of cytochrome C oxidase subunit VIII) and EYFP-ER (EYFP fused to Calreticulin ER-targeting sequence) were from Clontech (Takara Bio, Kusatsu, Japan); Vector for CRISPR/Cas9-mediated knock-out of BCL-2 was plentiCTRSPRv2 (Addgene #52961) with gRNA targeting exon 1 in BCL2; reagents for cell stainings and fixation used were Mitotracker green / red CMXRos (Thermo Fisher Scientific, Waltham, MA, USA), fixation solution ROTI Histofix 4% formaldehyde (Carl Roth, Karlsruhe, Germany), ProLong Diamond Antifade Mountant with DAPI (Thermo Fisher Scientific). Nuclear and ER staining for high content analysis was done with DRAQ5 and BODIPY-thapsigargin (ThermoFisher Scientific).

### Cell culture

Human embryonic kidney cell line HEK293FT and breast cancer cell line MCF-7 were both cultivated in Roswell Park Memorial Institute medium (RPMI, Thermo Fisher Scientific, Gibco) supplemented with 10% FCS (fetal calf serum, Merck, Darmstadt, Germany) and 1% penicillin/streptomycin (Thermo Fisher Scientific, Gibco). Prostate carcinoma cell line DU145 was cultivated in Dulbecco’s Modified Eagle’s medium (DMEM, Thermo Fisher Scientific, Gibco) supplemented with 10% FCS and 1% Penicillin/Streptomycin. All cell lines were maintained at 37°C and 5% CO2 and regularly authenticated via STR-profiling (Eurofins Genomics Germany, Ebersberg, Germany). For harvesting and seeding, cells were detached using 0.05% Trypsin-EDTA (Thermo Fisher Scientific, Gibco) and processed as described.

### Transgene expression

Cells were transfected with the indicated plasmids using PEI (polyethylenimine hydrochloride (PEI MAX 40K), Polysciences, Warrington, PA, USA) according to the manufacturer’s protocol. Samples used for microscopy were supplemented with 10 µM pan-caspase inhibitor Q-VD-OPh (Sellekchem, Houston, TX, USA). If not stated otherwise, transfected cells were kept at 37 °C and 5% CO2 for 18 h and subsequently harvested for indicated experiments.

### siRNA-mediated knock-down

Knock-down experiments were performed as described previously (66). Briefly, cells were seeded 24 h prior transfection and then transfected with ON-TARGET Plus Smartpool siRNA targeting BCL2 (siBCL-2) or non-targeted (siCTRL, Horizon Discovery, Waterbeach, UK) using DharmaFECT 1 reagent (Horizon Discovery) according to the manufacturer’s protocol. 24 h after knock-down cells were transfected with expression vectors encoding for EGFP or EGFP-BOK and incubated for another 18 h followed by flow cytometric analysis. For validation of knock-down efficiency cell lysates of transfected cells were analysed by Western Blot.

### Generation of vectors for transgene expression

Plasmids encoding for NanoBiT TMD fusion proteins were cloned using the NEBuilder system (New England Biolabs, Ipswich, MA, USA). Promega NanoBiT plasmid backbone (pBiT_1.1-C[TK/LgBiT], Promega, Madison, WI, USA) was combined with overlapping fragments generated by PCR which encoded for i) either mCitrine or mTurquoise2, ii) T2A sequence iii) LgBiT or SmBiT connected by a hydrophilic linker with iv) TMD sequences (TMD and primer sequences in Table S1, 2). T2A-SmBiT-TOM5-TMD fragment was generated via synthesis (GeneArt, Thermo Fisher Scientifc). Plasmids encoding for fluorescent TMD probes were cloned likewise combining the pEGFP-C1 backbone (Clontech, Kyoto, Japan) with overlapping fragments generated by PCR which encoded for mTurquoise2 and TMD sequences from the previously generated NanoBiT vectors (primer sequences in Table S3). Plasmids encoding for TMD chimeras were cloned likewise combining Fluorophore-fused protein-encoding plasmids containing protein cores (BOK/BCL-2) described previously (Einsele-Scholz et al. 2016) with overlapping fragments generated by PCR (Table S4) which encoded for TMD sequences (BAX/cb5/TOM5). Plasmid inserts were verified by sequencing.

### Flow cytometry

Cells and supernatant were harvested, cells were pelleted and washed in ice-cold PBS. Cell pellets were resuspended in 300 µl FACS PBS (PBS + 2% FCS) and if cells were not stained otherwise fluorescence was analyzed using a FACS Lyric flow cytometer (Becton Dickinson, Franklin Lakes, NJ, USA). Annexin-V-APC staining: After washing in PBS, cells were resuspended in 300 µl recombinant chicken Annexin-V-APC (ImmunoTools, Friesoythe, Germany) diluted 1:200 in Annexin-V-binding buffer (PBS, 2.5 mM CaCl2). Samples were incubated on ice for 10 min and analyzed using FACS Lyric (BD).

### Caspase 3/7 activity

Cells were seeded in 12-well plates (100,000 cells/well) and incubated at 37°C over night. On the next day, cells were transfected with indicated plasmids and incubated for another 18h. Then, cells were trypsinized and collected together with floating cells in supernatant medium. After centrifugation (500xg, 5 min), cells were washed with PBS once and distributed to three wells of a white-bottom 96-well plate (Thermo Fisher Scientific). Caspase-Glo 3/7 luciferase substrate (Promega) was added in a 1:1 ratio to each well followed by gentle shaking for 30 s. After 30 min incubation at RT, luminescence was detected with a Victor Nivo multimode plate reader (PerkinElmer, Waltham, MA, USA).

### Western Blot

Western Blot was done as described previously (Weller et al. 2022). In short, cells were harvested and cell pellets were resuspended in lysis buffer (50 mM Tris-HCl pH 7.6, 250 mM NaCl, 0.1% Triton X-100, 5 mM EDTA) supplemented with protease and phosphatase inhibitor cocktails. After sonification and clearance by centrifugation (15 min, 14000xg, 4°C) relative protein content was analyzed and cell lysates were diluted and incubated in Laemmli buffer (5 min, 95°C). Samples were then separated by SDS-PAGE and semi-dry blotted onto nitrocellulose membrane. After Blocking with 5% skim milk powder in TBS-T (TBS, 0.1% Tween-20) for 1 h, primary antibodies were applied in 5% BSA or 5% skim milk powder in TBS-T and incubated over night at 4°C. Blots were washed thrice in TBS-T for 10 min and afterwards HRP-coupled secondary antibodies diluted 1:2000 in 5% skim milk powder were applied for 1 h. TBS-T wash was repeated thrice for 10 min, ECL solution (SuperSignal West Dura, Thermo Fisher Scientific) was applied for 5 min and bands were detected using a STELLA imaging system (Raytest Isotopenmessgeräte GmbH, Straubenhardt, Germany).

### Generation of BMK/DKO cell lines

Baby mouse kidney BAX-/-/BAK-/- double knockout cells (BMK/DKO) (Degenhardt et al. 2002) were transduced with lentiviral particles (pLVX-EF1a-puro) for the expression of mTurquoise2-fused TMDs or were transfected to express the full-length mCerulean3-fused BIK as described previously (Osterlund et al. 2023). Individual clones with detectable expression were isolated by flow cytometry and clonal cell lines with similar fluorescence intensity were selected.

### Confocal laser scanning microscopy (cLSM)

Cells were grown on coverslips (#1, Paul Marienfeld. Lauda-Königshofen, Germany), transfected with indicated plasmids and subsequently fixed with 4% paraformaldehyde solution (ROTI Histofix, Carl Roth) for 20 min at 37°C. Then, cells were washed thrice with PBS and coverslips were mounted on microscopy slides (VWR, Radnor, PA, USA) with either DAKO fluorescent mounting medium (Agilent, Santa Clara, CA, USA) or ProLong Diamond antifade mountant with DAPI (Thermo Fisher Scientific).

#### Mitrotracker staining

Cells were stained prior to fixation in 100 nM of Mitotracker Red CMXRos (Thermo Fisher Scientific) in unsupplemented culture medium for 30 min at 37°C. Afterwards, cells were washed once in unsupplemented culture medium.

Images were acquired as z-stacks with a Leica TCS SP8 confocal laser-scanning microscope (Leica, Wetzlar, Germany) equipped with a HC PL APO CS2 63x/1.40 oil immersion objective and excitation lasers at 405 nm, 488 nm, 552 nm and 638 nm. Appropriate slit settings were used for detection of DAPI (415 - 480 nm), mTurquoise2 (415 – 480 nm), GFP (500 – 540 nm), mCitrine (500 – 540 nm), mCherry (570 - 700 nm) and mCarmine (650 – 770 nm) and specific fluorescence signals were acquired sequentially. Images were acquired and exported using the Leica Application Suite X software and further processed in Fiji software. For co-localization analysis, middle slices of cells were imaged, exported and analyzed using the Coloc2 plug-in of the Fiji software to yield Pearson’s correlation coefficients (Pearson’s r) of analyzed channels.

### Co-localization analysis in BMK/DKO cells

BMK/DKO cells expressing mTurquoise2-conjugated effector TMDs or mCerulean3-BIK were seeded in a 384-well plate (2000 cells/well). 24 hours later, these cells were stained with Draq5, Mitotracker red and BODIPY-Thapsigargin for 20 minutes at 37°C followed by imaging. Images were taken using a confocal Opera Phenix High-Content Screening System with a 63x water-immersion Objective (Perkin Elmer/revvity, Waltham, MA, USA) and a total of 1000 cells was randomly chosen for subsequent analysis. Pearson’s correlation coefficients between mTurquoise2/mCerulean3 and Mitotracker red or ER-marker BODIPY-Thapsigargin were determined in cytoplasmic regions of individual cells by excluding the Draq5-stained nuclei.

### NanoBiT luminescence assay

Indicated combinations of SmBiT- and LgBiT-TMD fusion proteins were expressed in cells by transient transfection of plasmids in a 1:1 ratio and incubation for 24 h. Cells were harvested, washed in PBS and resuspended in 150 µl Opti-MEM Reduced Serum Medium (Thermo Fisher Scientific). Luminescence and fluorescence were assessed in technical triplicates (50 µl cell suspension/well) in a white F-bottom 96-well plate (Greiner Bio-One, Frickenhausen, Germany) using an EnSpire multimode plate reader (PerkinElmer, Waltham, MA, USA). To each well 50 µl luciferase substrate coelenterazine-h (Promega) in Opti-MEM was added to obtain a final concentration of 5 µM and luminescence as well as fluorescence were detected immediately afterwards. Initially, fluorescence was assessed once for mTurquoise2 (excitation at 434 nm / detection at 474 nm) and mCitrine (ex. 516 nm / detection at 531 nm). Then, luminescence intensity was detected every 5 min for at least 30 min after substrate addition with a measurement duration of 1 s/well. For comparison of different samples, luminescence intensities 30 min after substrate addition were normalized to detected mTurquoise2 fluorescence. Normalized luminescence values for each technical triplicate were combined to a mean for each independent experiment.

### High-throughput multiscaling molecular dynamics simulations

Spontaneous self-association of BOK and BCL-2 TMDs (BOK-TMD comprises residues 180-S YNPGL RSHWL VAALC SFGRF LKAAF FVLLP ER-212, and BCL-2-TMD residues 208-PLF DFSWL SLKTL LSLAL VGACI TLGAY LGHK-239) was studied by DAFT (Wassenaar et al. 2015) using the Martini3 (Souza et al. 2021) coarse-grained force-field in a membrane mimic of the ER membrane. The model consisted of 75% 1-myristoyl-2-oleoyl-sn-glycero-3-phosphocholine (MOPC), 7% 1-myristoyl-2-oleoyl-sn-glycero-3-phosphoethanolamine (MOPE), 7% 1-palmitoyl-2-oleoyl-sn-glycero-3-phosphoinositol (POPI), 4% a-palmitoyl-2-oleoyl-sn-glycerol (PODG) and 7% cholesterol distributed symmetrically in the two membrane leaflets. After spontaneous association at CG resolution, the oligomers were clustered and representatives of the most often occurring conformations were converted back to atomistic resolution using backward (Wassenaar et al. 2014) and reequilibrated atomistically for 1µs using the CHARMM36m (Huang et al. 2017) force field and the TIP4p water model (Jorgensen und Madura 1985). All simulations were performed and analyzed using GROMACS 2020 (Abraham et al. 2015) and visualized in PyMOL (Schrödinger 2023). Gnuplot (Williams und Kelley 2013) was used to generate contact maps and secondary structure plots. Overview of performed simulations in Table S5.

### Statistical analysis

Statistical significance of differences was calculated by an unpaired t-test with Welch’s correction or one-way ANOVA with Tukey’s multiple comparison test as indicated using GraphPad Prism 9 software.

## Acknowledgments

Thanks go to Oliver Griesbeck and Arne Frabritius for kindly providing the mCarmine subcellular marker library. MD simulations were performed on the HoreKa supercomputer funded by the Ministry of Science, Research and the Arts Baden-Württemberg and by the Federal Ministry of Education and Research. This work was supported by the Deutsche Forschungsgemeinschaft under Germany’s Excellence Strategy – EXC 2075 – 390740016, the Stuttgart Center for Simulation Science (SC SimTech) and the Robert Bosch Stiftung GmbH.

## References

Abraham MJ, Murtola T, Schulz R, Páll S, Smith JC, Hess B, Lindahl E (2015) GROMACS: High performance molecular simulations through multi-level parallelism from laptops to supercomputers. SoftwareX: 19–25.

Andreu-Fernández V, Sancho M, Genovés A, Lucendo E, Todt F, Lauterwasser J, Funk K, Jahreis G, Pérez-Payá E, Mingarro I et al. (2017) Bax transmembrane domain interacts with prosurvival Bcl-2 proteins in biological membranes. Proceedings of the National Academy of Sciences of the United States of America. 2: 310–315.

Banjara S, Suraweera CD, Hinds MG, Kvansakul M (2020) The Bcl-2 Family: Ancient Origins, Conserved Structures, and Divergent Mechanisms. Biomolecules. 1:

Brito GC, Schormann W, Gidda SK, Mullen RT, Andrews DW (2019) Genome-wide analysis of Homo sapiens, Arabidopsis thaliana, and Saccharomyces cerevisiae reveals novel attributes of tail-anchored membrane proteins. BMC genomics. 1: 835.

Carpio MA, Means RE, Brill AL, Sainz A, Ehrlich BE, Katz SG (2021) BOK controls apoptosis by Ca2+ transfer through ER-mitochondrial contact sites. Cell reports. 10: 108827.

Carpio MA, Michaud M, Zhou W, Fisher JK, Walensky LD, Katz SG (2015) BCL-2 family member BOK promotes apoptosis in response to endoplasmic reticulum stress. Proceedings of the National Academy of Sciences of the United States of America. 23: 7201–7206.

Chang M-J, Zhong F, Lavik AR, Parys JB, Berridge MJ, Distelhorst CW (2014) Feedback regulation mediated by Bcl-2 and DARPP-32 regulates inositol 1,4,5-trisphosphate receptor phosphorylation and promotes cell survival. Proceedings of the National Academy of Sciences of the United States of America. 3: 1186–1191.

Chen H-C, Kanai M, Inoue-Yamauchi A, Tu H-C, Huang Y, Ren D, Kim H, Takeda S, Reyna DE, Chan PM et al. (2015) An interconnected hierarchical model of cell death regulation by the BCL-2 family. Nature cell biology. 10: 1270–1281.

Chi X, Nguyen D, Pemberton JM, Osterlund EJ, Liu Q, Brahmbhatt H, Zhang Z, Lin J, Leber B, Andrews DW (2020) The carboxyl-terminal sequence of bim enables bax activation and killing of unprimed cells. eLife.

Curran AR&Engelman DM (2003) Sequence motifs, polar interactions and conformational changes in helical membrane proteins. Current opinion in structural biology. 4: 412–417.

Deeks ED (2016) Venetoclax: First Global Approval. Drugs. 9: 979–987.

Degenhardt K, Sundararajan R, Lindsten T, Thompson C, White E (2002) Bax and Bak independently promote cytochrome C release from mitochondria. The Journal of biological chemistry. 16: 14127–14134.

Del Gaizo Moore V, Brown JR, Certo M, Love TM, Novina CD, Letai A (2007) Chronic lymphocytic leukemia requires BCL2 to sequester prodeath BIM, explaining sensitivity to BCL2 antagonist ABT-737. The Journal of clinical investigation. 1: 112–121.

DiNardo CD, Pratz K, Pullarkat V, Jonas BA, Arellano M, Becker PS, Frankfurt O, Konopleva M, Wei AH, Kantarjian HM et al. (2019) Venetoclax combined with decitabine or azacitidine in treatment-naive, elderly patients with acute myeloid leukemia. Blood. 1: 7–17.

Ding J, Zhang Z, Roberts GJ, Falcone M, Miao Y, Shao Y, Zhang XC, Andrews DW, Lin J (2010) Bcl-2 and Bax interact via the BH1-3 groove-BH3 motif interface and a novel interface involving the BH4 motif. The Journal of biological chemistry. 37: 28749–28763.

Echeverry N, Bachmann D, Ke F, Strasser A, Simon HU, Kaufmann T (2013) Intracellular localization of the BCL-2 family member BOK and functional implications. Cell death and differentiation. 6: 785–799.

Egan B, Beilharz T, George R, Isenmann S, Gratzer S, Wattenberg B, Lithgow T (1999) Targeting of tail-anchored proteins to yeast mitochondria in vivo. FEBS letters. 3: 243–248.

Einsele-Scholz S, Malmsheimer S, Bertram K, Stehle D, Johänning J, Manz M, Daniel PT, Gillissen BF, Schulze-Osthoff K, Essmann F (2016) Bok is a genuine multi-BH-domain protein that triggers apoptosis in the absence of Bax and Bak. Journal of cell science. 15: 3054.

England CG, Ehlerding EB, Cai W (2016) NanoLuc: A Small Luciferase Is Brightening Up the Field of Bioluminescence. Bioconjugate chemistry. 5: 1175–1187.

Fabritius A, Ng D, Kist AM, Erdogan M, Portugues R, Griesbeck O (2018) Imaging-Based Screening Platform Assists Protein Engineering. Cell chemical biology. 12: 1554–1561.e8.

Fernández-Marrero Y, Bleicken S, Das KK, Bachmann D, Kaufmann T, Garcia-Saez AJ (2017) The membrane activity of BOK involves formation of large, stable toroidal pores and is promoted by cBID. The FEBS journal. 5: 711–724.

Fresquet V, Rieger M, Carolis C, García-Barchino MJ, Martinez-Climent JA (2014) Acquired mutations in BCL2 family proteins conferring resistance to the BH3 mimetic ABT-199 in lymphoma. Blood. 26: 4111–4119.

Friess MD, Pluhackova K, Böckmann RA (2018) Structural Model of the mIgM B-Cell Receptor Transmembrane Domain From Self-Association Molecular Dynamics Simulations. Frontiers in immunology: 2947.

Gardai SJ, Hildeman DA, Frankel SK, Whitlock BB, Frasch SC, Borregaard N, Marrack P, Bratton DL, Henson PM (2004) Phosphorylation of Bax Ser184 by Akt regulates its activity and apoptosis in neutrophils. The Journal of biological chemistry. 20: 21085–21095.

Gössweiner-Mohr N, Siligan C, Pluhackova K, Umlandt L, Koefler S, Trajkovska N, Horner A (2022) The Hidden Intricacies of Aquaporins: Remarkable Details in a Common Structural Scaffold. Small (Weinheim an der Bergstrasse, Germany). 31: e2202056.

Guedes RP, Rocha E, Mahiou J, Moll HP, Arvelo MB, Taube JM, Peterson CR, Kaczmarek E, Longo CR, Da Silva CG et al. (2013) The C-terminal domain of A1/Bfl-1 regulates its anti-inflammatory function in human endothelial cells. Biochimica et biophysica acta. 6: 1553– 1561.

Han J, Pluhackova K, Böckmann RA (2016) Exploring the Formation and the Structure of Synaptobrevin Oligomers in a Model Membrane. Biophysical journal. 9: 2004–2015.

Horie C, Suzuki H, Sakaguchi M, Mihara K (2002) Characterization of signal that directs C-tail-anchored proteins to mammalian mitochondrial outer membrane. Molecular Biology of the Cell. 5: 1615–1625.

Hsu SY, Kaipia A, McGee E, Lomeli M, Hsueh AJ (1997) Bok is a pro-apoptotic Bcl-2 protein with restricted expression in reproductive tissues and heterodimerizes with selective anti-apoptotic Bcl-2 family members. Proceedings of the National Academy of Sciences of the United States of America. 23: 12401–12406.

Huang J, Rauscher S, Nawrocki G, Ran T, Feig M, Groot BL de, Grubmüller H, MacKerell AD (2017) CHARMM36m: an improved force field for folded and intrinsically disordered proteins. Nature methods. 1: 71–73.

Irving JA, Whisstock JC, Lesk AM (2001) Protein structural alignments and functional genomics. Proteins. 3: 378–382.

Ivanova H, Ritaine A, Wagner L, Luyten T, Shapovalov G, Welkenhuyzen K, Seitaj B, Monaco G, Smedt H de, Prevarskaya N, et al. (2016) The trans-membrane domain of Bcl-2α, but not its hydrophobic cleft, is a critical determinant for efficient IP3 receptor inhibition. Oncotarget. 34: 55704–55720.

Jeong S-Y, Gaume B, Lee Y-J, Hsu Y-T, Ryu S-W, Yoon S-H, Youle RJ (2004) Bcl-x(L) sequesters its C-terminal membrane anchor in soluble, cytosolic homodimers. The EMBO journal. 10: 2146–2155.

Jorgensen WL&Madura JD (1985) Temperature and size dependence for Monte Carlo simulations of TIP4P water. Molecular Physics. 6: 1381–1392.

Kale J, Kutuk O, Brito GC, Andrews TS, Leber B, Letai A, Andrews DW (2018a) Phosphorylation switches Bax from promoting to inhibiting apoptosis thereby increasing drug resistance. EMBO Reports. 9:

Kale J, Osterlund EJ, Andrews DW (2018b) BCL-2 family proteins: changing partners in the dance towards death. Cell death and differentiation. 1: 65–80.

Kaufmann T, Schlipf S, Sanz J, Neubert K, Stein R, Borner C (2003) Characterization of the signal that directs Bcl-x(L), but not Bcl-2, to the mitochondrial outer membrane. The Journal of Cell Biology. 1: 53–64.

Kim S, Jeon T-J, Oberai A, Yang D, Schmidt JJ, Bowie JU (2005) Transmembrane glycine zippers: physiological and pathological roles in membrane proteins. Proceedings of the National Academy of Sciences of the United States of America. 40: 14278–14283.

Korn V&Pluhackova K (2022) Not sorcery after all: Roles of multiple charged residues in membrane insertion of gasdermin-A3. Frontiers in Cell and Developmental Biology: 958957.

Korsmeyer SJ, Shutter JR, Veis DJ, Merry DE, Oltvai ZN (1993) Bcl-2/Bax: a rheostat that regulates an anti-oxidant pathway and cell death. Seminars in cancer biology. 6: 327–332.

Kwon M-J, Park J, Jang S, Eom C-Y, Oh E-S (2016) The Conserved Phenylalanine in the Transmembrane Domain Enhances Heteromeric Interactions of Syndecans. The Journal of biological chemistry. 2: 872–881.

Lai Y-C, Li C-C, Sung T-C, Chang C-W, Lan Y-J, Chiang Y-W (2019) The role of cardiolipin in promoting the membrane pore-forming activity of BAX oligomers. Biochimica et biophysica acta. Biomembranes. 1: 268–280.

Lalier L, Mignard V, Joalland M-P, Lanoé D, Cartron P-F, Manon S, Vallette FM (2021) TOM20-mediated transfer of Bcl2 from ER to MAM and mitochondria upon induction of apoptosis. Cell Death & Disease. 2: 182.

Leber B, Lin J, Andrews DW (2007) Embedded together: the life and death consequences of interaction of the Bcl-2 family with membranes. Apoptosis : an international journal on programmed cell death. 5: 897–911.

Letai A, Bassik MC, Walensky LD, Sorcinelli MD, Weiler S, Korsmeyer SJ (2002) Distinct BH3 domains either sensitize or activate mitochondrial apoptosis, serving as prototype cancer therapeutics. Cancer cell. 3: 183–192.

Liu Q, Osterlund EJ, Chi X, Pogmore J, Leber B, Andrews DW (2019) Bim escapes displacement by BH3-mimetic anti-cancer drugs by double-bolt locking both Bcl-XL and Bcl-2. eLife.

Llambi F, Moldoveanu T, Tait SWG, Bouchier-Hayes L, Temirov J, McCormick LL, Dillon CP, Green DR (2011) A unified model of mammalian BCL-2 protein family interactions at the mitochondria. Molecular cell. 4: 517–531.

Llambi F, Wang Y-M, Victor B, Yang M, Schneider DM, Gingras S, Parsons MJ, Zheng JH, Brown SA, Pelletier S et al. (2016) BOK Is a Non-canonical BCL-2 Family Effector of Apoptosis Regulated by ER-Associated Degradation. Cell. 2: 421–433.

Lucendo E, Sancho M, Lolicato F, Javanainen M, Kulig W, Leiva D, Duart G, Andreu-Fernández V, Mingarro I, Orzáez M (2020) Mcl-1 and Bok transmembrane domains: Unexpected players in the modulation of apoptosis. Proceedings of the National Academy of Sciences of the United States of America. 45: 27980–27988.

Lucken-Ardjomande S, Montessuit S, Martinou J-C (2008) Contributions to Bax insertion and oligomerization of lipids of the mitochondrial outer membrane. Cell death and differentiation. 5: 929–937.

Lutter M, Fang M, Luo X, Nishijima M, Xie X, Wang X (2000) Cardiolipin provides specificity for targeting of tBid to mitochondria. Nature cell biology. 10: 754–761.

Monaco G, Decrock E, Akl H, Ponsaerts R, Vervliet T, Luyten T, Maeyer M de, Missiaen L, Distelhorst CW, Smedt H de, et al. (2012) Selective regulation of IP3-receptor-mediated Ca2+ signaling and apoptosis by the BH4 domain of Bcl-2 versus Bcl-Xl. Cell death and differentiation. 2: 295–309.

Nechushtan A, Smith CL, Hsu YT, Youle RJ (1999) Conformation of the Bax C-terminus regulates subcellular location and cell death. The EMBO journal. 9: 2330–2341.

Osterlund EJ, Hirmiz N, Nguyen D, Pemberton JM, Fang Q, Andrews DW (2023) Endoplasmic reticulum protein BIK binds to and inhibits mitochondria-localized antiapoptotic proteins. The Journal of biological chemistry. 2: 102863.

Osterlund EJ, Hirmiz N, Pemberton JM, Nougarède A, Liu Q, Leber B, Fang Q, Andrews DW (2022) Efficacy and specificity of inhibitors of BCL-2 family protein interactions assessed by affinity measurements in live cells. Science advances. 16: eabm7375.

Pemberton JM, Nguyen D, Osterlund EJ, Schormann W, Pogmore JP, Hirmiz N, Leber B, Andrews DW (2023) The carboxyl-terminal sequence of PUMA binds to both anti-apoptotic proteins and membranes. eLife.

Pluhackova K, Gahbauer S, Kranz F, Wassenaar TA, Böckmann RA (2016) Dynamic Cholesterol-Conditioned Dimerization of the G Protein Coupled Chemokine Receptor Type 4. PLoS computational biology. 11: e1005169.

Pluhackova K&Horner A (2021) Native-like membrane models of E. coli polar lipid extract shed light on the importance of lipid composition complexity. BMC biology. 1: 4.

Rabe M, Aisenbrey C, Pluhackova K, Wert V de, Boyle AL, Bruggeman DF, Kirsch SA, Böckmann RA, Kros A, Raap J, et al. (2016) A Coiled-Coil Peptide Shaping Lipid Bilayers upon Fusion. Biophysical journal. 10: 2162–2175.

Roberts AW, Davids MS, Pagel JM, Kahl BS, Puvvada SD, Gerecitano JF, Kipps TJ, Anderson MA, Brown JR, Gressick L et al. (2016) Targeting BCL2 with Venetoclax in Relapsed Chronic Lymphocytic Leukemia. The New England journal of medicine. 4: 311– 322.

Robertson LE, Plunkett W, McConnell K, Keating MJ, McDonnell TJ (1996) Bcl-2 expression in chronic lymphocytic leukemia and its correlation with the induction of apoptosis and clinical outcome. Leukemia. 3: 456–459.

Roca-Portoles A, Rodriguez-Blanco G, Sumpton D, Cloix C, Mullin M, Mackay GM, O’Neill K, Lemgruber L, Luo X, Tait SWG (2020) Venetoclax causes metabolic reprogramming independent of BCL-2 inhibition. Cell Death & Disease. 8: 616.

Rong Y-P, Bultynck G, Aromolaran AS, Zhong F, Parys JB, Smedt H de, Mignery GA, Roderick HL, Bootman MD, Distelhorst CW (2009) The BH4 domain of Bcl-2 inhibits ER calcium release and apoptosis by binding the regulatory and coupling domain of the IP3 receptor. Proceedings of the National Academy of Sciences of the United States of America. 34: 14397–14402.

Russ WP&Engelman DM (2000) The GxxxG motif: a framework for transmembrane helix-helix association. Journal of molecular biology. 3: 911–919.

Schinzel A, Kaufmann T, Borner C (2004) Bcl-2 family members: integrators of survival and death signals in physiology and pathology corrected. Biochimica et biophysica acta. 2-3: 95–105.

Schrödinger L (2023) The PyMOL Molecular Graphics System, Version 2.0. pymol.org.

Schulman JJ, Szczesniak LM, Bunker EN, Nelson HA, Roe MW, Wagner LE, Yule DI, Wojcikiewicz RJH (2019) Bok regulates mitochondrial fusion and morphology. Cell death and differentiation. 12: 2682–2694.

Schulman JJ, Wright FA, Han X, Zluhan EJ, Szczesniak LM, Wojcikiewicz RJH (2016) The Stability and Expression Level of Bok Are Governed by Binding to Inositol 1,4,5-Trisphosphate Receptors. The Journal of biological chemistry. 22: 11820–11828.

Senes A, Gerstein M, Engelman DM (2000) Statistical analysis of amino acid patterns in transmembrane helices: the GxxxG motif occurs frequently and in association with beta-branched residues at neighboring positions. Journal of molecular biology. 3: 921–936.

Shalaby R, Diwan A, Flores-Romero H, Hertlein V, Garcia-Saez AJ (2023) Visualization of BOK pores independent of BAX and BAK reveals a similar mechanism with differing regulation. Cell death and differentiation. 3: 731–741.

Shamas-Din A, Bindner S, Chi X, Leber B, Andrews DW, Fradin C (2015) Distinct lipid effects on tBid and Bim activation of membrane permeabilization by pro-apoptotic Bax. The Biochemical journal. 3: 495–505.

Simonyan L, Renault TT, Da Novais MJC, Sousa MJ, Côrte-Real M, Camougrand N, Gonzalez C, Manon S (2016) Regulation of Bax/mitochondria interaction by AKT. FEBS letters. 1: 13–21.

Souza PCT, Alessandri R, Barnoud J, Thallmair S, Faustino I, Grünewald F, Patmanidis I, Abdizadeh H, Bruininks BMH, Wassenaar TA et al. (2021) Martini 3: a general purpose force field for coarse-grained molecular dynamics. Nature methods. 4: 382–388.

Stehle D, Grimm M, Einsele-Scholz S, Ladwig F, Johänning J, Fischer G, Gillissen B, Schulze-Osthoff K, Essmann F (2018) Contribution of BH3-domain and Transmembrane-domain to the Activity and Interaction of the Pore-forming Bcl-2 Proteins Bok, Bak, and Bax. Scientific reports. 1: 12434.

Todt F, Cakir Z, Reichenbach F, Youle RJ, Edlich F (2013) The C-terminal helix of Bcl-x(L) mediates Bax retrotranslocation from the mitochondria. Cell death and differentiation. 2: 333– 342.

Unterreitmeier S, Fuchs A, Schäffler T, Heym RG, Frishman D, Langosch D (2007) Phenylalanine promotes interaction of transmembrane domains via GxxxG motifs. Journal of molecular biology. 3: 705–718.

Walter F, D’Orsi B, Jagannathan A, Dussmann H, Prehn JHM (2022) BOK controls ER proteostasis and physiological ER stress responses in neurons. Frontiers in Cell and Developmental Biology: 915065.

Wassenaar TA, Pluhackova K, Böckmann RA, Marrink SJ, Tieleman DP (2014) Going Backward: A Flexible Geometric Approach to Reverse Transformation from Coarse Grained to Atomistic Models. Journal of chemical theory and computation. 2: 676–690.

Wassenaar TA, Pluhackova K, Moussatova A, Sengupta D, Marrink SJ, Tieleman DP, Böckmann RA (2015) High-Throughput Simulations of Dimer and Trimer Assembly of Membrane Proteins. The DAFT Approach. Journal of chemical theory and computation. 5: 2278–2291.

Weller S, Toennießen A, Schaefer B, Beigl T, Muenchow A, Böpple K, Hofmann U, Gillissen BF, Aulitzky WE, Kopp H-G et al. (2022) The BCL-2 inhibitor ABT-199/venetoclax synergizes with proteasome inhibition via transactivation of the MCL-1 antagonist NOXA. Cell death discovery. 1: 215.

Williams T&Kelley C (2013) Gnuplot 4.6: an interactive plotting program. gnuplot.sourceforge.net.

Zhang Z, Subramaniam S, Kale J, Liao C, Huang B, Brahmbhatt H, Condon SGF, Lapolla SM, Hays FA, Ding J et al. (2016) BH3-in-groove dimerization initiates and helix 9 dimerization expands Bax pore assembly in membranes. The EMBO journal. 2: 208–236.

Zhu W, Cowie A, Wasfy GW, Penn LZ, Leber B, Andrews DW (1996) Bcl-2 mutants with restricted subcellular location reveal spatially distinct pathways for apoptosis in different cell types. The EMBO journal. 16: 4130–4141.

