## supplemental information for "BCL-2 and BOK regulate apoptosis by interaction of their C-terminal transmembrane domains"

**Running title**

BCL-2/BOK transmembrane domain interaction at ER

**Competing interests**

The authors declare to have no competing interests to disclose.

**This PDF file includes:**

Supporting information text
Figures S1 to S18
Tables S1 to S9
SI References

### Supporting information text

#### *High-throughput multiscaling molecular dynamics simulation*

In detail, TMDs of BOK and BCL-2 were modelled as α-helices in PyMOL (Schrödinger 2023) based on their amino-acid sequence and coarse-grained (CG) by martinize2 (https://github.com/marrink-lab/vermouth-martinize). The CG peptides were aligned along the z-axis, randomly rotated along the z-axis and placed in 6 nm relative distance. The peptides were then surrounded by a membrane (ER or MOM) and water with 0.1 M NaCl by insane (Wassenaar et al. 2015). The simulation systems were consecutively energy minimized by 5000 steps of the steepest descent algorithm. Then velocities, reflecting the Boltzmann distribution at 310 K, were generated and the systems were equilibrated to 310 K and to 1 bar in 5 ns MD simulation with the time step of 20 fs, v-rescale thermostat with time constant of 1 ps, Berendsen barostat (Berendsen et al. 1984) with time constant of 12 ps, semiisotropic pressure coupling and compressibility of 3e-4 1/bar. The van der Waals interactions were shifted to 0 at 1.1 nm by the Potential-shift-Verlet algorithm and the electrostatics for among charged particles in a distance larger than 1.1 nm was described by reaction field and the $\varepsilon r$ of 15. The Verlet (Páll und Hess 2013) cut-off scheme was used for particle-based cut-offs and the neighbor list was updated every 20 steps. The bond constraints were achieved by LINCS (Hess 2008) and the center of mass of the simulation system was linearly removed every 1000 steps. Dimerization was simulated for 10 µs, tetramerization was simulated 50 µs and Parrinello-Rahman barostat (Parrinello und Rahman 1980, 1981) was used instead of the Berendsen barostat. For the complete list of simulations, see Table S5.

After resolution conversion to atomistic resolutions, all atom simulations were initiated by a short simulation (1 ps with 0.2 fs timestep) with position restraints on the peptides in which the system was heated up to 310 K by the Berendsen thermostat. Afterwards, a 10 ns simulation with the timestep of 2 fs was performed and position restraints on the peptides´ backbone atoms only was done to equilibrate the membrane and solvent around the oligomers. The temperature was coupled to a bath of 310 K by the Nosé-Hoover thermostat (Evans und Holian 1985) and coupling constant of 0.5 ps and the pressure was controlled by the Berendsen barostat each 1 ps to be 1 bar. The following production run simulations lasted 1 µs using the time step of 2 fs. The temperature of 310 K was modulated by the Nosé-Hoover thermostat (Evans und Holian 1985) and coupling constant of 0.5 ps. The pressure of 1 bar was controlled in a semiisotropic manner by the Parrinello-Rahman barostat using the compressibility of 4.5e-5 and coupling constant 1 ps. While the van der Waals interactions were switched to zero between 0.8 and 1.2 nm, the electrostatics was treated by PME (Darden et al. 1993) beyond 1.2 nm. The neighbor list was updated each 10 steps by the Verlet cut-off scheme and the center of mass of the system was removed every 500 steps. The bonds to hydrogen atoms were constrained by LINCS (Hess 2008).

#### *ER membrane model*

The ER membrane model was based on recent mass spectrometric data (Scrima et al. 2022). Because the ER membrane is suggested to be thinner than other membranes (Prasad et al. 2020), short-chain lipid tails were assumed. For the ER membrane mimic, 75% 1-myristoyl-2-oleoyl-sn-glycero-3-phosphocholine (MOPC), 7% 1-myristoyl-2-oleoyl-sn-glycero-3-phosphoethanolamine (MOPE), 7% 1-palmitoyl-2-oleoyl-sn-glycero-3-phosphoinositol (POPI), 4% a-palmitoyl-2-oleoyl-sn-glycerol (PODG) and 7% cholesterol were distributed symmetrically in the two membrane leaflets.

#### *Analysis details*

Conformation stability of oligomers consisting of BOK-TMD and BCL-2-TMD was evaluated by estimation of RMSDs on per dimer basis as described in the following in order to reduce the dependency of RMSD on the protein/peptide assembly size. At first the peptides in each AA simulation were clustered to remove splitting of the oligomers over periodic boundary conditions. Next, trajectories of all pairs of dimers were extracted from each simulation and fitted onto the frame at 0 ns (corresponding to the CG conformation, (Irving et al. 2001)). Then, average RMSD over 100-1000 ns AA simulation time relative to 0 ns was estimated for each pair of peptides and averaged over a simulation (i.e. in trimers RMSD of three pairs of peptides were averaged per simulation and in tetramers the RMSD of six pairs of peptides were averaged.) The average per simulation RMSD values were then used to calculate the average and SEM per simulation set (i.e. BOK-TMD homodimer, BOK-TMD/BCL-2-TMD heterodimer, etc.). Peptide secondary structure was estimated by the dictionary of protein secondary structure (DSSP, (Kabsch und Sander 1983)) over 100-1000 ns AA simulation. Average interaction energies per residue of one peptide with all other peptides were extracted for each residue of each peptide over the last 100 ns of the AA simulations and averaged over all BOK-TMD or BCL-2-TMD. These average interaction energies per residue were averaged over all simulations from the simulation set (i.e. BOK-TMD homodimer, BOK-TMD/BCL-2-TMD heterodimer etc.) which enabled also the estimation of SEM. Contact maps were calculated using the GROMACS tool mdmat and the settings of 10 levels and truncate distance of 1.2 nm for 1 µs frames. Lipid binding per residue was calculated over 100-1000 ns of AA simulations. In the first step the number of lipids and their type (e.g. CHOL, POPI etc.) that were in contact (i.e. within 0.3 nm) to each residue of each peptide were estimated in each trajectory frame. In the next step the relative lipid composition in the surroundings of each residue in each trajectory frame was evaluated and compared to the lipid composition in the bilayer, i.e. cholesterol 0.07, MOPC 0.75 MOPE 0.07, PODG 0.04 and POPI 0.07. We define lipid depletion and enrichment as relative abundance that is smaller or larger, respectively, around the residue than in the total bilayer. The relative hidden protein surface was calculated for each trajectory frame of AA simulations as the ratio between the surface unaccessible solvent surface (SASA) of the oligomer (i.e. the difference between the sum of SASAs of all peptides and of the oligomer SASA) and the total SASA of all peptides. Standard settings of GROMACS tool sasa were used. Average relative hidden protein surface for each simulation from 100-1000 ns was then calculated and these averages per simulation used to calculate the average and SEM over each simulation set.

Gnuplot (Williams und Kelley 2013) was used to generate contact maps and secondary structure plots, R (R Core Team 2022) was used to make the interaction energy plot, all images of molecules were rendered in PyMOL (Schrödinger 2023).

**Supplementary Figures**

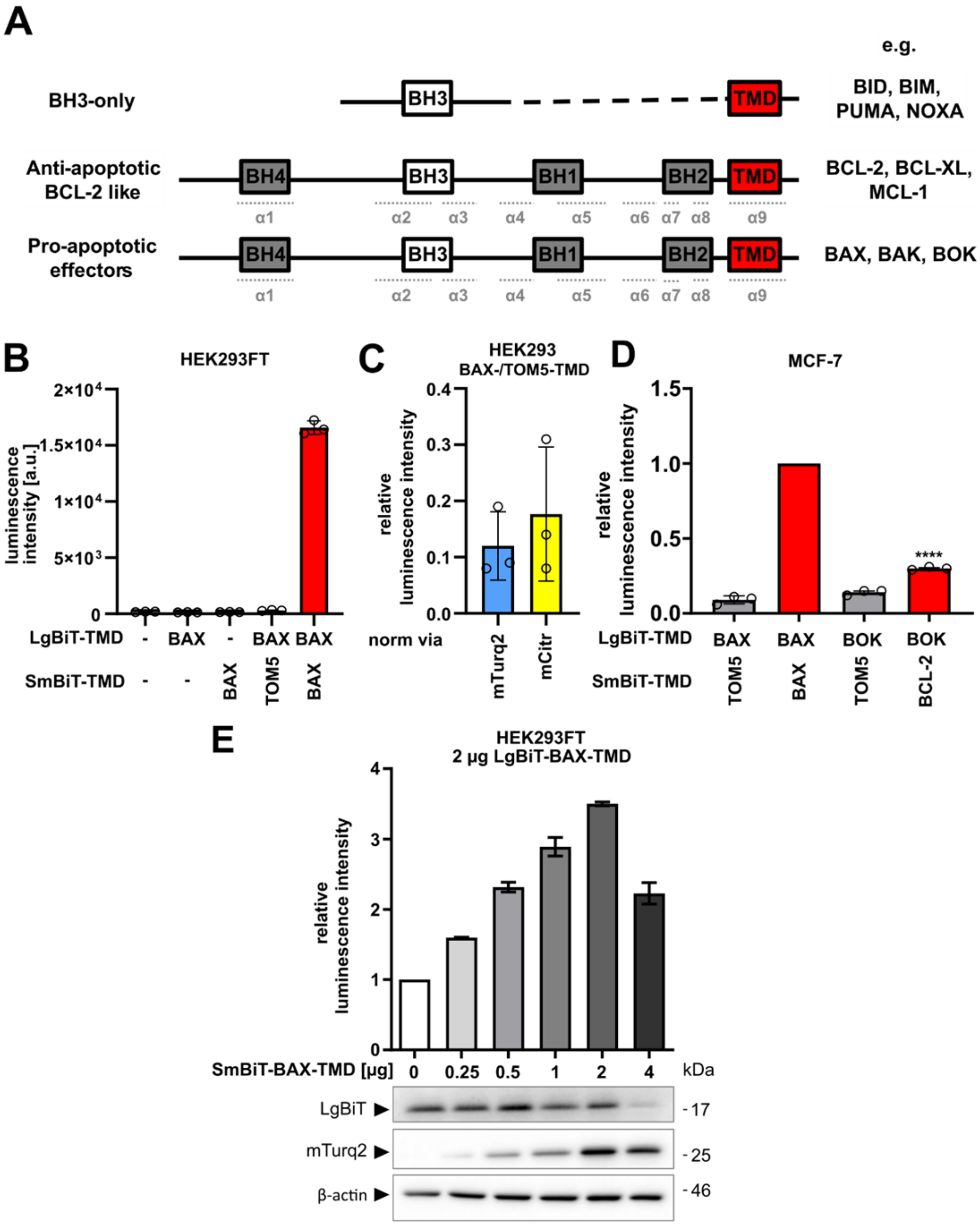

**Figure S1: NanoBiT split luciferase assay.**

**A** - Common structure of Bcl-2 family proteins. Shown are the BH (= Bcl-2 homology) motifs and TMD as boxes. Black dotted line indicates gaps. Position of alpha helices (α1-9) are indicated in dotted, light gray lines below.

**B** - HEK293FT cells were transfected with plasmids for the expression of LgBiT- and/or SmBiT-TMDs (or were left untransfected), harvested after 24 h and luminescence was detected in a multimode plate reader. Shown are luminescence intensites prior to data normalisation. Mean ± sd of technical triplicates from one representative experiment.

**C** - HEK293FT cells were co-transfected with plasmids for the expression of LgBiT-BAX-TMD and SmBiT-TOM5-TMD or SmBiT-BAX-TMD, harvested after 24 h and subjected to the split-luciferase assay. Detected luminescence was normalized to simultaneously acquired mTurquoise2 fluorescence (mTurq2) or mCitrine fluorescence (mCitr). Mean ± sd from three independent experiments shown in relation to simultaneously detected positive control (BAX-TMD/BAX-TMD, set to 1.00).

**D** - HEK293FT cells were transfected with increasing amounts of plasmid for the expression of SmBiT-BAX-TMD, while keeping co-transfected LgBiT-BAX-TMD-encoding plasmid constant. To achieve equal total amount of DNA per transfection, SmBiT-BAX-TMD was mixed with empty backbone vector. Cells were harvested 24 h post transfection and subjected to both split-luciferase assay as well as Western Blot. Luminescence intensities are shown as mean ± sd of technical triplicates from one representative experiment relative to transfection without SmBiT-BAX-TMD-encoding plasmid. Detection of LgBiT and mTurquoise2 (mTurq2) expression in same samples using Western Blot is shown below. β-actin was used as loading control.

**E** - Split-luciferase assay confirming BOK-TMD/BCL-2-TMD interaction in MCF-7 cells. MCF-7 cells were transfected with plasmids for the expression of indicated NanoBiT-TMD fusion proteins and harvested after 24 h to be subjected to NanoBiT assay. Luminescence intensities shown as mean ± sd from three independent experiment were set in relation to positive control (BAX-TMD/BAX-TMD).

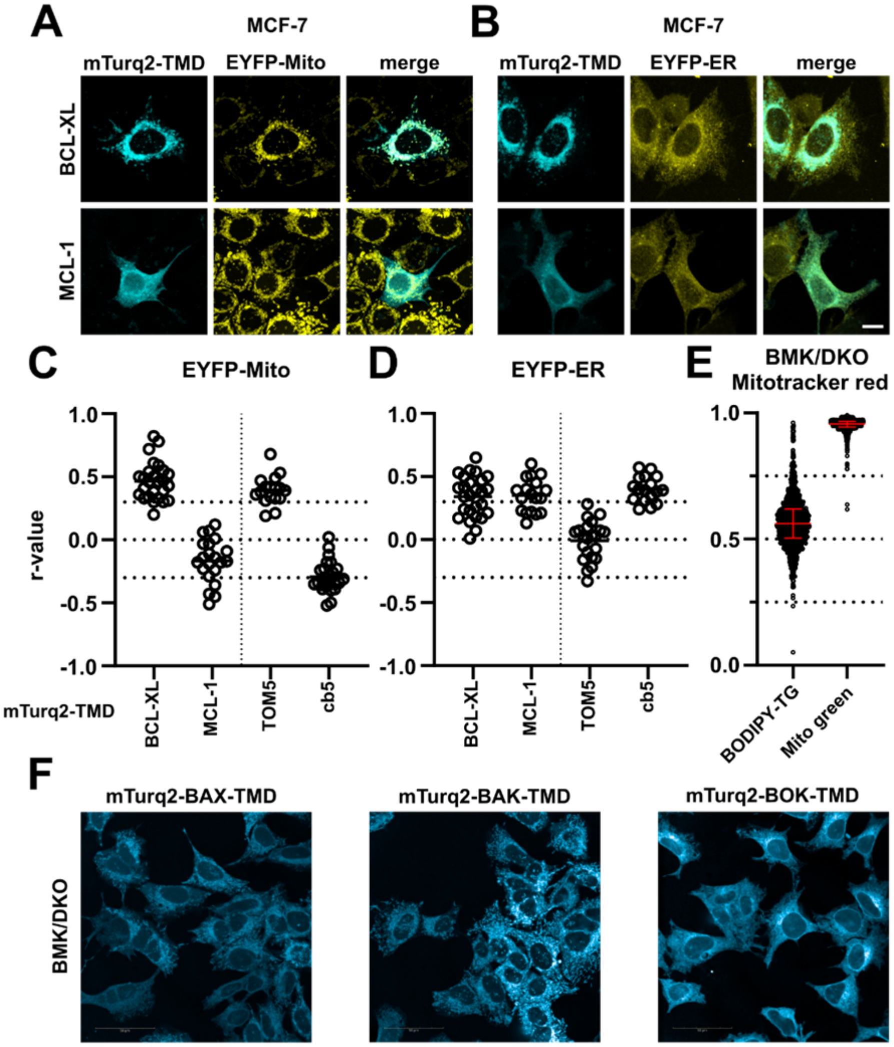
A
MCF-7
mTurq2-TMD
EYFP-Mito
merge
BCL-XL
MCL-1
B
MCF-7
mTurq2-TMD
EYFP-ER
merge
C
EYFP-Mito
r-value
1.0
0.5
0.0
-0.5
-1.0
mTurq2-TMD
BCL-XL
MCL-1
TOM5
cb5
D
EYFP-ER
1.0
0.5
0.0
-0.5
-1.0
BCL-XL
MCL-1
TOM5
cb5
E
BMK/DKO
Mitotracker red
1.0
0.5
0.0
BODIPY-TG
Mito green
F
mTurq2-BAX-TMD
mTurq2-BAK-TMD
mTurq2-BOK-TMD
BMK/DKO

**Figure S2: TMD peptide localization of MCL-1 and BCL-XL.**

**A, B** - Subcellular localization of MCL-1-TMD and BCL-XL-TMD. MCF-7 cells expressing (A) EYFP-Mito or (B) EYFP-ER were transfected with plasmids for the expression of mTurquoise2 (mTurq2)-labelled BCL-XL-TMD or MCL-1-TMD and 18 h post transfection cells were imaged by cLSM. Images are maximum projections of representative z-stacks from three independent experiments. Scale bar = 10 µm.

**C, D** - Quantitative analysis of mTurquoise2-fused BCL-XL-TMD or MCL-1-TMD co-localization with (C) EYFP-labelled mitochondria and (D) EYFP-labelled ER from cLSM images A, B. Graphs show Pearson's r correlation coefficient for a total of ≥ 15 cells combined from three independent experiments. The mean is marked as a horizontal line. Data for mTurquoise-TOM5-TMD and mTurquoise2-cb5-TMD are shown for comparison.

**E** - Co-localization analysis of subcellular markers in BMK/DKO cells. BMK/DKO cells labelled with DRAQ5 (nuclei) and Mitotracker red were stained with BODIPY-Thapsigargin (BODIPY-TG, ER control) or Mitotracker green (Mito control). Images of n ≥ 1000 cells were acquired using an Opera Phenix spinning disk system, images were analyzed and Pearson's r calculated. Shown is median + IQR.

**F** - Representative confocal spinning disk images of BMK/DKO cells expressing mTurquoise2-fused BAX-TMD, BAK-TMD or BOK-TMD.

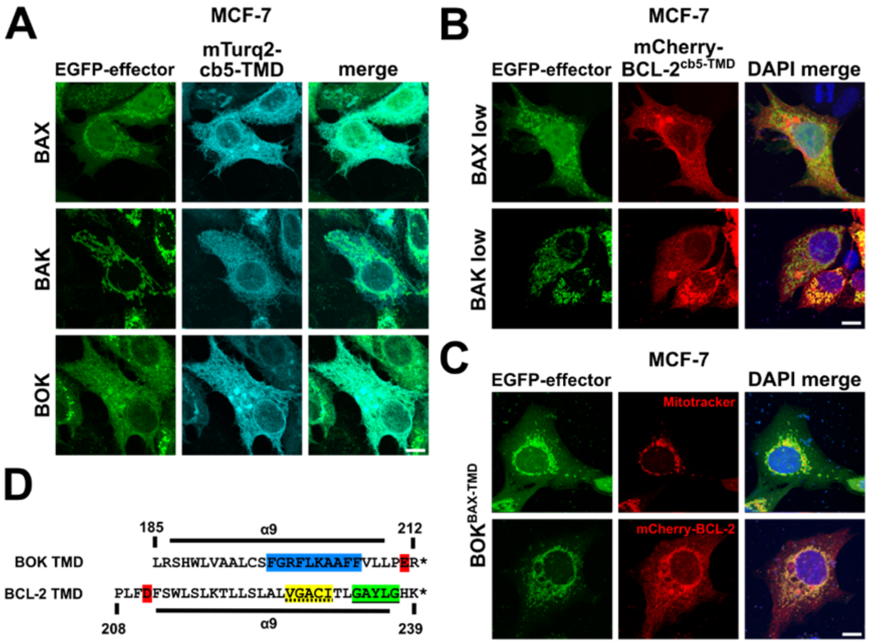


**Figure S3: Co-localization of BOK and BCL-2 at the ER is TMD dependent.**

**A** - Co-localization of BAX, BAK and BOK with ER. MCF-7 cells were co-transfected with plasmids for the expression of EGFP-BAX, EGFP-BAK or EGFP-BOK and mTurq2-cb5-TMD.

**B** - Co-localization of BAX and BAK (low expression) with BCL-2cb5-TMD. MCF-7 cells were co-transfected with plasmids for the expression of EGFP-BAX or EGFP-BAK and mCherry-BCL-2TOM5-TMD. Shown images represent cells without cluster formation of BAX or BAK.

**C** - Co-localization of BOKBAX-TMD with mitochondria and BCL-2. MCF-7 cells were transfected with plasmids for the expression of EGFP-BOKBAX-TMD and stained with Mitotracker red (upper panel) or co-transfected with plasmids for the expression of mCherry-BCL-2 (lower panel). (A. B, C) Cells were fixed after 24 h followed by cLSM. Images are maximum projections of z-stacks representative of two independent experiments. Scale bar = 10 µm.

**D** - Sequence of BOK-TMD and BCL-2-TMD showing interaction motifs GG4 (green), VI4 (yellow), and phenylalanine-rich region (blue). Negatively charged amino acids (Asp, Glu) are marked in red.

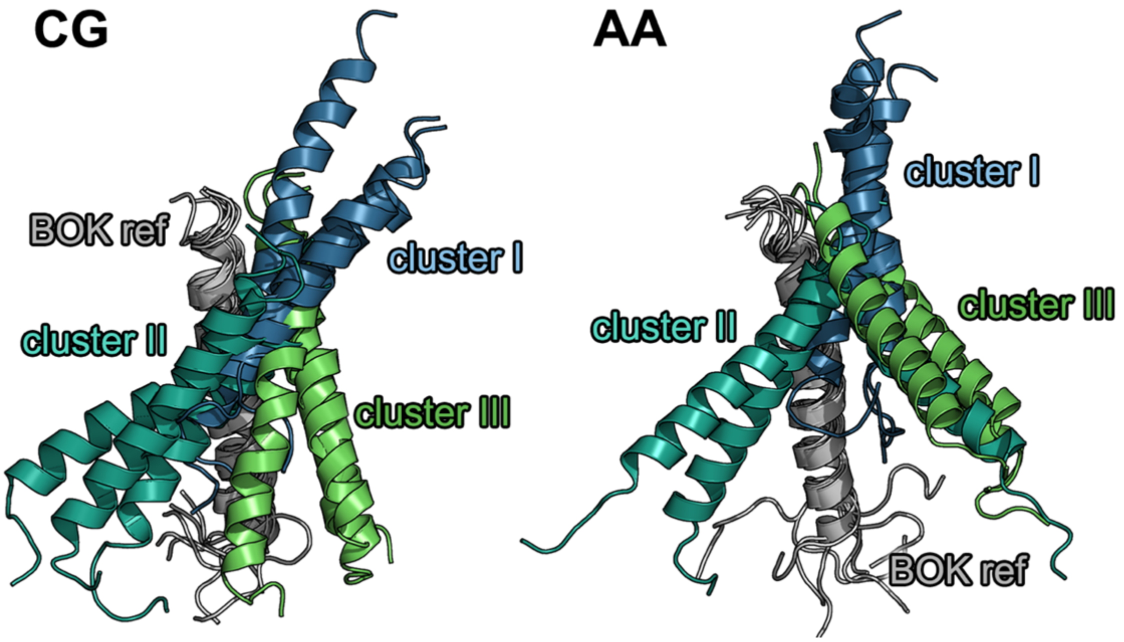


**Figure S4: BOK/BOK homodimers.**

Visualization of clusters of BOK/BOK homodimers after an overlay to an identical reference (BOK ref in grey). CG labels structures prior to AA simulations, as AA structures after 1µs AA simulation are shown. The nonoverlayed BOK TMDs in cluster I are colored blue, in cluster II aquamarine and in cluster III green.

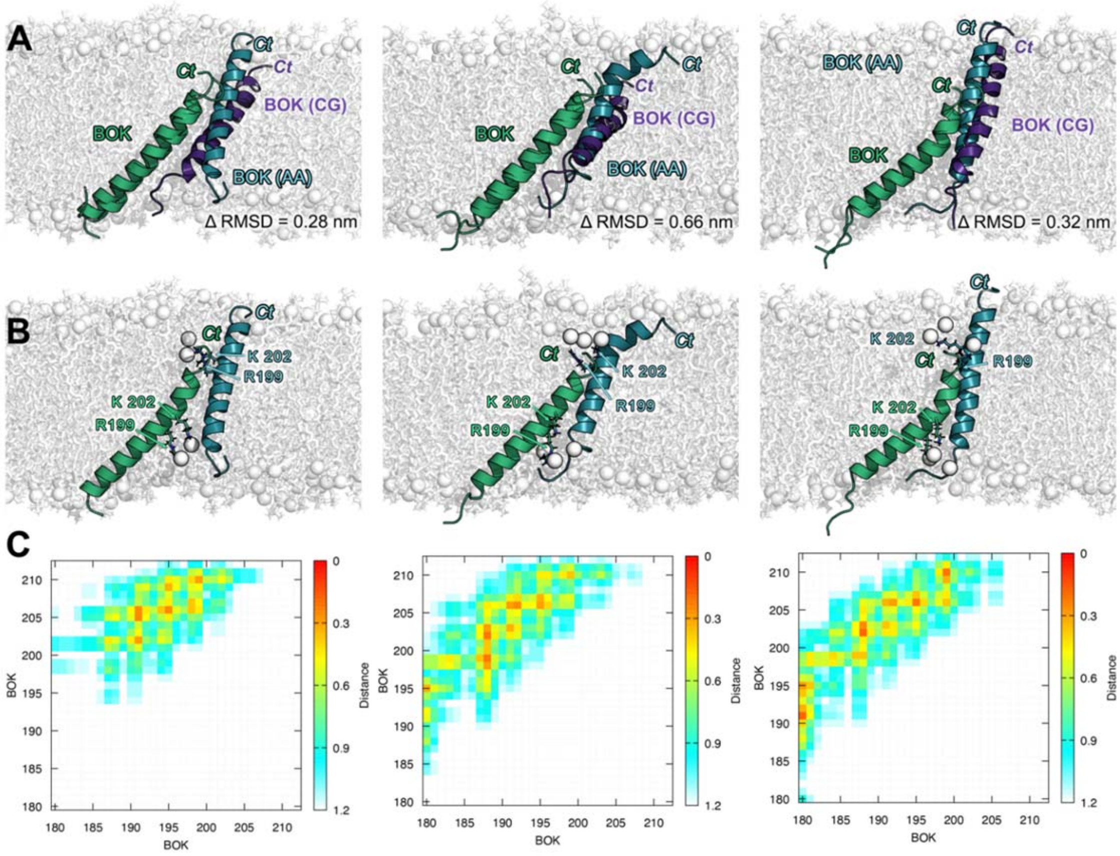


**Figure S5: Structures of three representative BOK/BOK-I homodimers.**

**A** - Orientation of the homodimers after 1 µs atomistic simulation visualized by alignment of one BOK-TMD (light green) and different coloring of the second BOK-TMD (original CG position in dark purple, after 1µs AA simulation in dark green). The deviation between the positions of the backbone atoms in the CG and AA structure is given as Δ RMSD.

**B** - Membrane indentation by R199 and K202 (shown as sticks). The phosphates of the deformed lipids are shown as nontransparent spheres. Unlike the others BOK/BOK homodimers, BOK/BOK-I deforms both membrane leaflets, because R199 and K202 of one BOK are anchored in the cytosolic membrane leaflet, while the R199 and K202 of the other BOK are anchored in the ER-facing leaflet

**C** - Contact maps of BOK-TMD in the corresponding dimers after 1µs AA simulations.

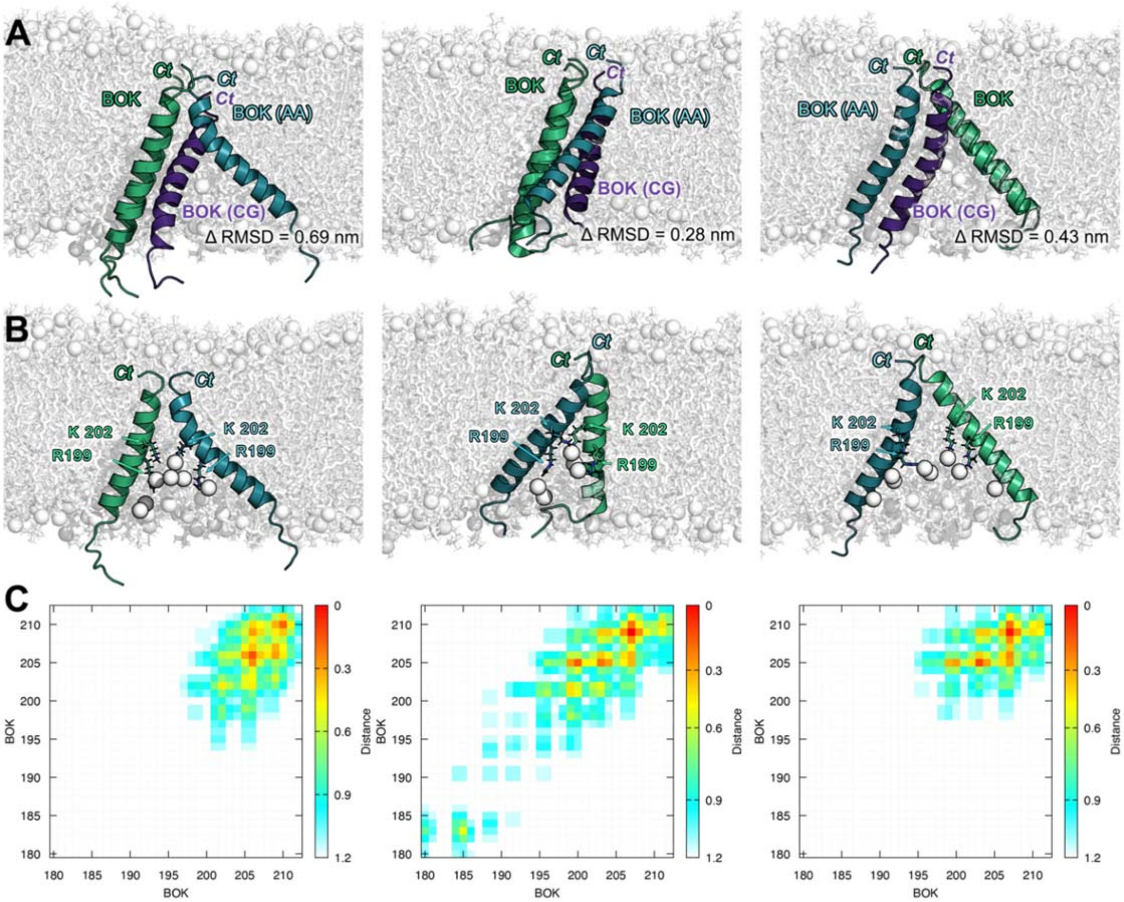


**Figure S6: Structures of three representative BOK-BOK-II homodimers.**

**A** - Orientation of the homodimers after 1μs atomistic simulation visualized by alignment of one BOK-TMD (light green) and different coloring of the second BOK-TMD (original CG position in dark purple, after 1μs AA simulation in dark green). The deviation between the positions of the backbone atoms in the CG and AA structure is given as Δ RMSD.

**B** - Membrane indentation by R199 and K202 (shown as sticks). The phosphates of the deformed lipids are shown as nontransparent spheres.

**C** - Contact maps of BOK-TMD in the corresponding dimers after 1μs AA simulations.

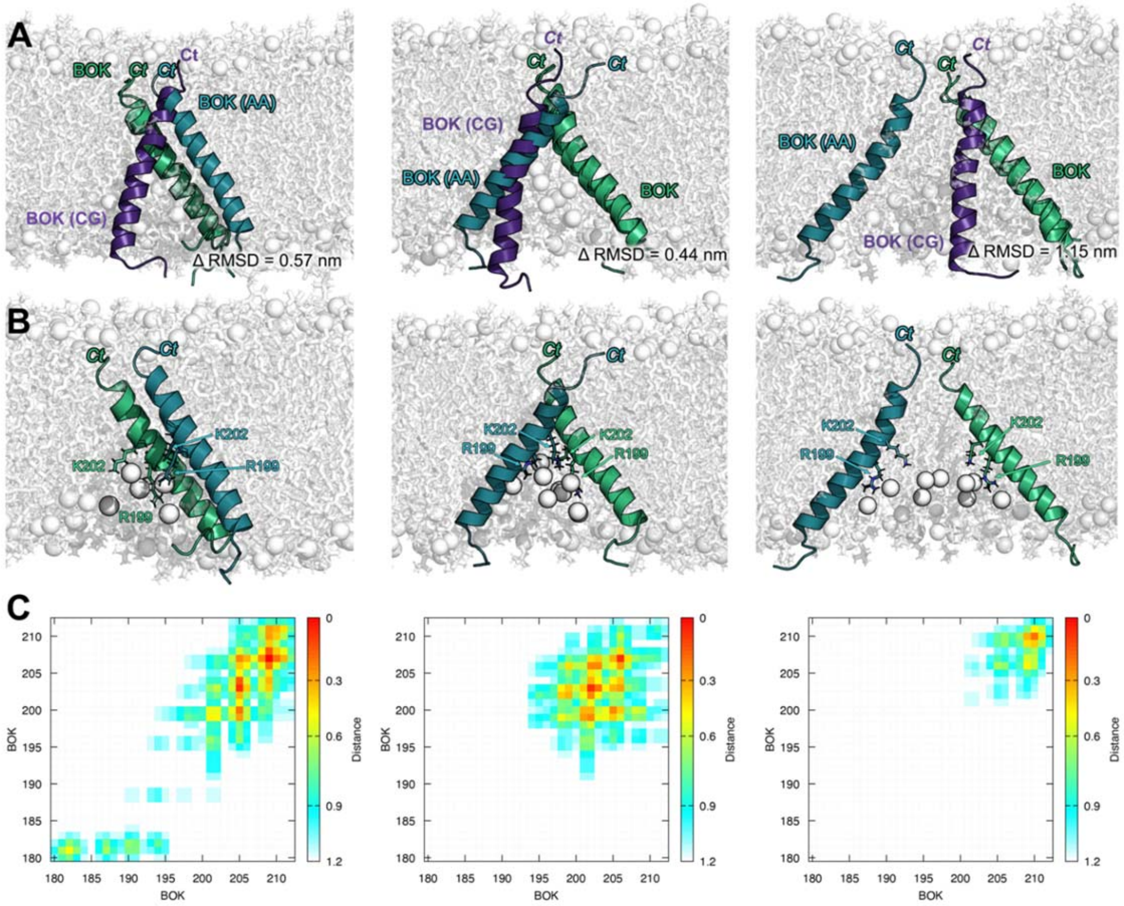


**Figure S7: Structures of three representative BOK-BOK-III homodimers.**

**A** - Orientation of the homodimers after 1µs atomistic simulation visualized by alignment of one BOK-TMD (light green) and different coloring of the second BOK-TMD (original CG position in dark purple, after 1µs AA simulation in dark green).

**B** - Membrane indentation by R199 and K202 (shown as sticks). The phosphates of the deformed lipids are shown as nontransparent spheres.

**C** - Contact maps of BOK-TMD in the corresponding dimers after 1µs AA simulations.

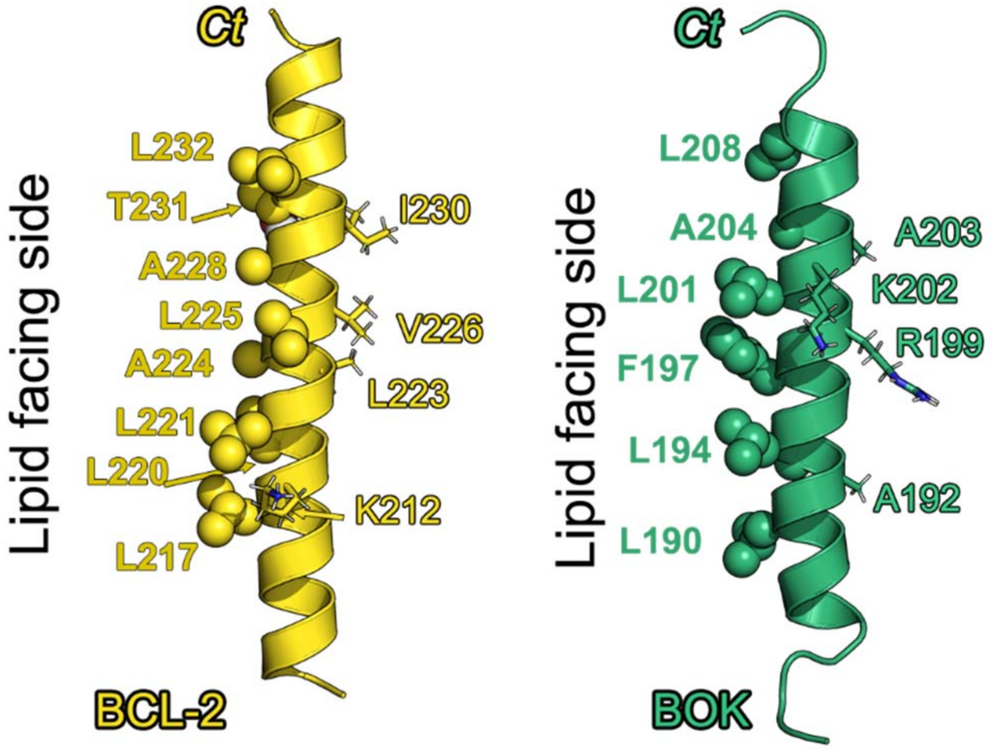


**Figure S8: BOK and BCL-2 TMD residues.**

Visualization of BCL-2-TMD and BOK-TMD helices with certain important residues highlighted as sticks (e.g. membrane anchors K212$^{BCL-2}$ and R199$^{BOK}$ and K202$^{BOK}$ or interaction interface residues I230$^{BCL-2}$ V226$^{BCL-2}$ L223$^{BCL-2,}$ A203$^{BOK}$ A192$^{BOK}$). The lipid facing residues are highlighted as spheres.

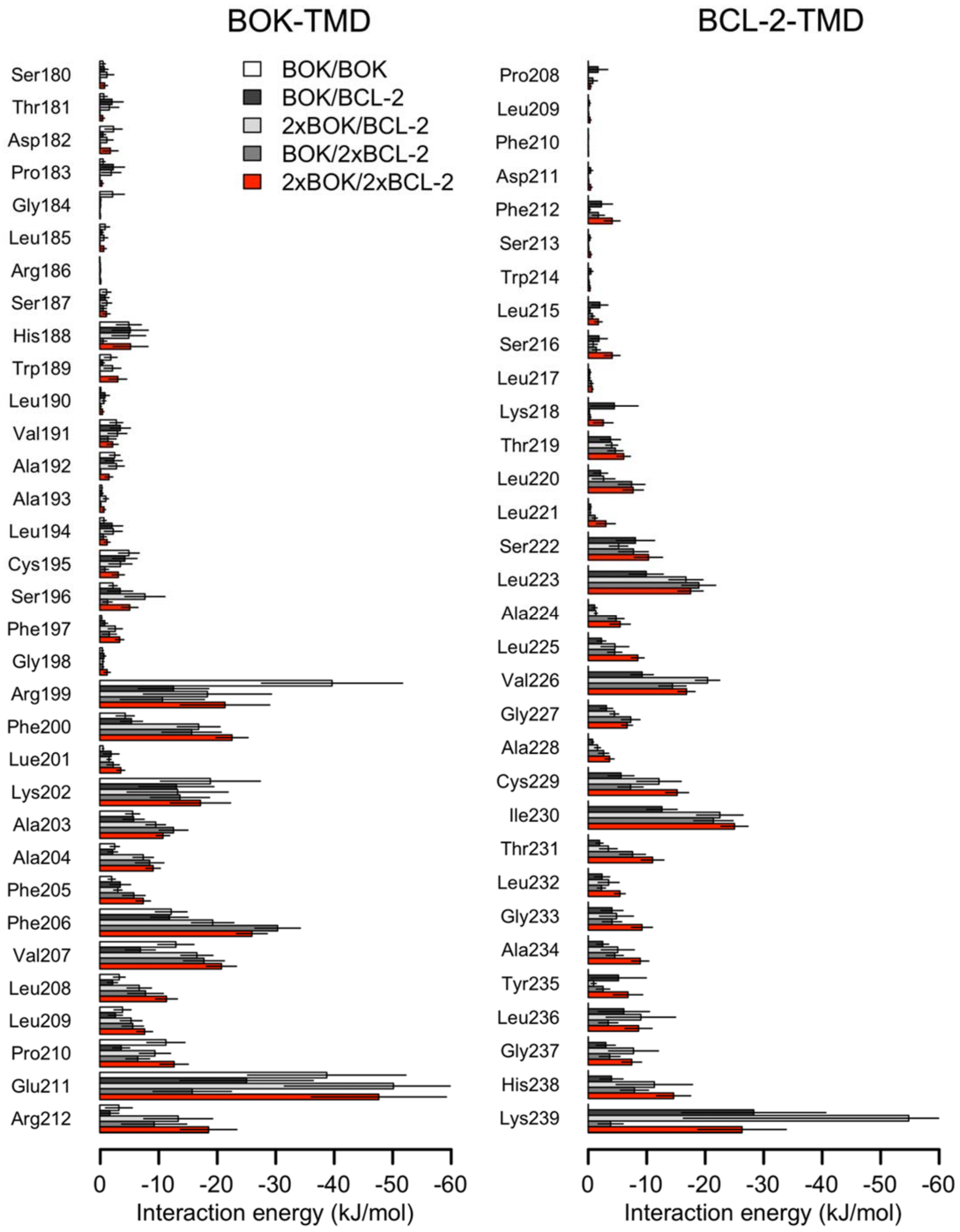


**Figure S9: interaction energies of BOK-TMD and BCL-2-TMD residues.**

Average interaction energies of each residue of BOK-TMD (left) or BCL-2-TMD (right) with all other peptides over the last 100 ns of AA simulations of dimers, trimer and tetramers. The error bars denote SEM over individual simulations. The outer error bars pf Glu211BOK in 2xBOK/BCL-2 trimer and Lys239BCL-2 2xBOK/BCL-2 trimer simulations are cut at -60 kJ/mol due to visualisation purposes.

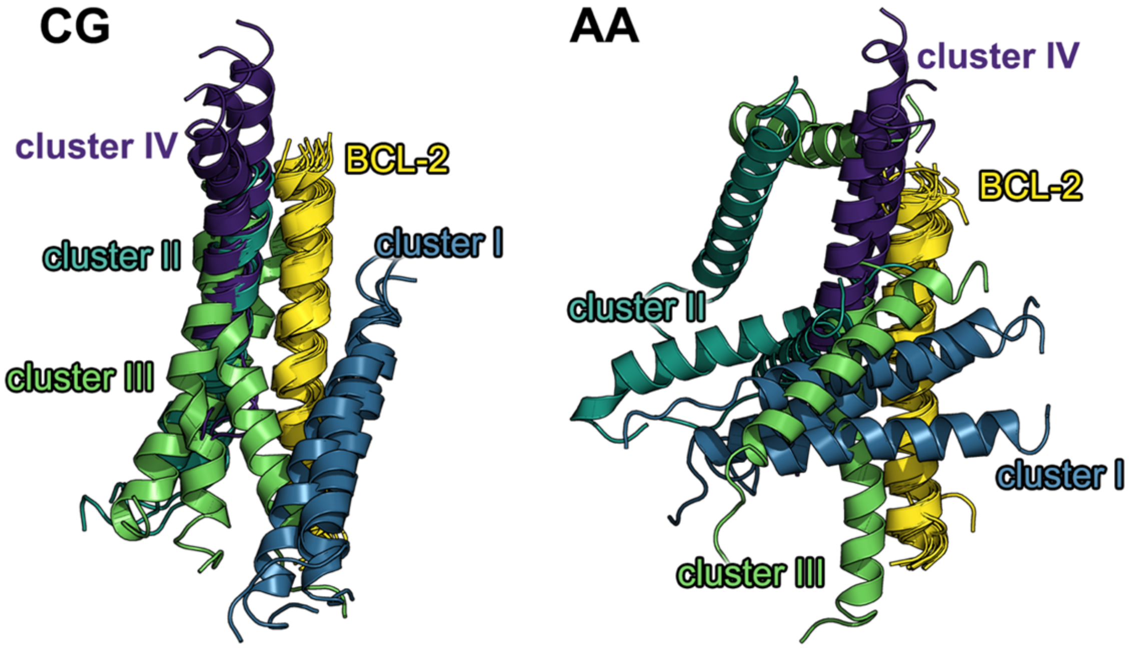


**Figure S10: BOK/BCL-2 heterodimer.**

Visualization of clusters of BOK/BCL-2 heterodimers after an overlay to an identical reference (BCL2 in yellow). CG labels structures prior to AA simulations, as AA structures after 1µs AA simulation are shown. The BOK TMDs in cluster I are colored blue, in cluster II aquamarine, in cluster III green and in cluster IV purple.

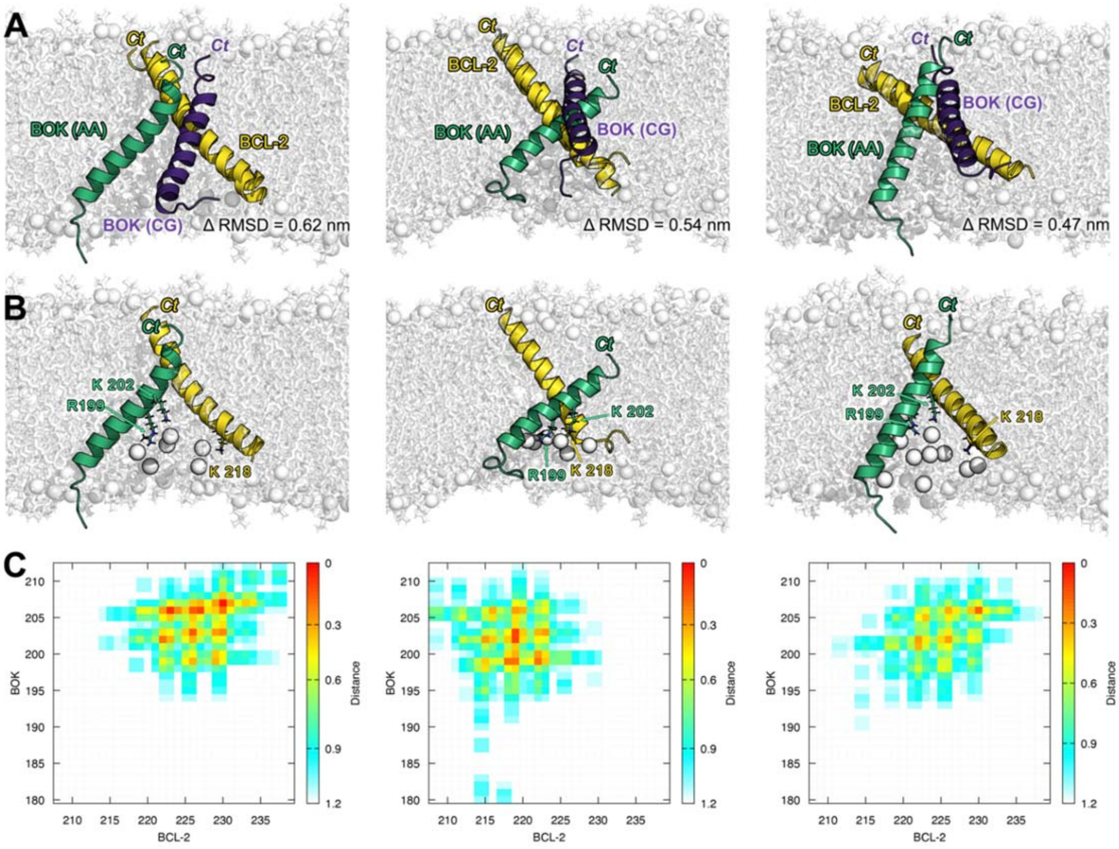


**Figure S11: Structures of three representative BOK/BCL-2-I heterodimers.**

**A** - Orientation of the heterodimers after 1µs atomistic simulation visualized by alignment of BCL-2-TMD (yellow) and different coloring of BOK-TMD (original CG position in dark purple, after 1µs AA simulation in green). The deviation between the positions of the backbone atoms in the CG and AA structure is given as Δ RMSD.

**B** - Membrane indentation by R199BOK and K202BOK as well as K218BCL-2 (shown as sticks). The phosphates of the deformed lipids are shown as nontransparent spheres.

**C** - Contact maps of the BOK/BCL-2 heterodimers after 1µs AA simulations.

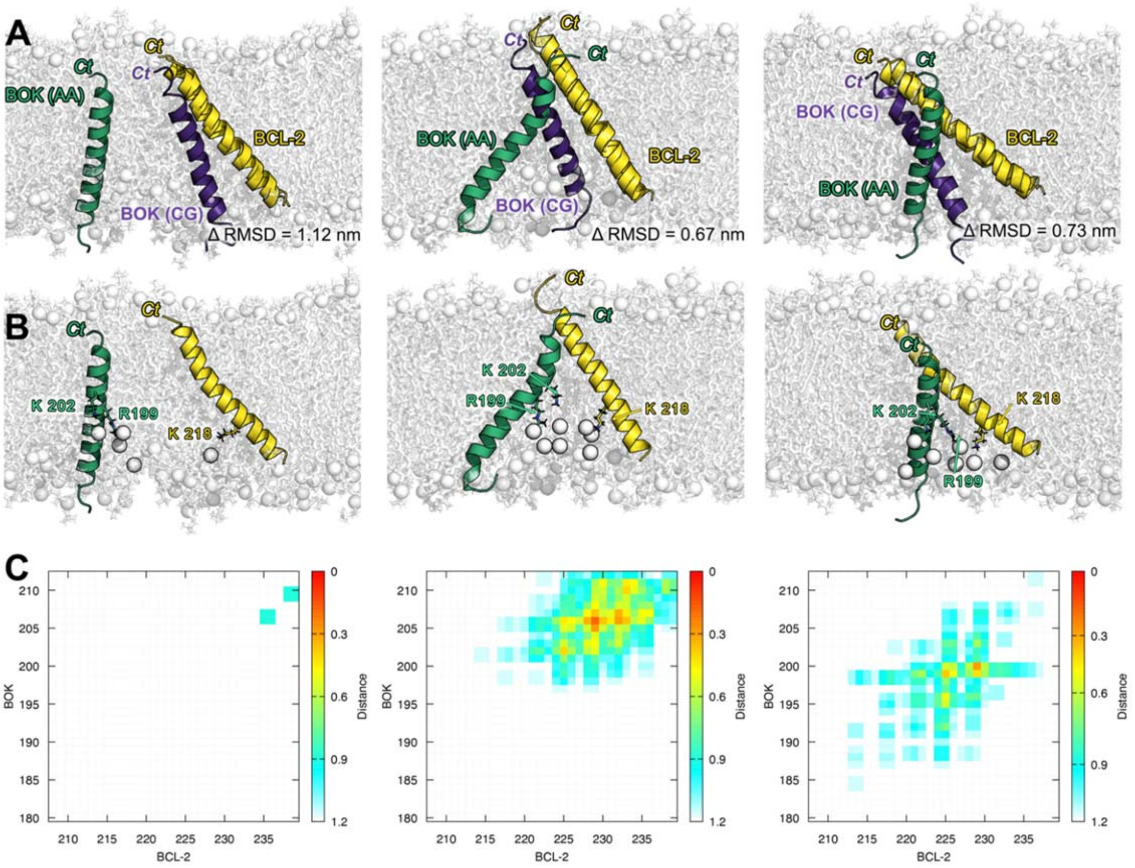


**Figure S12: Structures of three representative BOK/BCL-2-II heterodimers.**

**A** - Orientation of the heterodimers after 1µs atomistic simulation visualized by alignment of BCL-2-TMD (yellow) and different coloring of BOK-TMD (original CG position in dark purple, after 1µs AA simulation in green). The deviation between the positions of the backbone atoms in the CG and AA structure is given as Δ RMSD.

**B** - Membrane indentation by R199BOK and K202BOK as well as K218BCL-2 (shown as sticks). The phosphates of the deformed lipids are shown as nontransparent spheres.

**C** - Contact maps of the BOK/BCL-2 heterodimers after 1µs AA simulations.

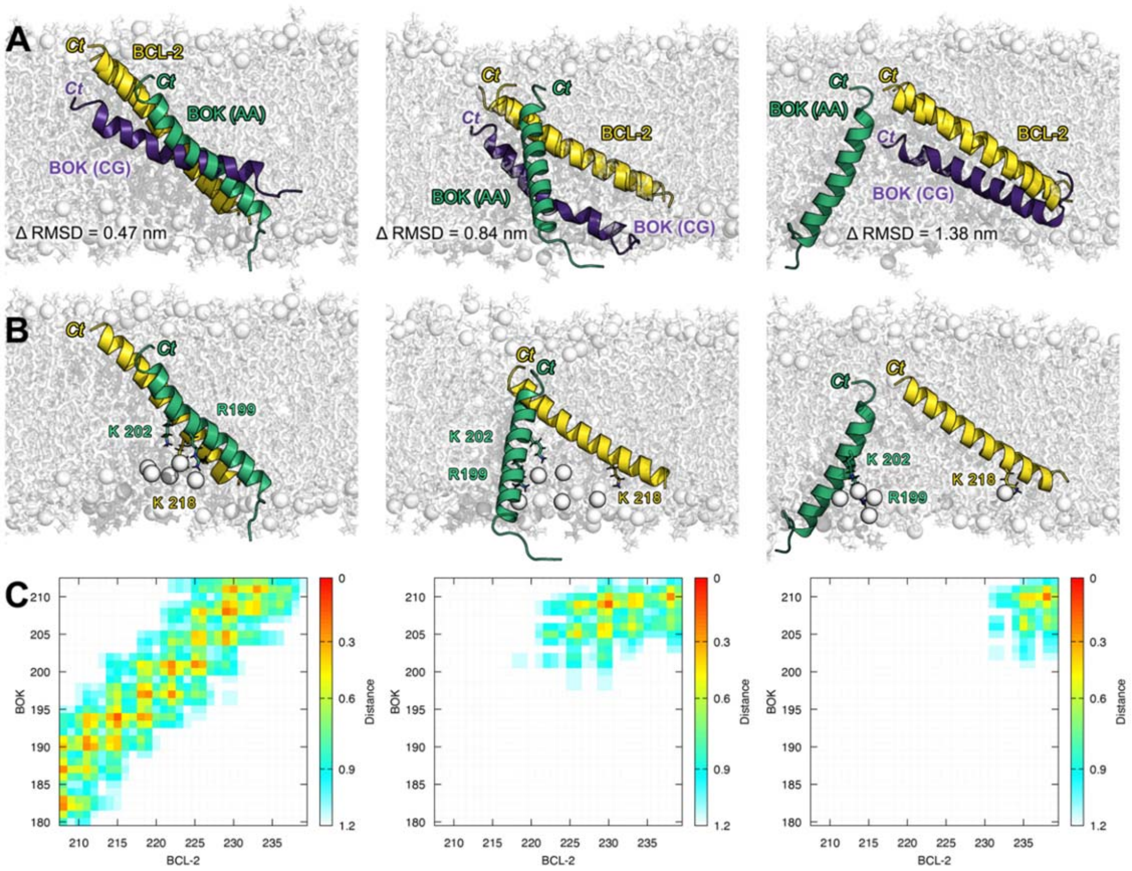


**Figure S13: Structures of three representative BOK/BCL-2-III heterodimers.**

**A** - Orientation of the heterodimers after 1µs atomistic simulation visualized by alignment of BCL-2-TMD (yellow) and different coloring of BOK-TMD (original CG position in dark purple, after 1µs AA simulation in green). The deviation between the positions of the backbone atoms in the CG and AA structure is given as Δ RMSD.

**B** - Membrane indentation by $R199^{BOK}$ and $K202^{BOK}$ as well as $K218^{BCL-2}$ (shown as sticks). The phosphates of the deformed lipids are shown as nontransparent spheres.

**C** - Contact maps of the BOK/BCL-2 heterodimers after 1µs AA simulations.

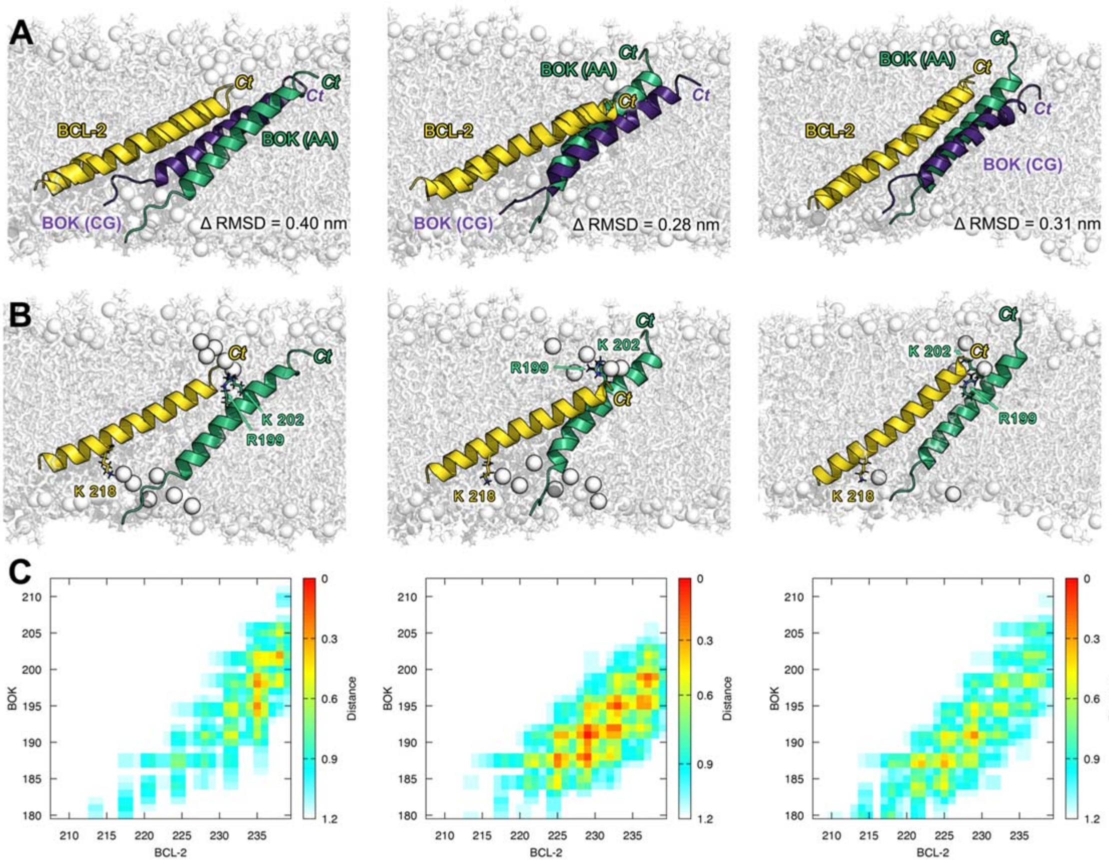


**Figure S14: Structures of three representative BOK/BCL-2-IV heterodimers.**

**A** - Orientation of the heterodimers after 1µs atomistic simulation visualized by alignment of BCL-2-TMD (yellow) and different coloring of BOK-TMD (original CG position in dark purple, after 1µs AA simulation in green). The deviation between the positions of the backbone atoms in the CG and AA structure is given as Δ RMSD.

**B** - Membrane indentation by $R199^{BOK}$ and $K202^{BOK}$ as well as $K218^{BCL\text{-}2}$ (shown as sticks). The phosphates of the deformed lipids are shown as nontransparent spheres.

**C** - Contact maps of the BOK/BCL-2 heterodimers after 1µs AA simulations.

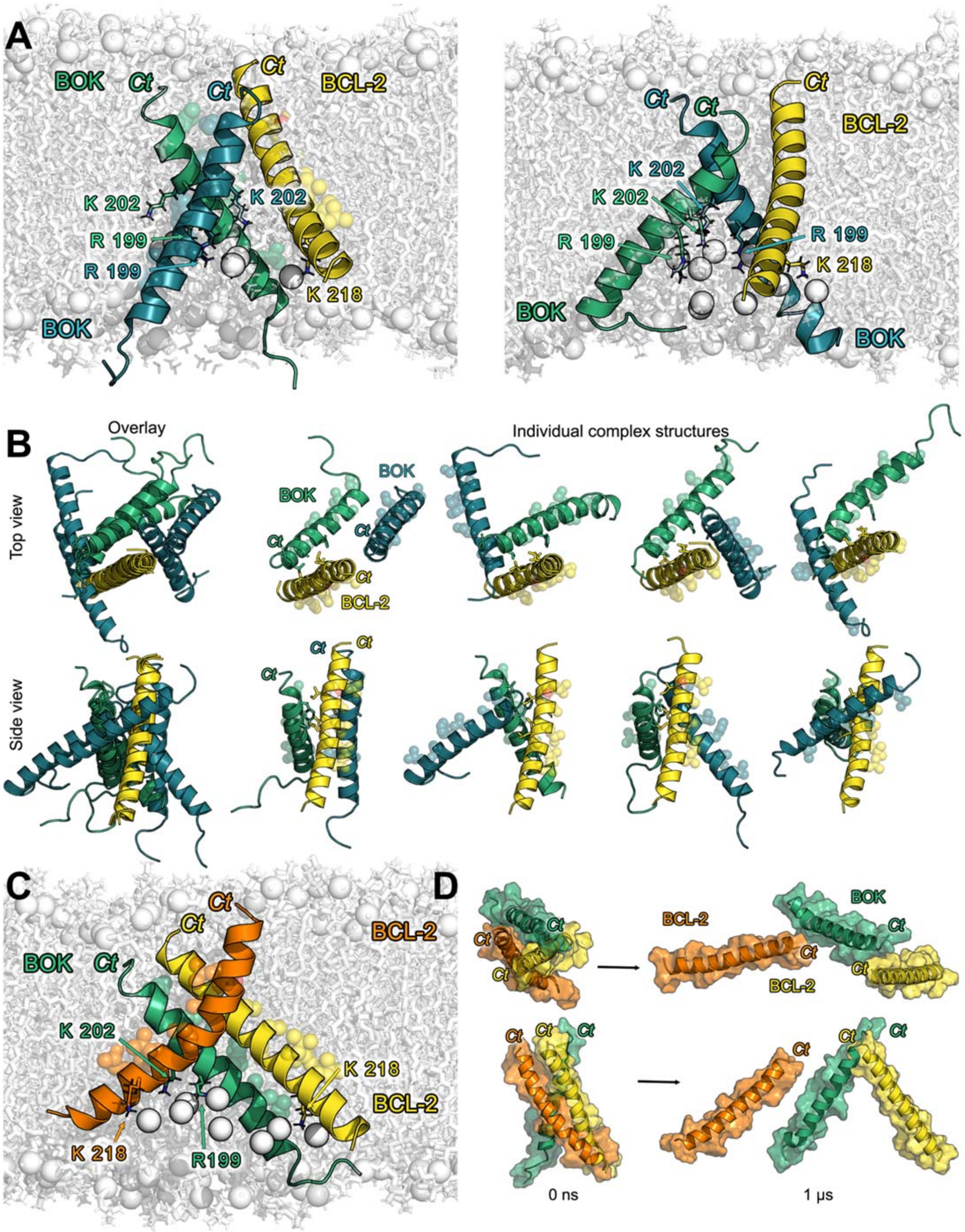
A
BOK
Ct
Ct
Ct
BCL-2
K 202
K 202
R 199
R 199
K 218
BOK
Ct
Ct
Ct
BCL-2
K 202
K 202
R 199
R 199
K 218
BOK
BOK
B
Overlay
Individual complex structures
Top view
BOK
BOK
Ct
Ct
Ct
BCL-2
Side view
Ct
Ct
Ct
C
Ct
Ct
BCL-2
BOK
Ct
K 202
K 218
BCL-2
K 218
R199
D
BOK
Ct
Ct
Ct
BCL-2
Ct
Ct
BCL-2
Ct
Ct
Ct
Ct
Ct
Ct
0 ns
1 µs

**Figure S15: Structures of BOK/2xBCL-2 heterotrimers.**

**A** - Membrane indentation caused by R199BOK, K202BOK and K218BCL-2, shown as sticks, in the two most common BOK/BOK/BCL-2 heterotrimers. The most often observed heterotrimer, comprising 58% of all BOK/BOK/BCL-2 heterotrimers formed is shown on the left. Here, BCL-2-TMD stabilizes the symmetric BOK-TMD/BOK-TMD dimer cluster II by L223BCL-2, V226BCL-2, C229BCL-2 and I230BCL-2. The two K202BOK residues are localized deeply in the membrane hydrophobic core, the amino groups are interacting with lipid carbonyls. The second most often (15%) observed BOK/BOK/BCL-2 heterotrimer, shown on the right, consists of BOK/BCL-2-TMD heterodimer cluster I ($A203^{BOK}$, $V207^{BOK}$ $F200^{BOK}$ interact with $L223^{BCL\text{-}2}$, $V226^{BCL\text{-}2}$, $C229^{BCL\text{-}2}$ and $I230^{BCL\text{-}2}$) and the second BOK-TMD attached in different positions (see also Figure S14). The phosphates of the deformed lipids are shown as non-transparent spheres. One BOK-TMD is colored green, another BOK-TMD is colored blue, BCL-2-TMD is colored yellow.

**B** - Structures of BOK/BOK/BCL-2 trimers with conserved BOK/BCL-2 cluster I heterodimer (BCL-2 colored yellow, BOK colored green). The putative other BOK peptide is shown in blue. On the left most, the overlay of all four trimer clusters is shown. The top row brings a view from the C-terminal side, the bottom row shows a side view.

**C** - Membrane indentation caused by $R199^{BOK}$, $K202^{BOK}$ and $K218^{BCL\text{-}2}$, shown as sticks, in the two most common BOK/BCL-2/BCL-2 heterotrimers for the most often observed heterotrimer, comprising 56% of all BOK/BCL-2/BCL-2 heterotrimers formed. Here, one BCL-2-TMD binds to BOK-TMD in the orientation of BOK/BCL-2-TMD heterodimer cluster I and the other BCL-2-TMD binds to BOK-TMD with the same interaction interface like in the BOK/BCL-2-TMD heterodimer cluster II. Additionally, the two BCL-2-TMDs form a symmetric interaction interface with I230 and T231 from both BCL-2-TMDs. The small alanine and glycine residues play again a crucial role for close contacts between the TMDs. $A204^{BOK}$ thus allows a close contact with one BCL-2-TMD and $A203^{BOK}$ with the other BCL-2-TMD. BOK-TMD is colored green, one BCL-2-TMD is yellow and the other BCL-2-TMD orange. The lipid phosphates in the indent are shown as non-transparent spheres.

**D** - Reorientation of a left-handed BOK/BCL-2/BCL-2 heterotrimer over the course of an atomistic simulation. Top row shows the view from the C-terminal side, bottom row a side view, with the C-termini on the top. The orange colored BCL-2-TMD dissociates completely from the BOK/BCL-2 heterotrimer, which converts from a compact left-handed to a V-shaped loosed dimer. BOK-TMD is colored green and the BCL-2-TMD which stays in the heterodimer is colored yellow. A surface representation of the peptides is included to highlight the compactness of the initial trimer and the full dissociation of one BCL-2-TMD from the two other peptides after 1µs.

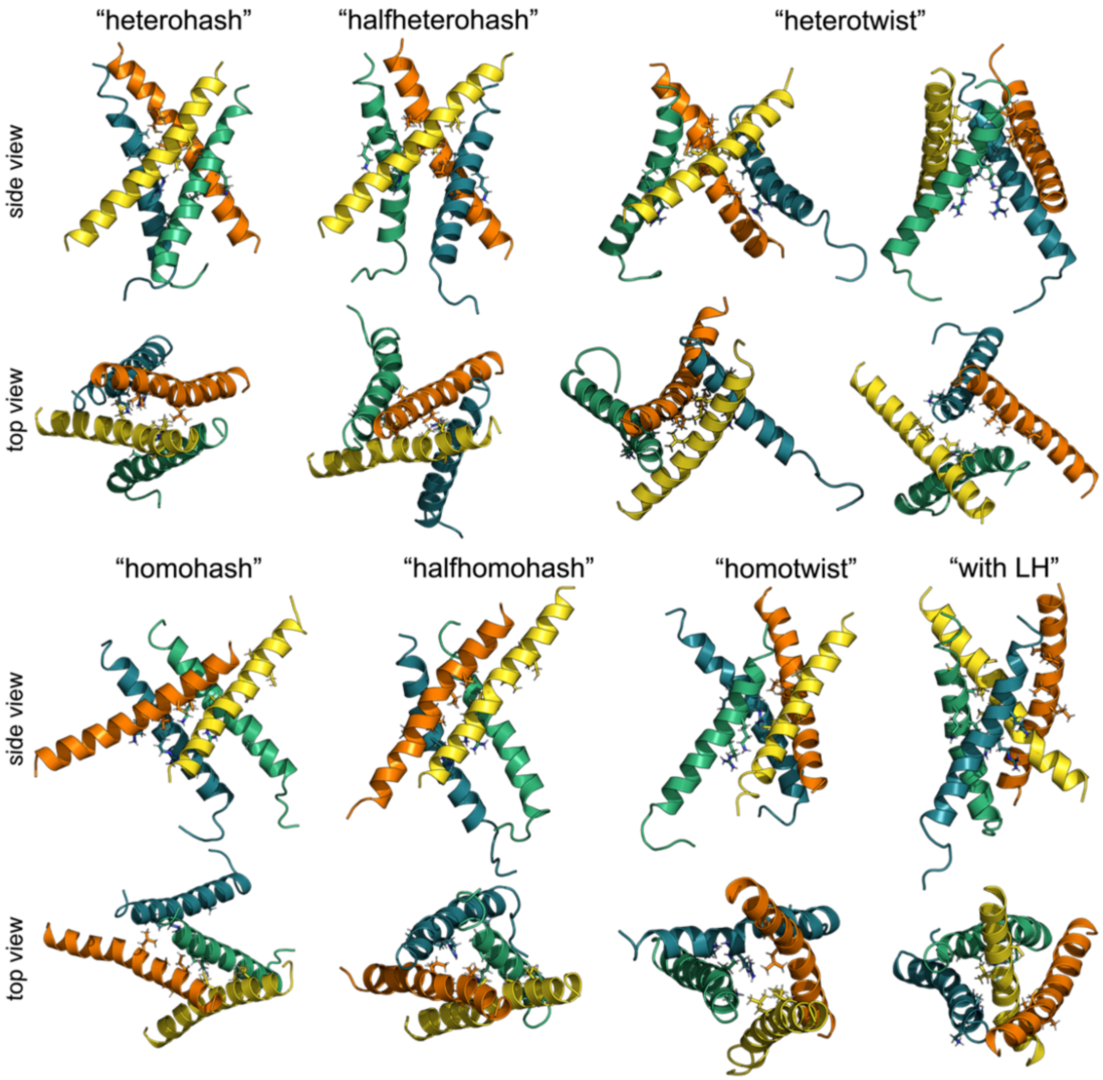


**Figure S16: Structures of representative heterotetramers.**

Representative heterotetramer structures which formed by spontaneous association of two BOK-TMD (colored green and teal) with two BCL-2-TMDs (colored yellow and orange) over 50 µs of CG simulations. The C-termini are located on the top in the side view and in the front in the top view. $R199^{BOK}$, $K202^{BOK}$, $I230^{BCL\text{-}2}$, $C229^{BCL\text{-}2}$, $V226^{BCL\text{-}2}$, and $L223^{BCL\text{-}2}$ are shown as sticks. "Hash" heterodimers consist of two parallel dimers, "halfhash" tetramers are made of one parallel and one crossed dimer, in the "twist" tetramers each peptide crosses its neighbour forming right-handed compact tetramers. In the structure labelled as "with LH" a compact trimer with lefhandedly attached monomer (here the orange BCL-2-TMD) is shown. After 18 AA simulations, 67 % of all heterodimers had compact "twist" shapes, 17% were homohash, 6% halfhomohash and in two cases (11%) linear dimer of dimers was observed.

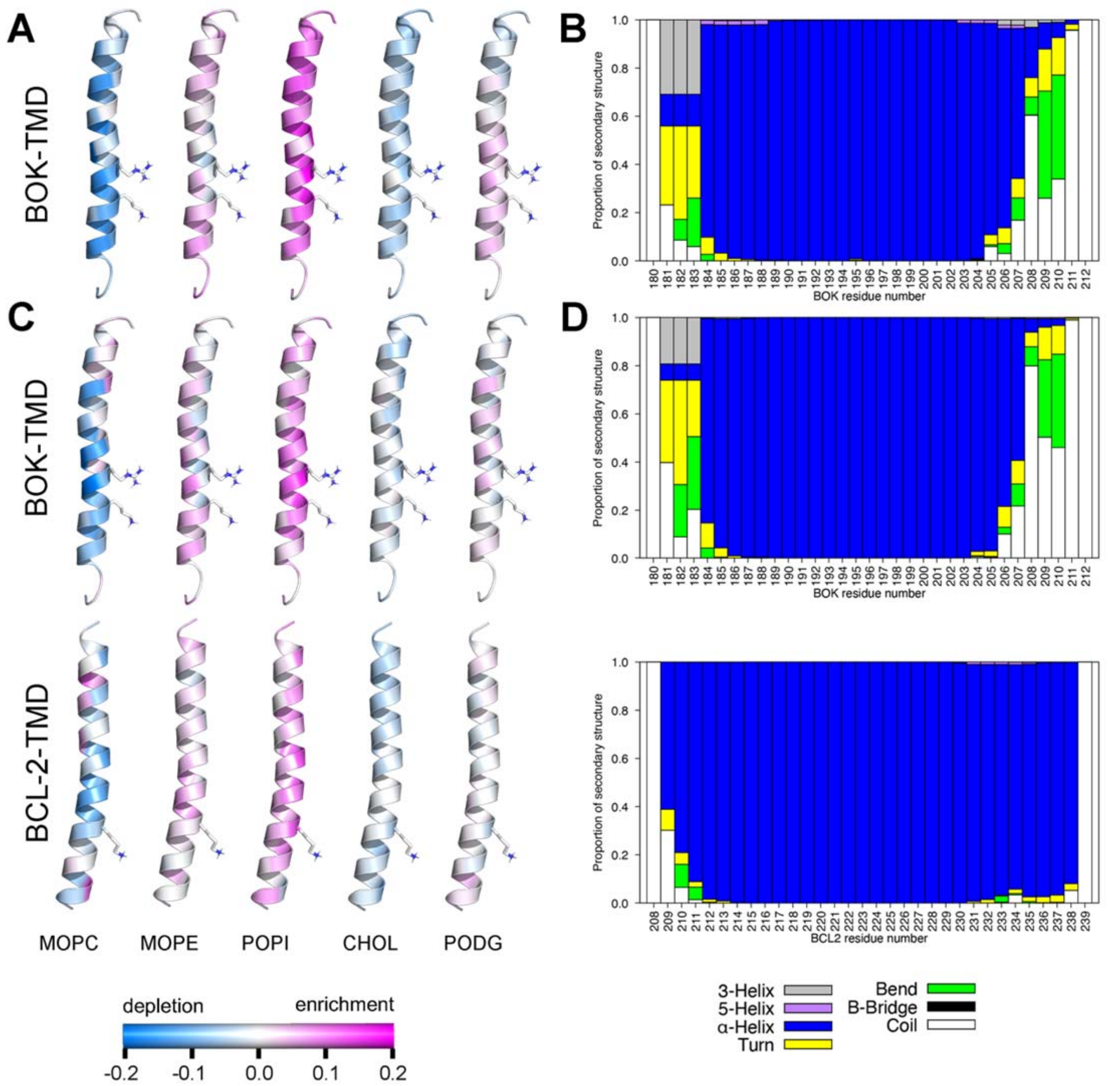


**Figure S17: Lipid-TMD interaction.**

Lipid enrichment or depletion around BOK-TMD or BCL-2-TMD and the secondary structure of the peptides. (A) Lipid enrichment/depletion in BOK-TMD/BOK-TMD homodimers. (B) Average BOK-TMD secondary structure per residue in BOK-TMD/BOK-TMD homodimers. (C) Lipid enrichment/depletion in BOK-TMD/BCL-TMD heterodimers and (D) average secondary structure per residue of BOK-TMD and BCL-2-TM2 in BOK-TMD/BCL-2/TMD heterodimers. The membrane indenting positively charged R199BOK, K202BOK, and K218BCL-2 are in A and C visualized as sticks.

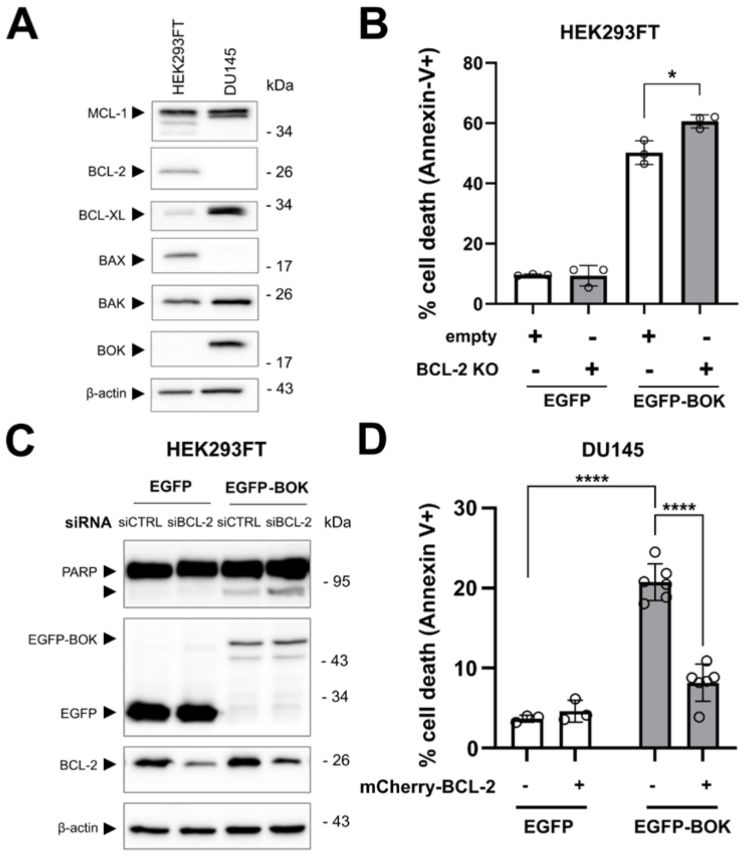
A
HEK293FT
DU145
kDa
MCL-1
- 34
BCL-2
- 26
BCL-XL
- 34
BAX
- 17
BAK
- 26
BOK
- 17
β-actin
- 43
B
HEK293FT
% cell death (Annexin-V+)
80
60
40
20
0
*
empty
BCL-2 KO
EGFP
EGFP-BOK
C
HEK293FT
EGFP
EGFP-BOK
siRNA
siCTRL
siBCL-2
siCTRL
siBCL-2
kDa
PARP
- 95
EGFP-BOK
- 43
- 34
EGFP
BCL-2
- 26
β-actin
- 43
D
DU145
% cell death (Annexin V+)
30
20
10
0
****
****
mCherry-BCL-2
EGFP
EGFP-BOK

**Figure S18: Functional relevance of BOK-TMD and BCL-2-TMD interaction in apoptosis.**

**A** - Detection of various Bcl-2 family proteins in whole cell lysates of HEK293FT and DU145 cells using Western Blot. β-actin was used as loading control. One independent experiment.

**B** - HEK293FT cells were co-transfected with plasmids for the expression of a CRISPR/Cas9 vector targeting BCL2 and EGFP or EGFP-BOK (empty = empty vector backbone control). After 18 h, cells were harvested, and cell death was assessed using Annexin-V-APC staining and flow cytometry. EGFP+/Annexin-V+ percentage of cells is shown as mean ± sd from three independent experiments.

**C** - HEK293FT cells were transfected with siRNA for BCL-2 (siBCL-2) or control siRNA (siCTRL) for 24 h and were subsequently transfected with plasmids for the expression of EGFP-BOK or EGFP. After 18 h, cells were harvested and analyzed by Western Blot. PARP, EGFP(-BOK) and BCL-2 expression were detected. β-actin was used as loading control. Representative blot from two independent experiments.

**D** - DU145 cells were transfected with plasmids for the expression of EGFP-BOK or EGFP in combination with mCherry-BCL-2 or an empty vector as a control. After 42 h, cells were stained with Annexin-V-APC and cell death (EGFP+/Annexin-V+ cells) was assessed using flow cytometry. Mean ± sd from n = 3 (EGFP) or n = 6 (EGFP-BOK) independent experiments.

**Table S1A: overview of protein sequences used for cloning.**

| Name | Function | Uniprot entry |
|---|---|---|
| BAX | Pro-apoptotic | Q07812 |
| BAK | Pro-apoptotic | Q16611 |
| BOK | Pro-apoptotic | Q9UMX3 |
| BCL-2 | Anti-apoptotic | P10415 |
| BCL-XL | Anti-apoptotic | Q07817 |
| MCL-1 | Anti-apoptotic | Q07820 |
| TOM5 | Mito control | Q8N4H5 |
| cb5 | ER control | P00167 |

**Table S1B: C-terminal TMD sequences used for plasmid cloning.**

| protein | function | amino acid sequence |
|---|---|---|
| BAX | Pro-apoptotic | $_{166}$GTPTWQTVTI FVAGVLTASL TIWKKMG$_{192}$ |
| BAK | Pro-apoptotic | $_{184}$GNGPILNVLV VLGVVLLGQF VVRRFFKS$_{211}$ |
| BOK | Pro-apoptotic | $_{185}$LRSHWLVAAL CSFGRFLKAA FFVLLPER$_{212}$ |
| BCL-2 | Anti-apoptotic | $_{208}$PLFDFSWLSL KTLLSLALVG ACITLGAYLG HK$_{239}$ |
| BCL-XL | Anti-apoptotic | $_{209}$RFNRWFLTGM TVAGVVLLGS LFSRK$_{233}$ |
| MCL-1 | Anti-apoptotic | $_{328}$LEGGIRNVLL AFAGVAGVGA GLAYLIR$_{350}$ |
| TOM5 | Mito control | $_{23}$DVISSIRNFL IYVALLRVTP FILKKLDSI$_{51}$ |
| cb5 | ER control | $_{100}$DTTIDSSSSW WTNWVIPAIS AVAVALMYRL YMAED$_{134}$ |

**Table S2: Primer sequences used for Gibson assembly.**

Gibson assembly mediated generation of plasmids for the expression of NanoBiT-TMD fusion proteins. Plasmid backbone pBiT_1.1-C[TK/LgBiT] was digested using restriction enzymes HindIII and XbaI. fw = forward; rv = reverse

| fragment | Orientation (adjacent fragment) | primer sequence |
|---|---|---|
| mCitrine/ mTurquoise2 | fw (backbone) | 5′-CCTGCAGCGACCCGCTTAAAGATATCATGGTGAGCAAGGGCGAG-3′ |
| | rv (T2A) | 5′-AGCCGAATTCCTTGTACAGCTCGTCCATGC-3′ |
| T2A | fw (mCitr/mTurq2) | 5′-GCTGTACAAGGAATTCGGCTCCGGCGAGGGCAGAGGAAGTC-3′ |
| | rv (LgBiT) | 5′-TGAAGACCATTGCGCAAGGGCCGGGATTCTCCTCC-3′ |
| LgBiT | fw (T2A) | 5′-CCCTTGCGCAATGGTCTTCACACTCGAAG-3′ |
| | rv (TMD) | 5′-AAGCTTGCTGCCGCCGCCGCCGCTGCCGCCGCCGCCACTGTTGATGGTTACTCG-3′ |
| BAX-TMD | fw (LgBiT/SmBiT) | 5′-GGAGGTGGAGGCTCAGGAGGTGGAGGCTCAAAGCTTGGGACGCCCACGTGGCAG-3′ |
| | rv (backbone) | 5′-GCGGCCGGCCGCCCCGACTCTCGAGTCAGCCCATCTTCTTCCAGATGGTG-3′ |
| BAK-TMD | fw (LgBiT/SmBiT) | 5′-CTCAGGAGGTGGAGGCTCAAAGCTTGGCAACGGCCCCATCCTG-3′ |
| | rv (backbone) | 5′-GCGGCCGGCCGCCCCGACTCTCGAGTCATGATTTGAAGAATCTTCGTACC-3′ |
| BOK-TMD | fw (LgBiT/SmBiT) | 5′-CTCAGGAGGTGGAGGCTCAAAGCTTCTGAGGAGCCACTGGCTG-3′ |
| | rv (backbone) | 5′-GCGGCCGGCCGCCCCGACTCTCGAGTCACCTCTCGGGCAGCAG-3′ |
| BCL-2-TMD | fw (LgBiT/SmBiT) | 5′-CAGCGGCGGCGGCGGCAGCAAGCTTCCTCTGTTTGATTTCTCC-3′ |
| | rv (backbone) | 5′-GCGGCCGGCCGCCCCGACTCTCGAGTCACTTGTGGCCCAGATAG-3′ |
| BCL-XL-TMD | fw (LgBiT/SmBiT) | 5′-CAGCGGCGGCGGCGGCAGCAAGCTTCGCTTCAACCGCTGGTTC-3′ |
| | rv (backbone) | 5′-GCGGCCGGCCGCCCCGACTCTCGAGTCATTTCCGACTGAAGAGTGAG-3′ |
| MCL-1-TMD | fw (LgBiT/SmBiT) | 5′-CAGCGGCGGCGGCGGCAGCAAGCTTCTAGAAGGTGGCATCAGG-3′ |
| | rv (backbone) | 5′-GCGGCCGGCCGCCCCGACTCTCGAGTCATCTTATTAGATATGCCAAACC-3′ |
| TOM5-TMD | fw (LgBiT/SmBiT) | 5′-CAGCGGCGGCGGCGGCAGCAAGCTTGACGTGATCAGCAGCATC-3′ |
| | rv (backbone) | 5′-GCGGCCGGCCGCCCCGACTCTCGAGTCAGATGCTGTCCAGCTTC-3′ |

**Table S3: Primer sequences used for Gibson assembly.**

Gibson assembly–mediated generation of plasmids for the expression of mCitrine or mTurquoise2-tagged Bcl-2 TMD fusion proteins. Plasmid backbone pEGFP-C1 (Clontech) was digested previously to NEBuilder cloning using restriction enzymes XhoI and AgeI.

| **fragment** | **Orientation (adjacent fragment)** | **primer sequence** |
|---|---|---|
| mTurquoise2 | fw (backbone) | 5′-CGTCAGATCCGCTAGCGCTAGCCACCATGGTGAGCAAGGGCGAG-3′ |
| | rv (Linker) | 5′-GCAGAATTCGAAGCTTGAGCCTCGAGTCAGCCCATCTTC-3′ |
| Linker + TMD | fw (fluo) | 5′-CATGGACGAGCTGTACAAGGAATTCGGAGGTGGAGGCTCAGGAG-3′ |
| BAX-TMD | rv (backbone) | 5′-GGAGAGGGGCACCGGTTCAGCCCATCTTCTTCCAG-3′ |
| BAK-TMD | rv (backbone) | 5′-GGAGAGGGGCACCGGTTCATGATTTGAAGAATCTTCGTACC-3′ |
| BOK-TMD | rv (backbone) | 5′-GGAGAGGGGCACCGGTTCACCTCTCGGGCAGCAG-3′ |
| BCL-2-TMD | rv (backbone) | 5′-GGAGAGGGGCACCGGTTCACTTGTGGCCCAGATAG-3′ |
| BCL-XL-TMD | rv (backbone) | 5′-GGAGAGGGGCACCGGTTCACTTCCTGCTGAACAG-3′ |
| MCL-1-TMD | rv (backbone) | 5′-GGAGAGGGGCACCGGTTCATCTTATTAGATATGCCAAACC-3′ |
| TOM5-TMD | rv (backbone) | 5′-GGAGAGGGGCACCGGTTCAGATGCTGTCCAGCTTC-3′ |

**Table S4: Primer sequences used for Gibson assembly.**

Gibson assembly-mediated generation of plasmids for the expression of chimeric fluorophore-fused full length Bcl-2 proteins with exchanged TMDs. Plasmids used were described previously (Einsele-Scholz et al. 2016).

| fragment | Orientation (adjacent fragment) | primer sequence |
|---|---|---|
| BOK Core | rv (BAX-TMD) | 5′-TGGGCGTCCCGCCAGGGTCTGTGCTGACC-3′ |
| BCL-2 Core | rv (cb5-TMD) | 5′-TAGTAGTGTCCCGCATGCTGGGGCCGTA-3′ |
| | rv (TOM5-TMD) | 5′-TGATCACGTCCCGCATGCTGGGGCCGTA-3′ |
| BAX-TMD | fw (BOK Core) | 5′-AGACCCTGGCGGGACGCCCACGTGGCAG-3′ |
| | rv (backbone) | 5′-GGAGAGGGGCACCGGTTCAGCCCATCTTCTTCCAG-3′ |
| Cb5-TMD | fw (BCL-2 Core) | 5′-CAGCATGCGGGACACTACTATTGATTCTAGTTC-3′ |
| | rv (backbone) | 5′-CGGCCGCCACTGTGCTGGATGAATTCTCAATCTTCAGCCATGTAC-3′ |
| TOM5-TMD | fw (BCL-2 Core) | 5′-CAGCATGCGGGACGTGATCAGCAGCATC-3′ |
| | rv (backbone) | 5′-CGGCCGCCACTGTGCTGGATGAATTCTCAGATGCTGTCCAGCTTC-3′ |

**Table S5: Summary of performed simulations.**

Heterotrimer structures for AA simulations were taken from 2xBOK-TMD/2xBCL-2-TMD simulations in which heterotrimers formed.

| System | resolution | Number of sims | Simulation length (μs) |
|---|---|---|---|
| **BOK-TMD/BOK-TMD** | CG | 50 | 10 |
| | AA | 13 | 1 |
| **BOK-TMD/BCL-2-TMD** | CG | 50 | 10 |
| | AA | 15 | 1 |
| **2xBOK-TMD/2xBCL-2-TMD** | CG | 50 | 50 |
| | AA | 18 | 1 |
| **2xBOK-TMD/BCL-2-TMD** | AA | 9 | 1 |
| **BOK-TMD/2xBCL-2-TMD** | AA | 10 | 1 |

**Table S6: BOK/BOK homodimers.**

Summary and characteristics of BOK/BOK dimers spontaneously formed in MD simulations. The errors denote standard errors of the mean over 3 representative structures for each cluster. Characteristics of CG structures were obtained at atomistic resolution after conversion by backward, i.e. at 0 ns of AA simulations. AA characteristics were taken after 1μs of AA simulation. RH stands for right-handed.

| BOK/BOK Cluster | Handed-ness | Frequency CG [%] | ΔRMSD AA/CG [nm] | COM distance CG [nm] | COM distance AA [nm] | Longitudinal shift CG [nm] | Longitudinal shift AA [nm] |
|---|---|---|---|---|---|---|---|
| BOK/BOK-I | RH | 55 | 0.43±0.10 | 1.92±0.17 | 2.11±0.04 | 1.56±0.19 | 1.81±0.06 |
| BOK/BOK-II | RH | 13 | 0.46±0.15 | 1.37±0.13 | 1.95±0.38 | 0.35±0.11 | 0.68±0.26 |
| BOK/BOK-III | RH | 11 | 0.72±0.27 | 1.43±0.06 | 2.01±0.75 | 0.39±0.03 | 0.86±0.54 |

**Table S7: BOK/BCL-2 heterodimers.**

Summary and characteristics of observed BOK/BCL-2 heterodimers. The errors denote standard errors of the mean over 3 representative structures for each cluster. Characteristics of CG structures were obtained at atomistic resolution after conversion by backward, i.e. at 0 ns of AA simulations. AA characteristcs were taken after 1μs of AA simulation.

| BOK/BCL-2 Cluster | Handed-ness | Frequency CG [%] | ΔRMSD (AA/CG) [nm] | COM distance CG [nm] | COM distance AA [nm] | Longitudinal shift CG [nm] | Longitudinal shift AA [nm] |
|---|---|---|---|---|---|---|---|
| BOK/BCL-2-I | RH | 44 | 0.54±0.05 | 1.24±0.05 | 1.29±0.13 | 1.29±0.02 | 0.94±0.24 |
| BOK/BCL-2-II | RH | 34 | 0.84±0.18 | 1.04±0.05 | 2.06±0.69 | 0.43±0.03 | 0.91±0.41 |
| BOK/BCL-2-III | LH | 8 | 0.90±0.32 | 0.97±0.12 | 2.15±0.75 | 0.60±0.11 | 1.57±0.42 |
| BOK/BCL-2-IV | RH | 7 | 0.33±0.04 | 2.24±0.02 | 2.24±0.09 | -1.57±0.03 | -1.29±0.07 |

**Table S8: Relative lipid enrichment/depletion in BOK-TMD (heterotetramers).**

Relative lipid enrichment (positive values) or depletion (negative values) per BOK-TMD residue in 2xBOK-TMD/2xBCL-2-TMD heterotetramer simulations.

| BOK | chol | MOPC | MOPE | PODG | POPI |
|---|---:|---:|---:|---:|---:|
| Ser180 | -6.9% | 2.8% | 0.1% | -3.1% | 7.0% |
| Thr181 | -6.7% | 3.4% | -0.2% | -3.6% | 7.1% |
| Asp182 | -6.7% | -1.0% | 4.0% | -3.0% | 6.7% |
| Pro183 | -6.0% | -1.3% | -2.7% | -1.9% | 12.0% |
| Gly184 | -6.9% | -3.8% | 0.3% | -3.8% | 14.2% |
| Leu185 | -5.4% | -3.8% | -0.7% | -1.4% | 11.3% |
| Arg186 | -3.7% | -10.9% | 2.5% | -2.5% | 14.5% |
| Ser187 | -5.9% | -2.9% | 0.7% | -1.4% | 9.5% |
| His188 | -6.9% | 2.3% | 0.0% | -2.0% | 6.6% |
| Trp189 | -2.7% | -11.9% | 1.6% | 0.9% | 12.0% |
| Leu190 | -4.0% | -11.0% | 2.9% | 0.8% | 11.3% |
| Val191 | -5.3% | 2.4% | 0.1% | -0.6% | 3.4% |
| Ala192 | -4.5% | 1.6% | -3.5% | -0.8% | 7.1% |
| Ala193 | -3.5% | -4.8% | -1.5% | 1.4% | 8.4% |
| Leu194 | -3.4% | -9.6% | 0.5% | 2.7% | 9.8% |
| Cys195 | -5.9% | 0.5% | -4.0% | -1.4% | 10.9% |
| Ser196 | -1.9% | -7.4% | -4.6% | -0.3% | 14.1% |
| Phe197 | -1.2% | -18.0% | -0.1% | 5.0% | 14.3% |
| Gly198 | -4.4% | -4.3% | -4.9% | 2.4% | 11.2% |
| Arg199 | -6.3% | -7.4% | -5.4% | -2.2% | 21.4% |
| Phe200 | -2.9% | -5.5% | -3.6% | 0.2% | 11.9% |
| Lue201 | -2.8% | -9.4% | -2.9% | 3.4% | 11.6% |
| Lys202 | -3.9% | -9.6% | -5.9% | 1.8% | 17.5% |
| Ala203 | -6.3% | 1.3% | -3.8% | -4.0% | 12.7% |
| Ala204 | -5.8% | 9.6% | -5.3% | -0.7% | 2.1% |
| Phe205 | -3.6% | -11.5% | -1.4% | 3.0% | 13.6% |
| Phe206 | -5.1% | 2.5% | -1.1% | -0.5% | 4.1% |
| Val207 | -4.7% | 7.7% | -1.6% | -2.6% | 1.3% |
| Leu208 | -5.2% | 1.0% | -2.1% | 0.0% | 6.2% |
| Leu209 | -4.7% | -3.8% | 1.2% | 1.1% | 6.2% |
| Pro210 | -3.8% | 4.0% | 1.7% | -0.6% | -1.2% |
| Glu211 | -5.9% | 5.0% | 5.5% | -3.3% | -1.3% |
| Arg212 | -5.4% | -4.2% | 1.1% | -0.5% | 9.0% |

**Table S9: Relative lipid enrichment/depletion in BCL-2-TMD (heterotetramers).**

Relative lipid enrichment (positive values) or depletion (negative values) per BCL-2-TMD residue in 2xBOK-TMD/2xBCL-2-TMD heterotetramer simulations.

| BCL-2 | chol | MOPC | MOPE | PODG | POPI |
|---|---:|---:|---:|---:|---:|
| Pro208 | -4.6% | -2.0% | 1.1% | -1.9% | 7.3% |
| Leu209 | -2.6% | -10.7% | 2.9% | 1.4% | 8.9% |
| Phe210 | -1.8% | -9.8% | 5.7% | 0.7% | 5.1% |
| Asp211 | -6.9% | -0.4% | 6.7% | -2.1% | 2.7% |
| Phe212 | -3.5% | -4.0% | -1.2% | 0.4% | 8.4% |
| Ser213 | -2.5% | -5.1% | 1.7% | 2.3% | 3.7% |
| Trp214 | -4.6% | -6.1% | 0.8% | 0.4% | 9.5% |
| Leu215 | -6.5% | -2.1% | -4.4% | -3.1% | 16.1% |
| Ser216 | -3.3% | -1.4% | -1.3% | 1.2% | 4.8% |
| Leu217 | -3.3% | -11.3% | 3.5% | 3.5% | 7.6% |
| Lys218 | -4.9% | -3.9% | -4.6% | -2.6% | 16.0% |
| Thr219 | -5.3% | -7.6% | -4.5% | -2.8% | 20.2% |
| Leu220 | -1.7% | -9.8% | 2.0% | 2.5% | 7.0% |
| Leu221 | -2.1% | -12.6% | -0.2% | 1.5% | 13.5% |
| Ser222 | -2.1% | -9.1% | -6.1% | -3.4% | 20.7% |
| Leu223 | -5.5% | 4.0% | -3.0% | -1.9% | 6.3% |
| Ala224 | -3.4% | 0.3% | -0.8% | -1.0% | 4.9% |
| Leu225 | -0.4% | -13.2% | -0.6% | -1.0% | 15.3% |
| Val226 | -3.7% | -4.4% | -3.5% | -3.1% | 14.8% |
| Gly227 | -5.9% | 17.6% | -3.8% | -2.6% | -5.3% |
| Ala228 | -3.5% | -1.3% | -1.0% | -1.2% | 7.0% |
| Cys229 | -3.5% | -1.1% | 0.5% | -1.3% | 5.3% |
| Ile230 | -6.0% | 10.2% | -0.6% | -2.4% | -1.2% |
| Thr231 | -3.5% | 8.2% | -2.4% | -2.6% | 0.2% |
| Leu232 | -2.9% | -8.0% | 1.6% | 0.5% | 8.8% |
| Gly233 | -5.2% | 1.5% | 2.2% | -1.6% | 3.1% |
| Ala234 | -6.5% | 15.6% | -2.6% | -3.1% | -3.5% |
| Tyr235 | -4.1% | -3.1% | 1.3% | -0.8% | 6.7% |
| Leu236 | -3.3% | -8.1% | 2.5% | 0.7% | 8.2% |
| Gly237 | -5.8% | 5.0% | -3.0% | -1.4% | 5.1% |
| His238 | -6.5% | 10.9% | -3.4% | -3.4% | 2.4% |
| Lys239 | -5.9% | -4.8% | 4.1% | -1.3% | 7.9% |

### SI References


Berendsen HJC, Postma JPM, van Gunsteren WF, DiNola A, Haak JR (1984) Molecular dynamics with coupling to an external bath. *The Journal of Chemical Physics.* 8: 3684–3690.

Darden T, York D, Pedersen L (1993) Particle mesh Ewald: An N ·log(N ) method for Ewald sums in large systems. *The Journal of Chemical Physics.* 12: 10089–10092.

Einsele-Scholz S, Malmsheimer S, Bertram K, Stehle D, Johänning J, Manz M, Daniel PT, Gillissen BF, Schulze-Osthoff K, Essmann F (2016) Bok is a genuine multi-BH-domain protein that triggers apoptosis in the absence of Bax and Bak. *Journal of cell science.* 15: 3054.

Evans DJ&Holian BL (1985) The Nose–Hoover thermostat. *The Journal of Chemical Physics.* 8: 4069–4074.

Hess B (2008) P-LINCS: A Parallel Linear Constraint Solver for Molecular Simulation. *Journal of chemical theory and computation.* 1: 116–122.

Irving JA, Whisstock JC, Lesk AM (2001) Protein structural alignments and functional genomics. *Proteins.* 3: 378–382.

Kabsch W&Sander C (1983) Dictionary of protein secondary structure: pattern recognition of hydrogen-bonded and geometrical features. *Biopolymers.* 12: 2577–2637.

Páll S&Hess B (2013) A flexible algorithm for calculating pair interactions on SIMD architectures. *Computer Physics Communications.* 12: 2641–2650.

Parrinello M&Rahman A (1980) Crystal Structure and Pair Potentials: A Molecular-Dynamics Study. *Phys. Rev. Lett.* 14: 1196–1199.

Parrinello M&Rahman A (1981) Polymorphic transitions in single crystals: A new molecular dynamics method. *Journal of Applied Physics.* 12: 7182–7190.

Prasad R, Sliwa-Gonzalez A, Barral Y (2020) Mapping bilayer thickness in the ER membrane. *Science advances.* 46:

R Core Team (2022) R: A language and environment for statistical computing. *R Foundation for Statistical Computing, Vienna, Austria.*

Schrödinger L (2023) The PyMOL Molecular Graphics System, Version 2.0. *pymol.org*.

Scrima S, Tiberti M, Campo A, Corcelle-Termeau E, Judith D, Foged MM, Clemmensen KKB, Tooze SA, Jäättelä M, Maeda K et al. (2022) Unraveling membrane properties at the organelle-level with LipidDyn. *Computational and Structural Biotechnology Journal*: 3604–3614.

Wassenaar TA, Ingólfsson HI, Böckmann RA, Tieleman DP, Marrink SJ (2015) Computational Lipidomics with insane: A Versatile Tool for Generating Custom Membranes for Molecular Simulations. *Journal of chemical theory and computation.* 5: 2144–2155.

Williams T&Kelley C (2013) Gnuplot 4.6: an interactive plotting program. *gnuplot.sourceforge.net*.
